# Universal 3D Motif dynamics in RNA: The A-minor Switch

**DOI:** 10.64898/2026.05.08.718354

**Authors:** Christian Steinmetzger, Hampus Karlsson, Carolina Fontana, Natalia Galindo Riera, Magdalena Riad, David Kosek, Judith Schlagnitweit, Emilie Steiner, Sarah Friebe Sandoz, Emma R. Andersson, Katja Petzold

## Abstract

A-minor motifs consist of adenosines docking into adjacent RNA minor grooves, are the most prevalent 3D interaction stabilizing RNA structures, and widely considered as being static. NMR spectroscopy reveals a secondary structure equilibrium of these motifs between engaged and disengaged states, which we term the A-minor switch. A switch in *E.coli* ribosome helix 44 consists of a sparsely populated, transient single-nucleotide register shift that sequesters adenosines from their 3D structural A-minor contacts. Mutational trapping of the NMR-defined, A-minor-engaged ground and-disengaged excited state, combined with cryo-electron microscopy, visualizes this dynamic switch mechanism. Additionally, A-minor switches were identified using secondary structure ensemble analysis and mutational trapping was found to impair bacterial growth, directly linking RNA dynamics and function. Because A-minor motifs are widespread in structured RNAs, these findings establish A-minor switches as a general regulatory layer between secondary structure dynamics and tertiary contacts, exposing a new therapeutic target class.

## Results and Discussion

A-minor motifs, in which the sugar edge of adenosine docks into the minor groove of an adjacent Watson–Crick base pair, are both highly abundant and conserved long-range tertiary contacts in structured RNA (**Figure 1**).^1–3^ They organize the tertiary architecture of riboswitches and catalytic RNAs such as ribozymes or RNase P; in the ribosome, hundreds of A-minor contacts stitch together the rRNA core (REF 10.1038/86221, 10.1016/j.sbi.2006.04.002).^4,5^ Like hydrogen-bond networks in folded proteins, A-minor contacts have been understood as static architectural anchors that are formed once during folding and persist throughout function. Unlike in proteins however, the energy landscape of RNA folding is dominated by secondary structure,^6^ and larger tertiary contacts can be reshaped by changes to secondary structure features, i.e. base pairing, of few nucleotides.^1,7^ This raises a question: are A-minor motifs formed as a static 3D interaction upon initial RNA folding, or do they have an inherent propensity for switching between engaged and disengaged states with functional consequences?

**Figure 1.**
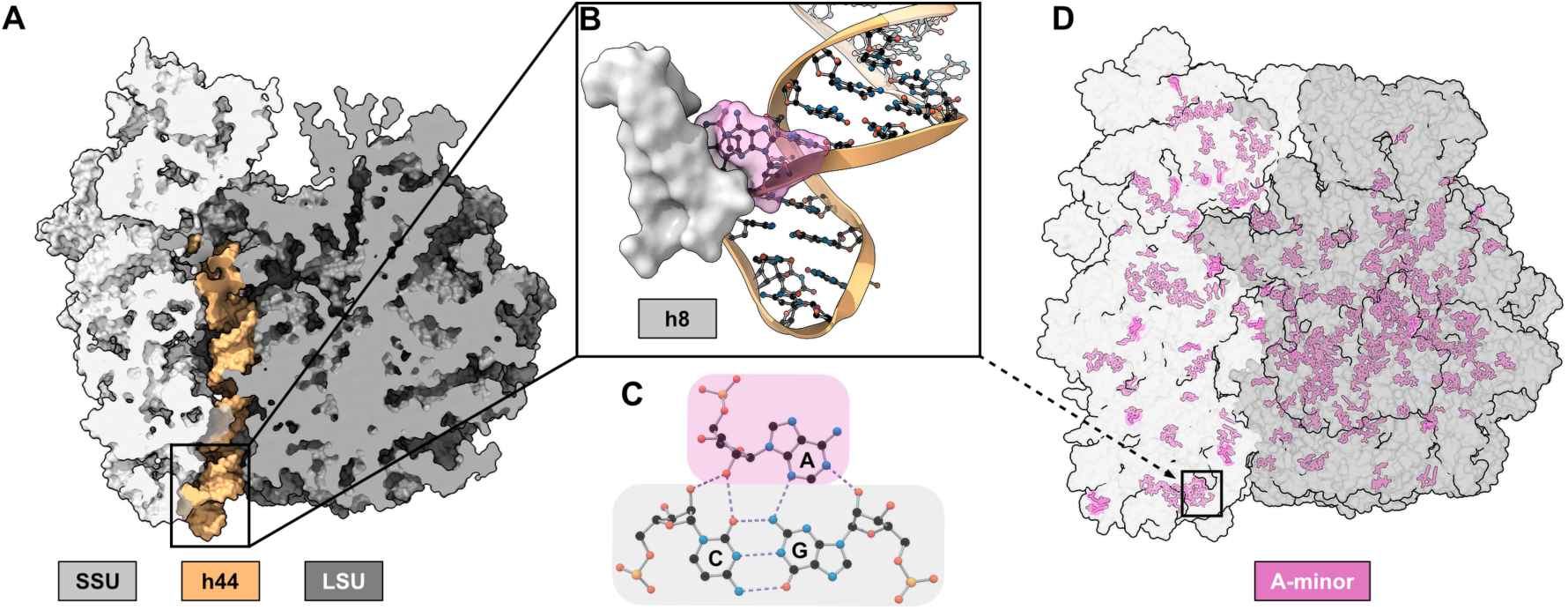
A-minor motifs in the bacterial ribosome. (**A**) A slice through the surface representation of the *E. coli* 70S ribosome (PDB 7K00) showing the 30S and 50S subunits (light and dark grey, respectively). Helix 44 (h44, orange) of the 16S rRNA spans the body of the SSU and forms bridging interactions to the 50S subunit. The apical stem-loop, h44-top, is highlighted as the focal point of this study. (**B**) Magnified view of the adenosine-rich bulge (pink) at the apex of h44 that is anchored to the 16S rRNA helix 8 (h8, light grey) through an A-minor contact (pink). (**C**) General structure of the A-minor interaction, in which the sugar face of adenosine (pink) is hydrogen-bonded with the minor groove of a Watson-Crick base pair^1,2^ (grey). (**D**) Distribution of all 276 unique adenosines engaged in A-minor motifs across the *E. coli* 70S ribosome (PDB 7K00). The h44–h8 interaction from panel (B) is highlighted. The high occurrence and spread of A-minor sites illustrates the prevalence of this tertiary structural motif, motivating closer elucidation of potential dynamics.

RNA function often involves structural exchange between a highly populated ground state (GS) and a transiently formed excited state (ES) that encode alternative functional and regulatory modes.^8^ Such dynamics are largely invisible to the methods used in high-resolution structure elucidation, e.g. X-ray crystallography, cryo-electron microscopy (cryo-EM) and conventional NMR spectroscopy: they report on GS conformations, but ESs typically evade direct structural characterization due to their low populations and lifetimes. For these structural equilibria on the microsecond–millisecond timescale, *R*_1ρ_ NMR relaxation dispersion (RD) experiments enable detection of such hidden states and provide chemical shift fingerprints that report on their structural features.^9–14^ *R*_1ρ_ is therefore an ideal tool for addressing the question of A-minor switching as a general phenomenon.

We chose the *E. coli* ribosome as a test system and catalogued a comprehensive set of 276 A-minor contacts in this large, structurally resolved RNA (**Figure 1D**). It is accessible to high-resolution structural and functional readouts, and the adenosine-rich bulge at the decoding-center A-site is already known to undergo transient flipping during codon recognition.^15–18^ We focused initially on the apical stem-loop of 16S rRNA helix 44 (h44-top), an A-minor contact between two conserved adenosines (A1446, A1447) and helix 8 (h8) (**Figure 1A to C**). Helix 44 is essential for subunit association,^19^ tRNA selection, translocation,^20^ and ribosome recycling,^21^ and constitutes the main binding site for aminoglycoside antibiotics at its basal A-site.^22^ Because of high sequence conservation between bacterial and mitochondrial A-sites, aminoglycosides exhibit poor target selectivity, often leading to severe side effects such as irreversible hearing loss.^23,24^ However, differences in the dynamic conformational landscapes of bacterial and eukaryotic rRNA have led to improved discrimination with certain antibiotics,^25,26^ making this site simultaneously a stringent test of whether reversible A-minor switching exists, and a context in which any switching would have direct functional and therapeutic consequences.

Here, we investigated the A-minor motif of the apical stem-loop of h44 by *R*_1ρ_ RD NMR and identified a transient single-nucleotide register shift within the native UUCG tetraloop that sequesters A1446 and A1447 from their A-minor interaction with SSU helix 8 (h8) (**Figure 1B**). This mechanism is structurally reminiscent of, but functionally distinct from, the dynamic behavior of A1492 and A1493 at the A-site,^17^ where these adenosines transiently flip out to monitor the codon–anticodon pairing.^18^ Analysis of predicted secondary structure ensembles around other ribosomal A-minor motifs reveals that reversible adenosine sequestration represents a recurring dynamic motif. Examining the biological consequences of trapping such A-minor switches in full ribosomes using cryo-EM and bacterial viability assays revealed functional impairment in ribosomes trapped in A-minor-sequestered states. Together, these findings demonstrate that A-minor interactions are not static architectural anchors but can be encoded as reversible structural equilibria within RNA.

### A register shift transiently disrupts the h44 A-minor contact

To investigate whether the adenosines involved in the apical A-minor interaction of h44 encode intrinsic conformational dynamics, we designed a 25 nt RNA hairpin (h44-top^WT^) comprising nucleotides 1442–1460 of wild-type (WT) *E. coli* 16S rRNA. A closing G:C base pair was added to facilitate in vitro transcription with T7 RNA polymerase^27^ and stabilize the basal stem. Chemical shift assignments for ^1^H, ^13^C and ^15^N resonances were obtained from a combination of 2D ^1^H–^1^H NOESY, ^1^H–^13^C HSQC, ^1^H–^15^N HSQC and HCN spectra measured on unlabeled and uniformly ^13^C,^15^N-labeled samples, respectively (**Figure S1**).^28^ The secondary structure of h44-top^WT^ agrees with the expected fold, with A1446, A1447 and A1456 forming a central bulge. Weak or absent imino NOE contacts for the flanking U1445:G1457 and C1448:G1455 base pairs indicate that this bulge region is highly dynamic, whereas the basal stem is stably formed. The apical stem-loop adopts a canonical cUUCGg tetraloop conformation with U1451 and C1452 in a C2’-endo ribose pucker (**Figure S1**). This is consistent with the full *E. coli* ribosome structure, in which A1446 and A1447 engage in an A-minor interaction with h8, while A1456 remains unpaired.^29^

To probe conformational exchange of h44-top^WT^ on the µs–ms time scale, we performed *R*_1ρ_ relaxation dispersion experiments.^9–11^ Elevated *R*_1ρ_ relaxation rates indicative of conformational exchange were observed across the central bulge and stem-loop (U1444–G1459) (**Figure 2A**, **B**, **Figure S2**, **Table S2**). Global fitting of the ^13^C and ^15^N relaxation dispersion profiles for the stem-loop region (C1449–G1455) yielded a well-defined excited state with a population (*p*_b_) of 2.74 ± 0.02% and exchange rate (*k*_ex_) of 1024 ± 8 s^−1^ (**Table S3**), consistent with a concerted structural rearrangement of the loop. In addition to this dominant excited state, all of the twelve dynamic nucleotides exhibited evidence for a second minor excited state (**Figure S2**, **Table S2**), indicating site-specific exchange processes superimposed on the global transition. The observed chemical shift changes associated with the major excited state (**Figure 2A**, **B**) are consistent with a single-nucleotide register shift within the apical stem-loop. In this rearranged state, U1450 is released from its wobble base pair with G1453 and incorporated into a cUUCg triloop, one of the more frequently occurring loop motifs in *E. coli* rRNA.^30^ Concomitantly, G1453 and G1454 form a WC base pair with C1449 and C1448, respectively, extending the upper stem. In this conformation, A1446 and A1447 adopt an intrahelical mismatch with A1456 and G1455, respectively, in the extended helix. In the context of the 16S rRNA, such a register shift would sequester A1446 and A1447 from their A-minor contact with h8 and thereby transiently disrupt this tertiary interaction, effectively switching off the A-minor motif.

**Figure 2.**
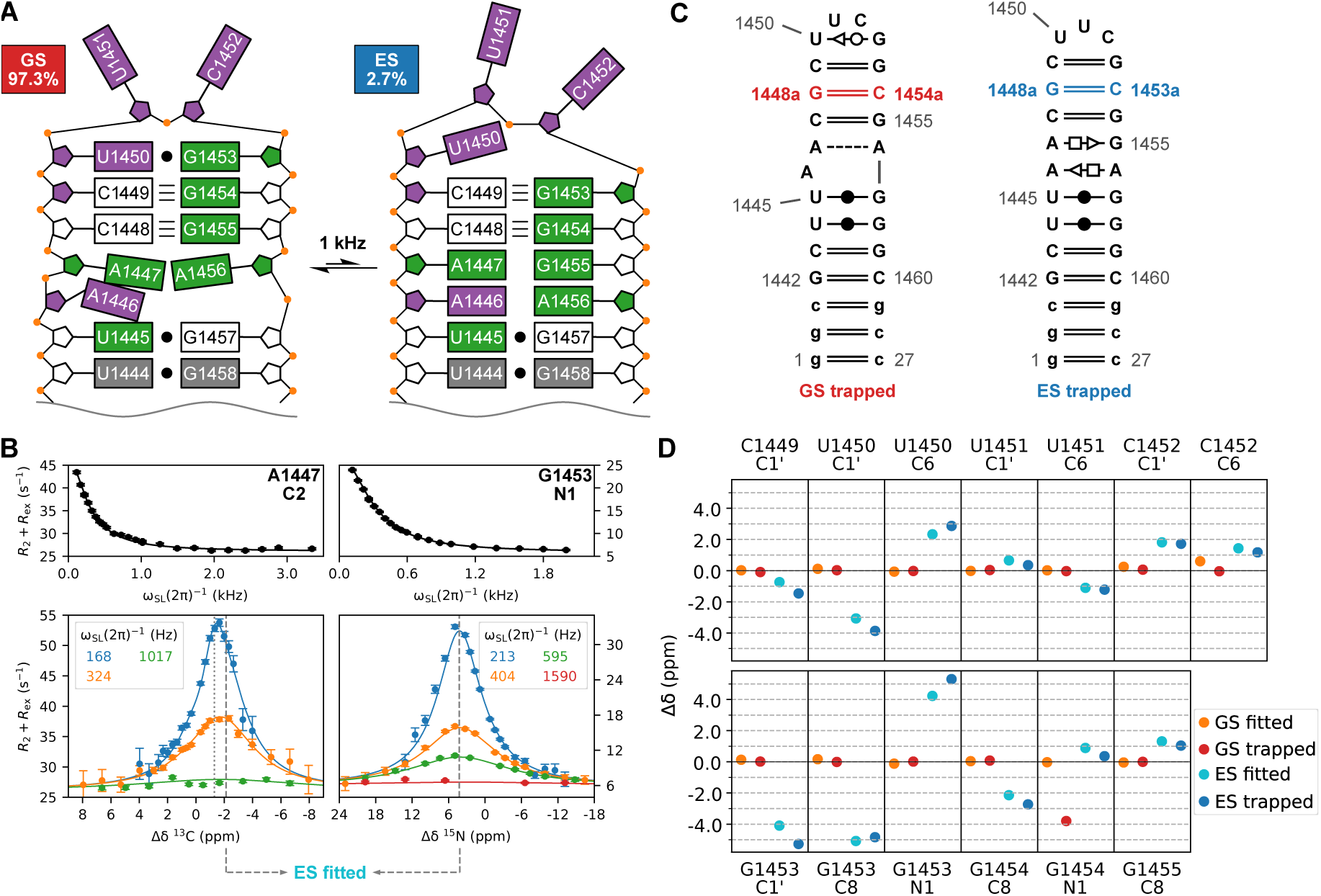
Identification of a reversible secondary structure switch disrupting the A-minor tertiary motif. (**A**) Equilibrium between the A-minor-engaged ground state (GS) and the A-minor-disengaged excited state (ES) in h44-top^WT^, with chemical shift changes derived from *R*_1ρ_ RD experiments (panel (B), **Figure S2**, **Tables S2** and **S3**) indicating the structural signature of the ES. Chemical shifts changes characteristic for helical RNA (green), unstacked/*syn*-nucleobase orientation, small exchange contributions (grey), or C2’-endo ribose pucker (violet) are indicated. Nucleotides 1453–1456 undergo a register shift towards the 3’-direction, resulting in a rearrangement from a UUCG tetraloop to a UUC triloop and concomitant collapse of the adenosine-rich bulge into a more helical conformation, which sequesters A1446 and A1447 from the A-minor interaction with 16S rRNA h8 (**B**) Representative on-and off-resonance *R*_1ρ_ RD curves of A1447-C2 (bulge) and G1453-N1 (loop) reveal concerted millisecond-timescale exchange, with a 2.7% ES population (dashed lines: fitted ES chemical shifts, dotted lines: minor additional ES). (**C**) Trapped constructs designed to lock the equilibrium in either state: trapped h44-top^GS^ (left) inserts G1448a:C1454a (red) to stabilize the engaged conformation. In contrast, trapped h44-top^ES^ (right) inserts G1448a:C1453a (blue) to stabilize the disengaged conformation. Base pairs are annotated using Leontis-Westhof nomenclature^47^. (**D**) Mutate-and-chemical-shift-fingerprint (MCSF)^13^ plot comparing the *R*_1ρ_-derived chemical shifts of the exchanging GS and ES in h44-top^WT^ (orange and cyan, respectively, **Figure S2**, **Table S2**) to the observed chemical shifts in the respective trapped h44-top^GS^ (red) and h44-top^ES^ (blue) constructs (Assignments in **Figures S3** and **S5**). G1454-N1 is an outlier due to proximity to the insertion site. Chemical shift changes (Δδ) are referenced to h44-top^WT^ (**Figure S1**). Excellent agreement between the *R*_1ρ_-derived ES fingerprint and the h44-top^ES^ construct confirms the register shift model.

### Mutational trapping captures both states of the A-minor switch

To directly test whether the register-shift model accounts for the excited state fingerprint observed by relaxation dispersion NMR, we sought to trap the predicted conformations by rational mutagenesis. Secondary structure predictions of h44-top^WT^ support the experimentally derived excited state model, placing the cUUCg triloop fold at +1.8 kcal mol^−1^ relative to the ground state. To further substantiate the proposed conformational exchange process, we employed the Mutate-and-chemical-shift-fingerprint (MCSF) approach^13^ and added a G:C WC base pair to the two constructs to stabilize either the ground state or excited state structure and prevent their interconversion (**Figure 2C**): G1448a was inserted into the stem-loop between C1448 and C1449 to form a base pair with C1454a (inserted between G1454 and G1455) in h44-top^GS^, while in h44-top^ES^ the inserted G1448a forms a base pair with C1453a (inserted between G1453 and G1454) instead. The ground state construct recapitulates the dominant ground state conformation of the WT construct as shown by near-identical ^1^H, ^13^C and ^15^N chemical shifts (**Figures S3**, **S4** and **S7**). Likewise, the chemical shifts of the excited state construct are in excellent agreement with the excited state fingerprint obtained from relaxation dispersion analysis (**Figure 2D**, **Figures S5** to **S7**), confirming that the triloop fold was stabilized.

### The conserved A-minor switch is a general feature

The trapped h44-top excited state suggests that the A-minor interaction of A1446 and A1447 with h8 can be disrupted by locking the adenosines in an intrahelical stack. This is reminiscent of the ribosomal decoding center, where A1492 and A1493 form an A-minor interaction with the codon–anticodon minihelix.^15^ These adenosines have been shown to undergo a transient intrahelical flipping accompanied by compensatory rearrangements in the surrounding bulge, including flipping of U1495.^17^ This parallel prompted us to investigate whether reversible sequestration of A-minor contacts represents a more general dynamic motif in the ribosome.

To identify such motifs, we devised a workflow combining (1) structural mining to define a minimal construct around each motif, (2) secondary structure ensemble prediction and (3) mutational trapping (**Figure 3**). To start with, a geometric search of a high-resolution *E. coli* 70S ribosome structure (PDB 7K00) using FR3D^31,32^ identified 276 individual adenosines engaged in A-minor interactions (**Figure 1D**). Of these, 206 occur in secondary structure contexts compatible with the extra-intrahelical transition proposed here (**Figure 3A**, **Supplemental Note S3.2.1**). Each site can contain one or multiple adenosines. These sites were embedded in minimal hairpin constructs comprising the A-minor adenosine(s), three flanking base pairs on either side plus any unpaired nucleotides in between, and a closing loop (**Figure 3B**, **Supplemental Note S3.2.2**), resulting in 52 test cases to perform (2) prediction of the secondary structure ensemble. This secondary structure ensemble prediction with MC-Flashfold^33^ revealed that many of these hairpins could undergo switching of A-minor adenosines between extra-and intrahelical states **(Figure 3C).** Two mechanistic scenarios emerged: (a) a register shift is accompanied by a change in native apical loop size, analogous to h44-top. Or (b) a bulge translocation along the helix couples adenosine sequestration to opening of a bulge elsewhere in the stem, reminiscent of the A-site and even present in non-A-minor bulges such as HIV-DIS.^17^ After excluding constructs with end fraying or insufficient structural context (**Supplemental Note S3.2.3**), eight well-structured candidates were used to study switching behavior within a physiologically relevant free-energy range (2–8 kcal mol⁻¹), including h44-top and the A-site.^17^ For four of the newly identified sites (h41, HII/III-middle, H2-middle and H28-top), targeted point mutations were sufficient to bias the ensemble heavily toward either ground state or excited state adenosine conformations (**Figure 3D**, **E**, **Figure S8**, **Table S1**), demonstrating that mutations can selectively stabilize either adenosine conformation at these sites and therefore influence A-minor switching.

**Figure 3.**
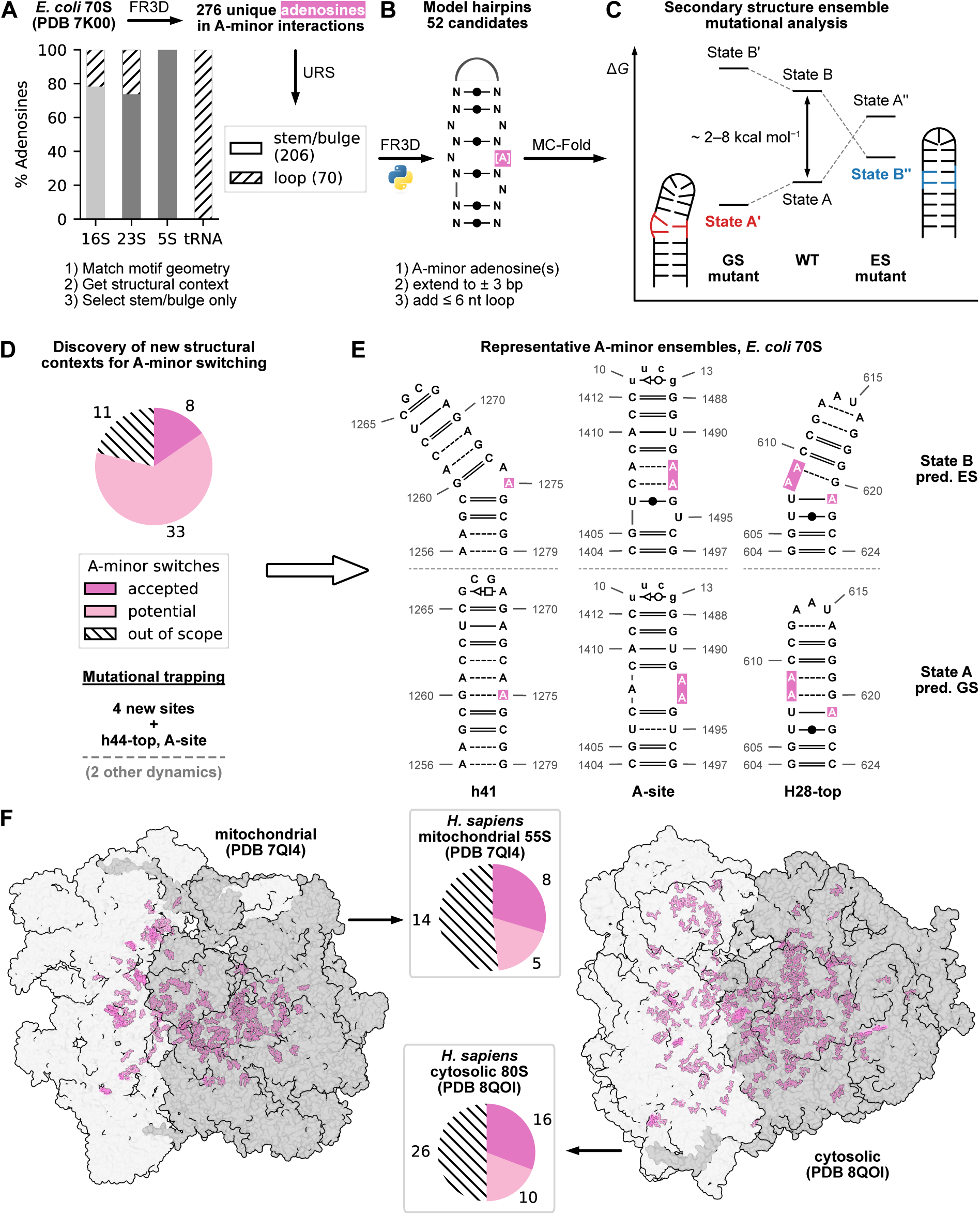
Identified A-minor switches in the ribosome. Workflow for structural mining, construct design, secondary structure ensemble analysis, and mutational trapping of A-minor switches. (**A**) A-minor sites in the *E. coli* 70S ribosome (PDB 7K00) were identified based on interaction geometries using FR3D and classified according to their secondary structure environment using the URS database (**Supplemental Note S3.2.1**) ^48^. Adenosines in hairpin loops (hatched) were excluded from further analysis. (**B**) 52 A-minor motifs (pink) comprising one or more of the 206 single adenosines were selected. Constructs included three native flanking base pairs (FR3D-annotated) and any unpaired nucleotides (N) present in the sequence context on either side and were capped with a native or artificial closing loop (**Supplemental Note S3.2.2**). (**C**) Secondary structure ensemble predictions with MC-Fold were used to assess A-minor switching between alternative conformations. (**D**) Eight well-structured A-minor switches within the relevant free energy range were identified, including the previously characterized A-site and h44-top switches (**Supplemental Note S3.2.3**). For four newly identified sites, mutational trapping of the ground state (GS) and excited state (ES) were feasible (**Table S1**). An additional 33 A-minor switches (potential) were identified but exhibited end fraying, indicating missing secondary structure context, while 11 A-minor sites involved more complex structural rearrangements requiring extended structural context (out of scope). (**E**) Representative GS and ES conformations of A-minor switches (pink) illustrate two mechanistic classes: compensatory bulge shifts (A-site) and loop register shifts (h41, H28-top). (**F**) Following the same workflow, A-minor switches were identified in the *H. sapiens* mitochondrial 55S (left/top, eight motifs) and cytosolic 80S (right/bottom, sixteen motifs) ribosomes, highlighting the pervasiveness of this dynamic 3D interaction (color code as in panel (D)).

Together with h44-top and the A-site, this establishes six well-structured and experimentally tractable test cases of switchable A-minor motifs distributed across the bacterial ribosome. These six candidates represent only a subset of the predicted switching repertoire. An additional 33 motifs exhibited alternative conformations consistent with A-minor sequestration, but involved more complex secondary structure rearrangements, such as dangling ends. Those were excluded from further testing because larger structural segments would have to be considered. The remaining 11 cases displayed A-minor sequestration combined with more significant structural rearrangements, likely reflecting the absence of extended secondary or tertiary structure context in the minimal hairpin models (**Figure 3D**). Thus, while only six motifs were suitable for mechanistic validation and mutational trapping, the structural mining suggests that reversible A-minor switching is substantially more widespread. At a minimum, the 33 intermediate-complexity cases are strong candidates for such dynamics, and A-minor exchange cannot be excluded for the remaining motifs without considering larger structural assemblies.

Using the same approach to identify A-minor switches in other families of rRNA, eight candidates emerged for the human mitochondrial 55S ribosome (PDB 7QI4), while the human cytosolic 80S ribosome (PDB 8QOI) yielded a further sixteen candidates (**Figure 3F**). Both constitute 30% of the total A-minor motifs annotated in either ribosome, while an additional ∼20% of sites are found to have the potential for switching; their secondary structures, however, are more complex, as described for the bacterial ribosome. The A-minor switch is therefore conserved across domains of life.

### Trapping A-minor switches alters ribosome function

At the decoding center, transient A-minor interactions formed by A1492 and A1493 are essential for accurate codon recognition.^34^ Mutations in the adjacent U1406:U1495 mismatch alter the dynamics of those adenosines both in silico^35^ and in vitro,^17^ and affect subunit association^36^ as well as antibiotic resistance.^37^ These observations suggest that perturbing the A-minor switching mechanism may have functional consequences beyond local structural effects.

To test this hypothesis, we selected two newly identified switches (h41 and H28-top) for functional analysis alongside h44-top and the A-site. Mutations designed to stabilize either the ground state or excited state conformers and therefore inhibit A-minor switching were evaluated for biological impact. The respective mutations (**Figure 3E**, **Table 1**, **Table S1**) were introduced into 16S or 23S rRNA and expressed in *E. coli* SQ171fg, a strain in which ribosomes are produced exclusively from a plasmid-encoded *rrnB* operon without genomic WT background.^38^

**Table 1.**
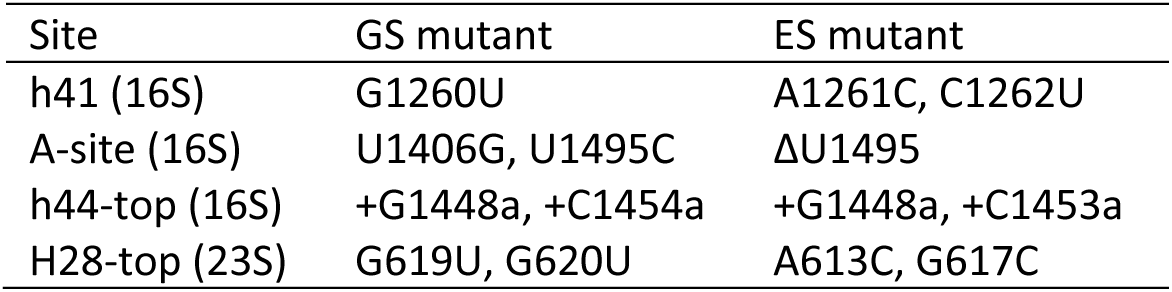
16S and 23S rRNA mutations incorporated into *E. coli* SQ171fg. WT pAM552 was subjected to site-directed mutagenesis to affect GS–ES equilibria in rRNA. + denotes insertion mutations 3’ of the given nucleotide, Δ indicates deletions.

| Site | GS mutant | ES mutant |
| --- | --- | --- |
| h41 (16S) | G1260U | A1261C, C1262U |
| A-site (16S) | U1406G, U1495C | ΔU1495 |
| h44-top (16S) | +G1448a, +C1454a | +G1448a, +C1453a |
| H28-top (23S) | G619U, G620U | A613C, G617C |

Homogenous ground state and excited state mutant strains were obtained for h44-top and h41. In contrast, A-site mutant strains retained a residual WT ribosome background. H28-top mutant strains were not obtained despite repeated attempts, indicating strong deleterious effects (**Supplemental Note S3.3**). Spot growth assays at 37 °C showed that h44-top^GS^ and A-site^GS^ ribosomes supported bacterial growth comparable to WT ribosomes, whereas the corresponding h44-top^ES^ and A-site^ES^ mutants displayed significantly reduced growth (**Figure 4A**, **Figure S9**). Given the residual WT background in the A-site^ES^ strain, the true fitness defect is likely underestimated (**Supplemental Note S3.3**). Mutations in h41 yielded the opposite pattern, with h41^GS^ showing impaired growth while h41^ES^ remained unaffected (**Figure 4B**). It is therefore expected that adenosines need to be able to move into a flexible bulge conformation for A-minor switching. The growth effects were amplified at 30 °C (**Figure S10**), indicating that ribosome fitness may be particularly sensitive to perturbation of A-minor switching under thermal stress conditions.

**Figure 4.**
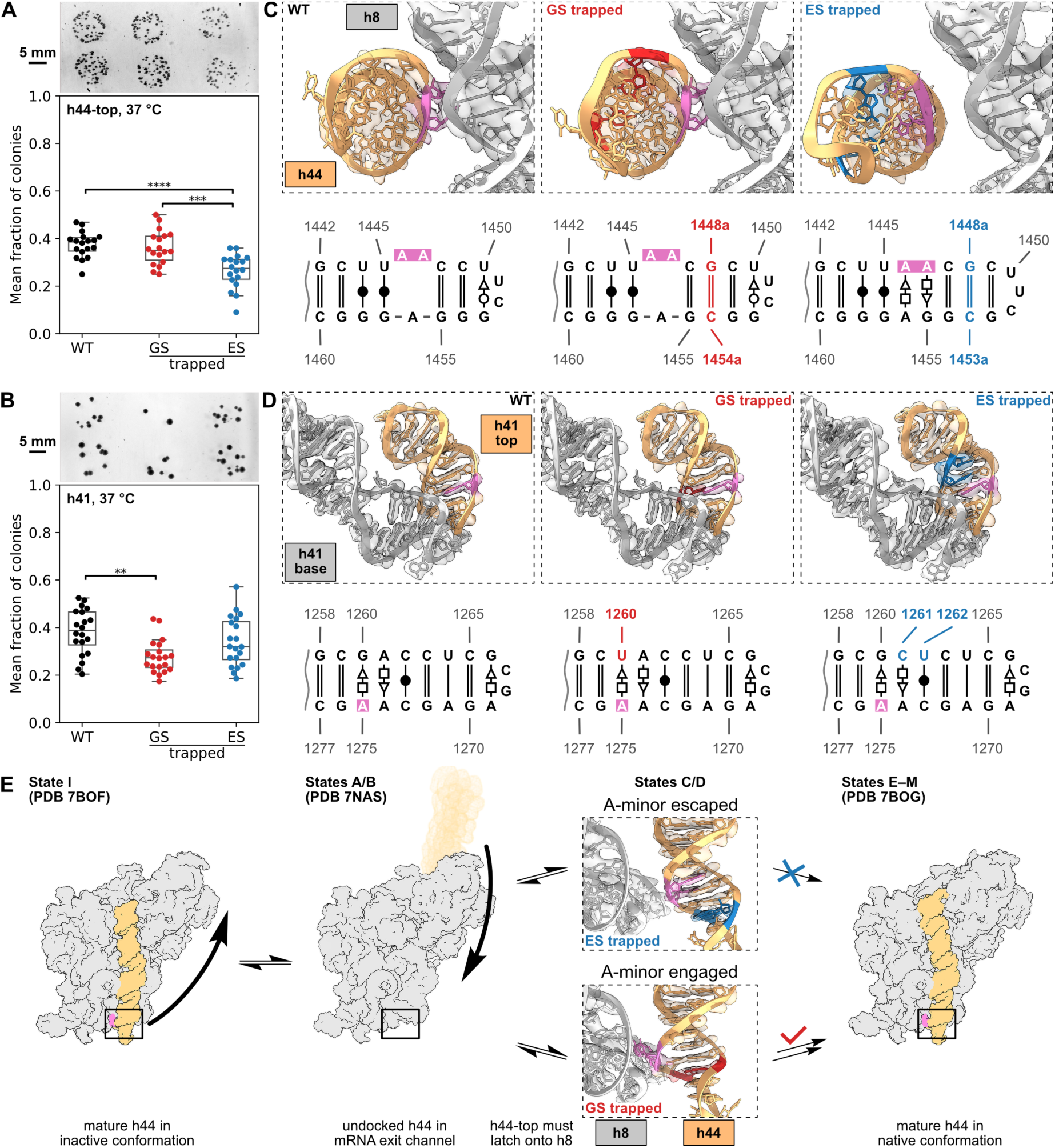
Mutational interference with A-minor switching affects bacterial viability. (**A**, **B**) Spot growth analysis of *E. coli* SQ171fg shows that trapped h44-top^ES^ (A) and h41^GS^ (B) in 16S rRNA lead to reduced bacterial growth relative to the WT at 37 °C, while the respective complementary trapped states have no significant effect. The direction-dependent fitness phenotype (h44-top sensitive to disengagement, h41 sensitive to engagement) indicates that productive function requires the *capacity* to switch, not a single static conformation. Scale bars indicate 5 mm length. Statistical analysis: one-way ANOVA with Bonferroni correction for multiple means comparisons. ** *P* < 0.01, *** *P* < 0.001, **** *P* < 0.0001 (**Figures S9** and **S10**). (**C**) Cryo-EM structures of unrotated h44-top (orange) carrying the A-minor interaction (pink) to h8 (light grey) is switched off in the trapped ES (blue, right) but remains intact in the trapped GS (red, middle) compared to h44-top^WT^ (black, left). Secondary structures of the orange region are shown below with matching color-coding (**Figure S11**). The h44-top^ES^ cryo-EM density directly visualizes the NMR-defined excited state inside the intact 70S ribosome, providing rare structural validation of an RD NMR-derived ES within its full biological context. (**D**) In 16S rRNA h41, the apical stem-loop (orange) interacts with the lower stem (light grey) of the same helix (**Figure S12**). The type III A-minor interaction of A1275 (pink) is unaffected by both GS (red, middle) and ES (blue, right) mutations in the mature 70S ribosome. Secondary structures of the orange region are shown below with matching color-coding. (**E**) Model: late-stage 30S maturation requires h44 to dock onto the SSU body and latch onto h8 via the apical loop A-minor contact^49^. h44 cycles between body-docked (state I) and exit-tunnel-disordered (states A/B) conformations during maturation; we propose that competent latching (states C/D) requires the A-minor-engaged GS conformation of h44-top. Trapping h44-top in the A-minor-disengaged ES prevents productive latching and stalls maturation, consistent with the observed fitness defect. These A-minor switching events function as a regulator not only of translation (decoding-center A-site^17^) but also of ribosome maturation, together implicating A-minor switching as a general mechanism that couples local secondary-structure dynamics to large-scale RNA function.

To determine whether these phenotypes arise from structural alterations in intact 70S ribosomes, we performed cryo-EM analysis of ribosomes isolated from WT, h44-top mutant and h41 mutant SQ171fg.

After consensus reconstruction and classification into rotated and unrotated states, high-resolution reconstructions (2.0–3.5 Å) were obtained and atomic models were built for representative functional states (**Table S7**).

In h44-top^GS^ ribosomes, the apical bulge adopts the WT conformation, with A1446 and A1447 engaged in an A-minor interaction with h8 (**Figure 4C**). Density for the inserted base pair G1448a:C1454a is clearly visible, and the UUCG tetraloop adopts its canonical fold. In contrast, h44-top^ES^ ribosomes displayed the register-shifted stem and the tandem mismatch that sequester A1446 and A1447 intrahelically (A1446:A1456 *trans* Sugar edge/Hoogsteen and A1447:G1455 *trans* Hoogsteen/Sugar edge), thereby switching off the A-minor interaction with h8 (**Supplemental Note S3.4**). The register-shifted upper stem is well defined, while the UUC triloop density is weaker (**Figure S11**). The h44-top^ES^ structure matches the NMR-derived excited state model, providing a rare example of direct validation of the divide-and-conquer approach within the intact ribosome. Structural differences between h44-top^ES^, h44-top^GS^ mutant, and WT ribosomes were confined to the immediate vicinity of the A-minor motif and mutation sites, with no detectable global rearrangements in unrotated or rotated states (**Figure S13A**, **B**).

Similarly, h41^GS^ and h41^ES^ mutations were clearly resolved in the density (**Figure 4D**, **Figure S12**). The cryo-EM models suggest that the noncanonical interaction network within h41 is neither strongly stabilized nor destabilized by the introduced ground state and excited state mutations, and that no global structural changes in the ribosome are present. Nonetheless, h41^GS^ mutation G1260U, which yields a *trans* Sugar edge/Hoogsteen base pair with A1275, significantly impairs bacterial growth. In contrast, the type III A-minor interaction of A1275 with the A1254:U1283 base pair is preserved in h41^ES^ ribosomes (**Figure S13C**, **D**).

Together, these findings indicate that the fitness defects associated with trapped A-minor switches are not due to structural distortion of the mature 70S ribosome, i.e. they do not impair translation. Instead, they likely arise from perturbation of transient conformational equilibria during ribosome biogenesis.

Multiple studies support a dynamic role for h41 and h44 during 30S assembly. In-cell SHAPE probing has revealed alternative secondary structures in these two and several other 16S rRNA helices under different growth conditions.^39^ Additionally, chemical modification of h41 interferes with SSU head assembly.^40^ Proper folding of h44 represents one of the final checkpoints in 30S maturation,^41,42^ and inactive conformations in which h44 is displaced from the SSU body have been observed by in-cell SHAPE and cryo-EM.^43–46^ Stabilization of an A-minor-off conformation in h44, or restricting transient flipping events in h41, may therefore hinder critical maturation steps and reduce the abundance of translationally competent ribosomes (**Figure 4E**).

We therefore propose that A-minor switches act not only during translation, as exemplified by the A-site, but also during ribosome assembly. These local secondary structure equilibria, which transiently permit or sequester A-minor contacts, appear to be encoded within ribosomal RNA architecture as regulatory elements guiding biogenesis and functional activation.

A-minor interactions have long been regarded as static structural elements of RNA. Our findings challenge this view by demonstrating that these ubiquitous tertiary contacts can be dynamically altered through RNA secondary structure rearrangements. Using relaxation dispersion NMR, large-scale structural data analyses, mutational trapping, bacterial growth assays and cryo-electron microscopy of intact ribosomes, we identify a register shift termed the A-minor switch. We demonstrate that stabilizing the engaged or disengaged conformational states impairs bacterial growth, directly linking RNA excited state dynamics to cellular phenotype. Beyond the well-characterized decoding center adenosines integral to translation fidelity, structural mining indicates that similar switching motifs are encoded at multiple ribosomal sites across domains of life. Capturing an NMR-defined excited state within the intact 70S ribosome using cryo-EM establishes that such dynamic equilibria are intrinsic features of RNA structure that introduce a previously unrecognized layer of regulation. A-minor motifs are pervasive in structured RNAs, such as riboswitches, group I and II introns, RNase P, and kink-turn-containing RNAs. Therefore, wherever an A-minor contact is supported by a flanking secondary-structure context that admits an alternative pairing, the same switching mechanism is, in principle, available. Our ensemble-prediction workflow is sequence-and RNA origin-agnostic and can be applied to any structured RNA whose secondary structure is predictable. We therefore expect A-minor switching to be a general regulatory layer of RNA tertiary structure. These dynamic A-minor switches reveal a previously unrecognized class of druggable RNA states, that open new opportunities for therapeutic development by targeting transient RNA conformational states, e.g. novel antibiotics targeting the ribosome.

## Methods

### In vitro transcription

Uniformly ^13^C,^15^N-labeled RNA oligonucleotides were prepared by in vitro transcription with T7 RNA polymerase according to previously reported procedures^50^.

Briefly, DNA template and T7 promoter, each at a concentration of 25 µM, were annealed in a 3 mM solution of MgCl_2_. Transcription reactions were performed at 37 °C for 16 h in a total volume of 10 mL containing for example 0.4 µM annealed DNA, 100 mM Tris-HCl pH 8.0, 60 mM MgCl_2_, 30 mM DTT, 2 mM spermidine, 3 mM of each NTP and GMP, 0.2–0.4 mg/mL T7 RNAP, 0.025 mg/mL IPPase, 0.1 U/µL RNaseOut and 15% DMSO. The exact concentrations were optimized for each template to achieve optimal yields with minimal production of ±1 nt products. Transcripts were purified by one to two rounds of ion-exchange HPLC.

### Solid-phase synthesis

Unlabeled RNA oligonucleotides were prepared by solid-phase synthesis on a K&A H-8 using 5’-*O*-DMT/2’-*O*-TBDMS-protected 3’-β-cyanoethyl phosphoramidites of *N*^6^-benzoyladenosine, *N*^4^-acetylcytidine, *N*^2^-acetylguanosine and uridine (purchased from Glen Research) with activator ETT (purchased from Sigma Aldrich).

The unlabeled h44-top^WT^ oligonucleotide was synthesized on a 1 µmol scale according to standard protocols. All phosphoramidites were dissolved in anhydrous MeCN at a concentration of 50 mmol L^−1^ and the coupling time was 12 min. Cleavage from the solid support was performed with a 1:1 mixture of 25% aq. NH_3_ and 40% aq. MeNH_2_ for 30 min at ambient temperature. For complete removal of the base-labile protecting groups the solution was then heated to 65 °C for 30 min. Silyl deprotection was performed with a mixture of Et_3_N·3HF and Et_3_N in anhydrous DMSO at 65 °C for 2.5 h. The crude RNA was desalted by ethanol precipitation and purified by ion-exchange HPLC.

### Ion-exchange HPLC purification

In vitro transcription and solid-phase synthesis products were purified by preparative anion-exchange HPLC on a Thermo Scientific Ultimate 3000 system equipped with a DNAPac PA200 column (22×250 mm) at 75 °C. Buffer A: 20 mM NaOAc pH 6.5, 20 mM NaClO_4_, 10% MeCN. Buffer B: 20 mM NaOAc pH 6.5, 600 mM NaClO_4_, 10% MeCN. A typical gradient was for example 15–35% buffer B in 10 CV at a flow rate of 5 mL/min. The gradient was optimized for each construct to achieve optimal separation^50^.

HPLC fractions were analyzed by denaturing PAGE (15% acrylamide/bis-acrylamide 19:1, 8 M urea) using 1x TBE (89 mM Tris, 89 mM boric acid, 2 mM EDTA, pH 8.3) as running buffer. Gels were stained with SYBR Gold according to the manufacturer’s protocol and imaged with a GE Healthcare ImageQuant LAS 4000. Fractions containing sufficiently pure RNA were pooled and desalted using Amicon Ultra centrifugal filters with a 3 kDa MWCO.

### NMR sample preparation

Desalted RNA after ion-exchange HPLC purification was exchanged into NMR buffer (15 mM NaP, 25 mM NaCl, 0.5 mM EDTA, pH 6.5) using Amicon Ultra centrifugal filters with a 3 kDa MWCO. Folding of the hairpins was performed at an RNA concentration of 20–30 µM by heating to 95 °C for 5 min followed by snap cooling on ice for 30 min. Samples were concentrated to a final volume of 250 µL including 10% D_2_O for locking and transferred into 5 mM Shigemi tubes.

### NMR resonance assignment

The set of experiments for resonance assignment included ^1^H–^1^H NOESY and SELOPE^12^ on unlabeled samples as well as ^1^H–^13^C HSQC of the aromatic, anomeric and ribose regions, ^1^H–^15^N HSQC of the imino and amino regions, 2D HCN to correlate H6/8 and H1’ to N1/9 in the same nucleotide, and HNN-COSY to correlate N1 with N3 across hydrogen bonds in Watson-Crick base pairs on ^13^C,^15^N-labeled samples. ^1^H chemical shifts are referenced to DSS as an external standard; ^13^C and ^15^N referencing were calculated from the ^1^H value using the respective Ξ ratios^51^ (ν_X_ = ν_DSS_ × Ξ_X_ / 100). NMR spectra were assigned with the help of CcpNmr^52^ and plotted using custom python scripts.

### NMR relaxation dispersion

Rotating frame relaxation rates (*R*_1ρ_) were analyzed with analytical expressions reported by London and coworkers^53^, Palmer and coworkers^54,55^ or Al-Hashimi and coworkers^56,57^, respectively, for different kinds of exchange models, which are described in detail in the supplemental information.

### Preparation of WT and mutant ribosome *E. coli* strains

Plasmid pAM552 containing the WT *rrnB* operon was used to generate point mutations in the 16S and 23S rRNA for trapping GS and ES RNA conformations. Sequences of mutagenic forward and reverse primers for h44-top, h44-A-site and h41 in the 16S rRNA as well as H28 in the 23S rRNA (designed using NEBaseChanger) are given in **Table S1**. A Q5 site-directed mutagenesis kit (New England Biolabs E0554S) was used according to the manufacturer’s protocol.

For each mutant, between three and five colonies were picked, cultured overnight (LB, ampicillin, 5 mL, 37 °C), and the plasmids were isolated by Miniprep (QIAGEN QIAprep Spin Miniprep Kit 27104 or Thermo Fisher Scientific PureLink Quick Plasmid Miniprep Kit K210011) from a 1.5 mL aliquot of the culture according to the manufacturer’s protocol. Successful mutagenesis was confirmed by Sanger sequencing (Eurofins Genomics) using the forward and reverse primers for each site given in **Table S1**. When positive, the remainder of the culture was expanded to 200 mL, plasmids were isolated by Maxiprep (MACHEREY-NAGEL NucleoBond Xtra Maxi 740414.50M) and checked again by Sanger sequencing.

Additionally, for h44-A-site mutants, WT pAM552 was double-digested with XbaI and KpnI in 1x CutSmart buffer for 2 h at 37 °C to remove the 16S sequence. CIP was added to the reaction for simultaneous removal of 5’-phosphate. The linearized plasmid was purified by agarose gel electrophoresis (0.5%, QIAGEN QIAquick Gel Extraction Kit 28704). Commercially sourced inserts for the h44-A-site mutant 16S sequences with the same restriction sites (IDT gBlocks) were double-digested as above without dephosphorylation and purified using a PCR cleanup kit (Invitrogen K220001) according to the manufacturer’s protocol. Linearized plasmid and the respective insert were ligated with T4 DNA ligase (10 µL reaction in 1x T4 DNA ligase reaction buffer, NEB B0202S), 5 µL of the ligation mixture were added to 25 µL OneShot TOP10 chemically competent *E. coli* (Thermo Fisher Scientific C404010) and incubated on ice for 10 min, followed by heat shock at 42 °C for 30 s and further incubation on ice for 2 min. The transformed bacteria were streaked on LB agar with ampicillin, selected colonies were cultured overnight, plasmids were isolated by Miniprep and checked by Sanger sequencing.

Following a published protocol^58^, WT and mutant pAM552 plasmids (ampicillin-resistant) were transformed into CaCl_2_-competent *E. coli* SQ171fg carrying pCSacB (WT *rrnB* operon, kanamycin-resistant, sucrose-sensitive) and ptRNA67 (spectinomycin-resistant). 150 µL of the transformation mix was diluted with 1.85 mL of S.O.C. medium including ampicillin, spectinomycin and 0.25% sucrose and incubated overnight with shaking at 37 °C. A small amount was then cultured on LB agar with ampicillin, spectinomycin and 5% sucrose. At this stage, the plasmids were isolated again by Miniprep and checked by Sanger sequencing before proceeding with spot plating and ribosome isolation for cryo-EM. Expulsion of pCSacB following successful transformation was checked by PCR with the primers given in **Table S1**.

N.B.: These procedures yielded pure h44-top^GS/ES^ and h41^GS/ES^ mutant ribosomes, whereas h44-A-site^GS/ES^ contained residual WT (not detectable by PCR). Mutations to H28 were unsuccessful (see also **Supplemental Note S3.3**).

### Spot plating

*E. coli* SQ171fg carrying WT or h44-top/h44-A-site/h41 mutant ribosome plasmids were cultured overnight (LB, ampicillin and spectinomycin, 5 mL, 37 °C) and the OD was determined (1 OD = 8 × 10^8^ cells mL^−1^). An LB agar plate containing ampicillin and spectinomycin was spotted with the following dilutions (20 µL each):

- 1 × 10^3^ cells mL^−1^ of WT, GS or ES, each in duplicate
- 8 × 10^2^ cells mL^−1^ of WT, GS or ES, each in duplicate

i.e. 12 spots for counting of colonies. As a control, 20 µL each of 8 × 10^7^ cells mL^−1^ for WT, GS and ES mutants were spotted in duplicate. The duplicates constitute technical replicates. On a given day, four or six such plates were prepared for a set of WT and the respective mutants. Overnight, half of the plates were incubated at 37 °C and the other half at 30 °C. This procedure was repeated an additional two times starting from freshly prepared overnight cultures as biological replicates. Plates were imaged on a black tray using an ImageQuant LAS 4000 (GE Healthcare) with 1/15 s exposure time and colonies were counted. Counts from the technical replicates on each plate were averaged and divided by the sum of all averages for the same plate to normalize against the total number of colonies. For a given temperature, the resulting mean fractions of colonies across multiple plates and biological replicates were aggregated as a set of sample observations with normally distributed mean and standard deviation for the WT, GS mutant and ES mutant. The statistical significance of differences between samples was assessed by one-way ANOVA using the Bonferroni method for multiple comparisons.

### Cryo-EM sample preparation

70S ribosome were isolated from SQ171fg cells, a modified *E. coli* strain devoid of genomic rrn operons^59^ expressing WT and mutated GS and ES ribosomes from a single plasmid based *rrnB* operon as described above. Briefly, the cells were grown in LB medium containing ampicillin and spectinomycin. Cells were pelleted by centrifugation and resuspended in opening buffer (20 mM Tris-HCl, 30 mM NH_4_Cl, 10 mM MgCl_2_, 150 mM KCl, 0.5 mM EDTA, 1 mM DTT) before addition of DNase. Cells were sonicated and the lysate cleared by centrifugation at 30000 × g. The ribosomes were purified on a sucrose cushion (20 mM HEPES-K, 50 mM NH_4_Cl, 5 mM Mg(OAc)_2_, 4 mM 2-mercaptoethanol, 1 M sucrose) in a Ti-45 rotor at 36500 rpm for 1 h. The pellet was washed and resuspended in washing buffer (20 mM HEPES-K, 50 mM NH_4_Cl, 5 mM Mg(OAc)_2_, 4 mM 2-mercaptoethanol). The sample was prepared for cryo-EM immediately after purification.

Quantifoil R2/2 200 mesh copper grids with 2 nm continuous carbon were glow-discharged for 2 min at 25 mA using an EMS 100 glow discharge unit. 3 µL of the respective ribosome sample were incubated on a grid for 30 s, blotted for 1 s with blot force 0, and plunge-frozen in liquid ethane using a Vitrobot Mark IV at 4 °C and 100% humidity.

### Cryo-EM data acquisition and image processing

Cryo-EM images were collected using a Titan Krios G3i microscope (Thermo Fisher Scientific) operating at 300 kV equipped with a K3 BioQuantum detector and energy filter using a 10 eV slit width (Gatan) at the Karolinska Institutet 3D-EM facility in Solna, Sweden. The data was acquired using EPU software (Thermo Fisher Scientific) with a pixel size of 0.505 Å/px (nominal magnification 165000x), electron dose of ∼50 e^−^/Å^2^, 50 frames per movie, and a defocus range of −0.3 to −1.5 µm.

The total number of movies per sample is reported in **Figures S14 to S18** and **Table S7**.

All processing was done in CryoSPARC^60^ (versions 4.4.1–4.6.2). Masks for 3D variability analysis were generated in ChimeraX^61^ (version 1.8).

Gain-corrected movies were imported, subjected to patch motion correction with dose weighting followed by patch CTF estimation and good exposures were curated.

Particles were picked as circular blobs (230–250 Å) and suitable picks were extracted with a box size of 864 px Fourier-cropped to a reduced size of 216 px for 2D classification. Obvious junk classes were excluded, and good classes were used as input for template picking. The results from both rounds of picking were combined with removal of duplicates, particles were re-extracted and subjected to a second round of 2D classification. Good classes were used for ab-initio reconstruction with five to ten classes. All volumes resembling intact 70S ribosomes as well as one junk volume were subjected to heterogenous refinement. The resulting ribosome classes were used for homogeneous refinement (minimizing over per-particle scale, optimizing per-particle defocus, optimizing per-group CTF parameters) with particles re-extracted at a 576 px Fourier-crop box size to generate a consensus map.

3D classification with ten classes and a filter resolution of 6 Å followed by homogenous refinement was used to separate higher-resolution (∼2 Å) unrotated and rotated ribosomes from lower-resolution particles that were not processed further. Reference-based motion correction was applied, and homogenous refinement was repeated with negative EWS correction to obtain maps of unrotated and rotated ribosomes with mixed tRNA occupancies.

To generate masks for 3D variability analysis^62^ of tRNA occupancy, existing models of unrotated^29^ (PDB 7K00) and rotated^63^ (PDB 8R8M) ribosomes were rigid-body-docked into the respective maps in ChimeraX. Mask bases were generated around the tRNA chains with a resolution of 16 Å on the grid of the input map using the ‘molmap’ command, binarized at an appropriate threshold, dilated by 5 px and padded by 12 px. 3D variability analysis was performed with a filter resolution of 8 Å to generate three principal components in cluster mode. This yielded unrotated ribosomes with tRNA in the A and P site (used for model building), tRNA in the P site only, and tRNA-free, as well as rotated ribosomes with tRNA in the a/P and p/E state (used for model building), A and p/E state, and p/E state only. The global resolution of each reconstruction was determined by half-map FSC (0.143 cutoff) using the tight mask generated by CryoSPARC during homogeneous refinement, and maps filtered to the local resolution were calculated to aid in model building.

Processing details for each sample are given in **Figures S14 to S18**.

### Model building and refinement

To calibrate the magnification of the cryo-EM maps, a high-resolution crystal structure of the bacterial ribosome^64^ (PDB 4YBB) was rigid-body-docked into the unrotated 70S reconstructions in ChimeraX and the voxel size was adjusted until maximal map-to-model correlation was reached.

For the unrotated WT ribosome, a previous cryo-EM model^29^ (PDB 7K00) was fitted first as a rigid body and then chain-by-chain into the unsharpened cryo-EM map using ChimeraX. Partially modeled protein bL9 was removed and replaced with a complete AlphaFold model, which was rigid-body-docked into the corresponding density. The antibiotic spectinomycin was placed into its known binding site^65^ in SSU h34 and the assembled model was manually adjusted with Isolde. This model was refined against unsharpened half-maps with jelly-body restraints using the refine_spa_norefmac routine of Servalcat^66,67^ (version, 0.4.88) followed by inspection and manual adjustments as needed in Coot^68^ (version 0.9.8.95). While no particular individual tRNAs or mRNA were employed during ribosome isolation, the map density supported identification of modified nucleotides consistent with A-site tRNA-Val and P-site tRNA-fMet^69,70^. The modeled mRNA fragment was retained from the starting model 7K00. Nucleotides and amino acids not sufficiently supported by density were removed. Mg^2+^ and K^+^ ions and were placed with the help of a locally filtered map from CryoSPARC and the difference map from Servalcat and checked against a previous cryo-EM model^71^ (PDB 8EMM). Metal ion types were distinguished based on their coordination environments^72^. After further Servalcat refinement without jelly-body restraints, water was added to the locally filtered map using phenix.douse. Rounds of manual adjustments in Coot, refinement in Servalcat and validation with MolProbity^73^ and phenix.validation_cryoem were alternated and final refinement was performed with Refmac via the refine_spa routine of Servalcat to obtain the deposited unrotated WT ribosome model.

The unrotated WT model prior to addition of ions and water was used as a starting point for modeling the unrotated mutant ribosomes. For h44-top_GS, 16S rRNA nucleotides 1449–1454 (CUUCGG loop) were moved up in register and insertions G1448a/C1454a were added in Coot. For h44-top_ES, 16S rRNA nucleotides 1446–1457 were removed and modeled de-novo together with insertions G1448a/C1453a into the density of the trapped ES conformation in Coot. For the h41 variants, 16S rRNA mutations G1260U (GS) or A1261C/C1262U were generated in Coot. Afterwards, addition of ions and waters, refinement and validation were repeated as described above for the WT model.

All rotated ribosome models were created analogously using the unrotated WT model prior to addition of ions and water as a starting point. Following chain-by-chain fitting into the rotated WT map, previously published cryo-EM models of rotated ribosomes^63,74^ (PDB 8FZI, 8R8M) were used to guide necessary adjustments and metal ion placement.

Refinement statistics for each model are given in **Table S7**. Visualizations of 3D structures were made with ChimeraX.

### A-minor motif secondary structure ensemble analysis

A detailed description of the analysis pipeline is provided in the supplemental information. RNA secondary structure diagrams were drawn with RNAcanvas^75^.

## Data availability

The gain-corrected, unaligned multiframe micrographs for all five datasets and final particle stacks for each class have been deposited in the Electron Microscopy Public Image Archive (EMPIAR) under accession code EMPIAR-13345. The corresponding cryo-EM maps for each class have been deposited in the Electron Microscopy Data Bank (EMDB) under the following accession codes: EMD-56070, EMD-56091, EMD-56128, EMD-56827, EMD-56828, EMD-56829, EMD-56830, EMD-56831, EMD-56832, EMD-56833, EMD-56834, EMD-56835, EMD-56836, EMD-56837, EMD-56839, EMD-56840, EMD-56841, EMD-56842, EMD-56843, EMD-56844, EMD-56845, EMD-56846, EMD-56847, EMD-56848, EMD-56849, EMD-56850, EMD-56851, EMD-56852, EMD-56853, EMD-56854. Refined atomic models for classes of unrotated ribosomes with A-and P-site tRNA and rotated ribosomes with A/P-and P/E-site tRNA have been deposited in the Protein Data Bank (PDB) under the following accession codes: 9TMI, 9NTL, 9TQA, 28UH, 28UI, 28UJ, 28UK, 28UL, 28UM, 28UN.

Atomic coordinates for the *H. marismortui*, *E. coli*, *H.sapiens* cytosolic or mitochondrial ribosomes and maturation intermediates have been retrieved from the PDB under accession codes 1FFK, 4YBB, 7BOF, 7BOG, 7K00, 7NAS, 7QI4, 8FZI, 8QOI and 8R8M.

## Code availability

Python 3.x scripts for relaxation dispersion data analysis, A-minor motif classification, construct design and secondary structure analysis are available from https://github.com/PetzoldLab or upon request.

## Supporting information

Tabulated data underlying the figures

Spin-lock-dependent experimental and fitted R1rho rates for h44-top_WT in Tables S2 and S3

Spin-lock-dependent experimental and fitted R1rho rates for h44-top_ES in Table S5

Spin-lock-dependent experimental and fitted R1rho rates for h44-top_GS in Table S4

## Acknowledgements

We thank C. S. Mandava and S. Sanyal for sharing and giving advice on working with *E. coli* strain SQ171fg and plasmid pAM552 WT; I. Berzina for advice on cryo-EM sample preparation and data acquisition; D. Larsson, G. Gaullier and A. González López for many valuable suggestions and helpful discussions about various aspects of model building and refinement.

T7 RNA polymerase was prepared by the Karolinska Institutet/SciLifeLab Protein Science Core Facility (http://ki.se/psf). EM data was collected at the Karolinska Institutet 3D-EM facility (https://ki.se/cmb/3d-em).

## Funding

C. S. acknowledges funding by an EMBO postdoctoral fellowship (ALTF 1011-2020). J. S. acknowledges funding by a Marie Skłodowska-Curie Individual Fellowship (EU H2020/project number 747446). K. P. is grateful for funding from the Swedish Foundation for Strategic Research (project no. ICA14-0023), the Swedish Research Council Starting and Consolidator grants (2014-04303 & 2018-00250), Åke Wiberg Stiftelse (467080968 and M14-0109), Karolinska Institutet (career position to K. P. and support for the purchase of a 600 MHz Bruker NMR spectrometer), Harald och Greta Jeansson Stiftelse (JS20140009), Carl Tryggers Stiftelse (CTS14-383 and 15-383), Eva och Oscar Ahréns Stiftelse, the Ragnar Söderberg Foundation (M91/14).

## Author contributions

Conceptualization: CS, HK, CF, KP Data curation: CS, CF, HK

Formal Analysis: CS, HK, CF, NGR, KP Funding acquisition: CS, JS, KP Investigation: CS, HK, CF, NGR, MR, DK Methodology: CS, HK, CF, DK, JS, ES, SFS Project administration: KP

Software: CS Supervision: ERA, KP Visualization: CS

Writing – original draft: CS Writing – review & editing: CS, KP

## Competing interests

The authors declare no competing interests.

## Supplemental information

**Document S1**

Figures S1 to S18

Tables S2 to S6

Supplemental Notes

Supplemental Methods

Supplemental References

**Table S1**

List of DNA and RNA sequences used in this study.

**Table S7**

Cryo-EM data collection, refinement and validation statistics for 70S WT, 70S h44-top^GS^, 70S h44-top^ES^, 70S h41^GS^ and 70S h41^ES^ datasets.

**Data S1**

Source data underlying the figures.

**Data S2**

Experimental parameters and fit results for *R*_1ρ_ relaxation dispersion experiments on h44-top^WT^ RNA.

**Data S3**

Experimental parameters and fit results for *R*_1ρ_ relaxation dispersion experiments on h44-top^GS^ RNA.

**Data S4**

Experimental parameters and fit results for *R*_1ρ_ relaxation dispersion experiments on h44-top^ES^ RNA.

## Notes

### Competing Interest Statement

The authors have declared no competing interest.

## References

1. Cate, J.H., Gooding, A.R., Podell, E., Zhou, K., Golden, B.L., Szewczak, A.A., Kundrot, C.E., Cech, T.R., and Doudna, J.A. (1996). RNA Tertiary Structure Mediation by Adenosine Platforms. Science 273, 1696–1699. 10.1126/science.273.5282.1696.

2. Nissen, P., Ippolito, J.A., Ban, N., Moore, P.B., and Steitz, T.A. (2001). RNA tertiary interactions in the large ribosomal subunit: The A-minor motif. Proc. Natl. Acad. Sci. 98, 4899–4903. 10.1073/pnas.081082398.

3. Baulin, E.F. (2021). Features and Functions of the A-Minor Motif, the Most Common Motif in RNA Structure. Biochem. Mosc. 86, 952–961. 10.1134/S000629792108006X.

4. Doherty, E.A., Batey, R.T., Masquida, B., and Doudna, J.A. (2001). A universal mode of helix packing in RNA. Nat. Struct. Biol. 8, 339–343. 10.1038/86221.

5. Torres-Larios, A., Swinger, K.K., Pan, T., and Mondragón, A. (2006). Structure of ribonuclease P — a universal ribozyme. Curr. Opin. Struct. Biol. 16, 327–335. 10.1016/j.sbi.2006.04.002.

6. Chen, Y., and Varani, G. (2010). RNA Structure. In Encyclopedia of Life Sciences (John Wiley & Sons, Ltd). 10.1002/9780470015902.a0001339.pub2.

7. Mustoe, A.M., Brooks, C.L., and Al-Hashimi, H.M. (2014). Hierarchy of RNA Functional Dynamics. Annu. Rev. Biochem. 83, 441–466. 10.1146/annurev-biochem-060713-035524.

8. Baronti, L., Guzzetti, I., Ebrahimi, P., Friebe Sandoz, S., Steiner, E., Schlagnitweit, J., Fromm, B., Silva, L., Fontana, C., Chen, A.A., et al. (2020). Base-pair conformational switch modulates miR-34a targeting of Sirt1 mRNA. Nature 583, 139–144. 10.1038/s41586-020-2336-3.

9. Hansen, A.L., Nikolova, E.N., Casiano-Negroni, A., and Al-Hashimi, H.M. (2009). Extending the Range of Microsecond-to-Millisecond Chemical Exchange Detected in Labeled and Unlabeled Nucleic Acids by Selective Carbon R1ρ NMR Spectroscopy. J. Am. Chem. Soc. 131, 3818–3819. 10.1021/ja8091399.

10. Lee, J., Dethoff, E.A., and Al-Hashimi, H.M. (2014). Invisible RNA state dynamically couples distant motifs. Proc. Natl. Acad. Sci. U. S. A. 111, 9485–9490. 10.1073/pnas.1407969111.

11. Steiner, E., Schlagnitweit, J., Lundström, P., and Petzold, K. (2016). Capturing Excited States in the Fast-Intermediate Exchange Limit in Biological Systems Using 1H NMR Spectroscopy. Angew. Chem. Int. Ed. 55, 15869–15872. 10.1002/anie.201609102.

12. Schlagnitweit, J., Steiner, E., Karlsson, H., and Petzold, K. (2018). Efficient Detection of Structure and Dynamics in Unlabeled RNAs: The SELOPE Approach. Chem. Eur. J. 24, 6067–6070. 10.1002/chem.201800992.

13. Riad, M., Hopkins, N., Baronti, L., Karlsson, H., Schlagnitweit, J., and Petzold, K. (2021). Mutate-and-chemical-shift-fingerprint (MCSF) to characterize excited states in RNA using NMR spectroscopy. Nat. Protoc. 16, 5146–5170. 10.1038/s41596-021-00606-1.

14. Dasgupta, R., Steinmetzger, C., Ilgen, J., and Petzold, K. (2026). 1H R1ρ relaxation identifies a hidden intermediate in DNA base-pairing. Nat. Commun. 17, 4114. 10.1038/s41467-026-72559-6.

15. Ogle, J.M., Brodersen, D.E., Clemons, W.M., Tarry, M.J., Carter, A.P., and Ramakrishnan, V. (2001). Recognition of Cognate Transfer RNA by the 30S Ribosomal Subunit. Science 292, 897–902. 10.1126/science.1060612.

16. Lescoute, A., and Westhof, E. (2006). The A-minor motifs in the decoding recognition process. Biochimie 88, 993–999. 10.1016/j.biochi.2006.05.018.

17. Dethoff, E.A., Petzold, K., Chugh, J., Casiano-Negroni, A., and Al-Hashimi, H.M. (2012). Visualizing transient low-populated structures of RNA. Nature 491, 724–728. 10.1038/nature11498.

18. Rozov, A., Demeshkina, N., Westhof, E., Yusupov, M., and Yusupova, G. (2015). Structural insights into the translational infidelity mechanism. Nat. Commun. 6, 7251. 10.1038/ncomms8251.

19. Qin, D., Liu, Q., Devaraj, A., and Fredrick, K. (2012). Role of helix 44 of 16S rRNA in the fidelity of translation initiation. RNA 18, 485–495. 10.1261/rna.031203.111.

20. Tran, D.K., Finley, J., Vila-Sanjurjo, A., Lale, A., Sun, Q., and O’Connor, M. (2011). Tertiary interactions between helices h13 and h44 in 16S RNA contribute to the fidelity of translation. FEBS J. 278, 4405–4412. 10.1111/j.1742-4658.2011.08363.x.

21. Agrawal, R.K., Sharma, M.R., Kiel, M.C., Hirokawa, G., Booth, T.M., Spahn, C.M.T., Grassucci, R.A., Kaji, A., and Frank, J. (2004). Visualization of ribosome-recycling factor on the Escherichia coli 70S ribosome: Functional implications. Proc. Natl. Acad. Sci. 101, 8900–8905. 10.1073/pnas.0401904101.

22. Moazed, D., and Noller, H.F. (1987). Interaction of antibiotics with functional sites in 16S ribosomal RNA. Nature 327, 389–394. 10.1038/327389a0.

23. Kaul, M., Barbieri, C.M., and Pilch, D.S. (2005). Defining the basis for the specificity of aminoglycoside-rRNA recognition: a comparative study of drug binding to the A sites of Escherichia coli and human rRNA. J. Mol. Biol. 346, 119–134. 10.1016/j.jmb.2004.11.041.

24. O’Sullivan, M.E., Perez, A., Lin, R., Sajjadi, A., Ricci, A.J., and Cheng, A.G. (2017). Towards the Prevention of Aminoglycoside-Related Hearing Loss. Front. Cell. Neurosci. 11. 10.3389/fncel.2017.00325.

25. Panecka, J., Šponer, J., and Trylska, J. (2015). Conformational dynamics of bacterial and human cytoplasmic models of the ribosomal A-site. Biochimie 112, 96–110. 10.1016/j.biochi.2015.02.021.

26. Sabbavarapu, N.M., Pieńko, T., Zalman, B.-H., Trylska, J., and Baasov, T. (2018). Exploring eukaryotic versus prokaryotic ribosomal RNA recognition with aminoglycoside derivatives. MedChemComm 9, 503–508. 10.1039/C8MD00001H.

27. Karlsson, H., Baronti, L., and Petzold, K. (2020). A robust and versatile method for production and purification of large-scale RNA samples for structural biology. RNA 26, 1023–1037. 10.1261/rna.075697.120.

28. Fürtig, B., Richter, C., Wöhnert, J., and Schwalbe, H. (2003). NMR Spectroscopy of RNA. ChemBioChem 4, 936–962. 10.1002/cbic.200300700.

29. Watson, Z.L., Ward, F.R., Méheust, R., Ad, O., Schepartz, A., Banfield, J.F., and Cate, J.H. (2020). Structure of the bacterial ribosome at 2 Å resolution. eLife 9, e60482. 10.7554/eLife.60482.

30. Shu, Z., and Bevilacqua, P.C. (1999). Isolation and Characterization of Thermodynamically Stable and Unstable RNA Hairpins from a Triloop Combinatorial Library. Biochemistry 38, 15369–15379. 10.1021/bi991774z.

31. Xin, Y., Laing, C., Leontis, N.B., and Schlick, T. (2008). Annotation of tertiary interactions in RNA structures reveals variations and correlations. RNA 14, 2465–2477. 10.1261/rna.1249208.

32. Petrov, A.I., Zirbel, C.L., and Leontis, N.B. (2011). WebFR3D--a server for finding, aligning and analyzing recurrent RNA 3D motifs. Nucleic Acids Res. 39, W50–W55. 10.1093/nar/gkr249.

33. Dallaire, P., and Major, F. (2016). Exploring Alternative RNA Structure Sets Using MC-Flashfold and db2cm. Methods Mol. Biol. Clifton NJ 1490, 237–251. 10.1007/978-1-4939-6433-8_15.

34. Gromadski, K.B., Daviter, T., and Rodnina, M.V. (2006). A Uniform Response to Mismatches in Codon-Anticodon Complexes Ensures Ribosomal Fidelity. Mol. Cell 21, 369–377. 10.1016/j.molcel.2005.12.018.

35. Romanowska, J., McCammon, J.A., and Trylska, J. (2011). Understanding the Origins of Bacterial Resistance to Aminoglycosides through Molecular Dynamics Mutational Study of the Ribosomal A-Site. PLOS Comput. Biol. 7, e1002099. 10.1371/journal.pcbi.1002099.

36. Pulk, A., Maiväli, Ü., and Remme, J. (2006). Identification of nucleotides in E. coli 16S rRNA essential for ribosome subunit association. RNA 12, 790–796. 10.1261/rna.2275906.

37. Pfister, P., Hobbie, S., Vicens, Q., Böttger, E.C., and Westhof, E. (2003). The Molecular Basis for A-Site Mutations Conferring Aminoglycoside Resistance: Relationship between Ribosomal Susceptibility and X-ray Crystal Structures. ChemBioChem 4, 1078–1088. 10.1002/cbic.200300657.

38. Kofman, C., Watkins, A.M., Kim, D.S., Willi, J.A., Wooldredge, A.C., Karim, A.S., Das, R., and Jewett, M.C. (2022). Computationally-guided design and selection of high performing ribosomal active site mutants. Nucleic Acids Res. 50, 13143–13154. 10.1093/nar/gkac1036.

39. McGinnis, J.L., and Weeks, K.M. (2014). Ribosome RNA Assembly Intermediates Visualized in Living Cells. Biochemistry 53, 3237–3247. 10.1021/bi500198b.

40. Xu, Z., and Culver, G.M. (2010). Differential assembly of 16S rRNA domains during 30S subunit formation. RNA 16, 1990–2001. 10.1261/rna.2246710.

41. Sharma, I.M., and Woodson, S.A. (2020). RbfA and IF3 couple ribosome biogenesis and translation initiation to increase stress tolerance. Nucleic Acids Res. 48, 359–372. 10.1093/nar/gkz1065.

42. Jomaa, A., Stewart, G., Martín-Benito, J., Zielke, R., Campbell, T.L., Maddock, J.R., Brown, E.D., and Ortega, J. (2011). Understanding ribosome assembly: the structure of in vivo assembled immature 30S subunits revealed by cryo-electron microscopy. RNA 17, 697–709. 10.1261/rna.2509811.

43. McGinnis, J.L., Liu, Q., Lavender, C.A., Devaraj, A., McClory, S.P., Fredrick, K., and Weeks, K.M. (2015). In-cell SHAPE reveals that free 30S ribosome subunits are in the inactive state. Proc. Natl. Acad. Sci. U. S. A. 112, 2425–2430. 10.1073/pnas.1411514112.

44. Jahagirdar, D., Jha, V., Basu, K., Gomez-Blanco, J., Vargas, J., and Ortega, J. (2020). Alternative conformations and motions adopted by 30S ribosomal subunits visualized by cryo-electron microscopy. RNA 26, 2017–2030. 10.1261/rna.075846.120.

45. Sun, J., Kinman, L.F., Jahagirdar, D., Ortega, J., and Davis, J.H. (2023). KsgA facilitates ribosomal small subunit maturation by proofreading a key structural lesion. Nat. Struct. Mol. Biol. 30, 1468–1480. 10.1038/s41594-023-01078-5.

46. May, M.B., Lopez-Perez, G.S., and Davis, J.H. (2026). Capturing ribosomal structures in cellular extracts with cryoPRISM: A purification-free cryoEM approach reveals novel structural states. Proc. Natl. Acad. Sci. 123, e2521210123. 10.1073/pnas.2521210123.

47. Leontis, N.B., and Westhof, E. (2001). Geometric nomenclature and classification of RNA base pairs. RNA 7, 499–512. 10.1017/S1355838201002515.

48. Baulin, E., Yacovlev, V., Khachko, D., Spirin, S., and Roytberg, M. (2016). URS DataBase: universe of RNA structures and their motifs. Database J. Biol. Databases Curation 2016, baw085. 10.1093/database/baw085.

49. Schedlbauer, A., Iturrioz, I., Ochoa-Lizarralde, B., Diercks, T., López-Alonso, J.P., Lavin, J.L., Kaminishi, T., Çapuni, R., Dhimole, N., de Astigarraga, E., et al. (2021). A conserved rRNA switch is central to decoding site maturation on the small ribosomal subunit. Sci. Adv. 7, eabf7547. 10.1126/sciadv.abf7547.

50. Karlsson, H., Feyrer, H., Baronti, L., and Petzold, K. (2021). Production of Structured RNA Fragments by In Vitro Transcription and HPLC Purification. Curr. Protoc. 1, e159. 10.1002/cpz1.159.

51. Wishart, D.S., Bigam, C.G., Yao, J., Abildgaard, F., Dyson, H.J., Oldfield, E., Markley, J.L., and Sykes, B.D. (1995). 1H, 13C and 15N chemical shift referencing in biomolecular NMR. J. Biomol. NMR 6, 135–140. 10.1007/BF00211777.

52. Skinner, S.P., Fogh, R.H., Boucher, W., Ragan, T.J., Mureddu, L.G., and Vuister, G.W. (2016). CcpNmr AnalysisAssign: a flexible platform for integrated NMR analysis. J. Biomol. NMR 66, 111–124. 10.1007/s10858-016-0060-y.

53. Davis, D.G., Perlman, M.E., and London, R.E. (1994). Direct Measurements of the Dissociation-Rate Constant for Inhibitor-Enzyme Complexes via the *T*1ρ and *T*2 (CPMG) Methods. J. Magn. Reson. B 104, 266–275. 10.1006/jmrb.1994.1084.

54. Trott, O., and Palmer III, A.G. (2004). Theoretical study of R1ρ rotating-frame and R2 free-precession relaxation in the presence of n-site chemical exchange. J. Magn. Reson. 170, 104–112. 10.1016/j.jmr.2004.06.005.

55. Miloushev, V.Z., and Palmer, A.G. (2005). R1ρ relaxation for two-site chemical exchange: General approximations and some exact solutions. J. Magn. Reson. 177, 221–227. 10.1016/j.jmr.2005.07.023.

56. Bothe, J.R., Stein, Z.W., and Al-Hashimi, H.M. (2014). Evaluating the uncertainty in exchange parameters determined from off-resonance R1ρ relaxation dispersion for systems in fast exchange. J. Magn. Reson. 244, 18–29. 10.1016/j.jmr.2014.04.010.

57. Kimsey, I.J., Petzold, K., Sathyamoorthy, B., Stein, Z.W., and Al-Hashimi, H.M. (2015). Visualizing transient Watson–Crick-like mispairs in DNA and RNA duplexes. Nature 519, 315–320. 10.1038/nature14227.

58. Orelle, C., Carlson, E.D., Szal, T., Florin, T., Jewett, M.C., and Mankin, A.S. (2015). Protein synthesis by ribosomes with tethered subunits. Nature 524, 119–124. 10.1038/nature14862.

59. Asai, T., Zaporojets, D., Squires, C., and Squires, C.L. (1999). An Escherichia coli strain with all chromosomal rRNA operons inactivated: complete exchange of rRNA genes between bacteria. Proc. Natl. Acad. Sci. U. S. A. 96, 1971–1976. 10.1073/pnas.96.5.1971.

60. Punjani, A., Rubinstein, J.L., Fleet, D.J., and Brubaker, M.A. (2017). cryoSPARC: algorithms for rapid unsupervised cryo-EM structure determination. Nat. Methods 14, 290–296. 10.1038/nmeth.4169.

61. Meng, E.C., Goddard, T.D., Pettersen, E.F., Couch, G.S., Pearson, Z.J., Morris, J.H., and Ferrin, T.E. (2023). UCSF ChimeraX: Tools for structure building and analysis. Protein Sci. 32, e4792. 10.1002/pro.4792.

62. Punjani, A., and Fleet, D.J. (2021). 3D variability analysis: Resolving continuous flexibility and discrete heterogeneity from single particle cryo-EM. J. Struct. Biol. 213, 107702. 10.1016/j.jsb.2021.107702.

63. Koller, T.O., Berger, M.J., Morici, M., Paternoga, H., Bulatov, T., Di Stasi, A., Dang, T., Mainz, A., Raulf, K., Crowe-McAuliffe, C., et al. (2024). Paenilamicins are context-specific translocation inhibitors of protein synthesis. Nat. Chem. Biol. 20, 1691–1700. 10.1038/s41589-024-01752-9.

64. Noeske, J., Wasserman, M.R., Terry, D.S., Altman, R.B., Blanchard, S.C., and Cate, J.H.D. (2015). High-resolution structure of the Escherichia coli ribosome. Nat. Struct. Mol. Biol. 22, 336–341. 10.1038/nsmb.2994.

65. Borovinskaya, M.A., Shoji, S., Holton, J.M., Fredrick, K., and Cate, J.H.D. (2007). A Steric Block in Translation Caused by the Antibiotic Spectinomycin. ACS Chem. Biol. 2, 545–552. 10.1021/cb700100n.

66. Yamashita, K., Palmer, C.M., Burnley, T., and Murshudov, G.N. (2021). Cryo-EM single-particle structure refinement and map calculation using Servalcat. Acta Crystallogr. Sect. Struct. Biol. 77, 1282–1291. 10.1107/S2059798321009475.

67. Yamashita, K., Wojdyr, M., Long, F., Nicholls, R.A., and Murshudov, G.N. (2023). GEMMI and Servalcat restrain REFMAC5. Acta Crystallogr. Sect. Struct. Biol. 79, 368–373. 10.1107/S2059798323002413.

68. Emsley, P., Lohkamp, B., Scott, W.G., and Cowtan, K. (2010). Features and development of *Coot*. Acta Crystallogr. D Biol. Crystallogr. 66, 486–501. 10.1107/S0907444910007493.

69. Kimura, F., Harada, F., and Nishimura, S. (1971). Primary sequence of tRNA-Val-1 from Escherichia coli B. II. Isolation of large fragments by limited digestion with RNases, and overlapping of fragments to reduce the total primary sequence. Biochemistry 10, 3277–3283. 10.1021/bi00793a018.

70. Dube, S.K., and Marcker, K.A. (1969). The Nucleotide Sequence of N-Formyl-Methionyl-Transfer RNA. Eur. J. Biochem. 8, 256–262. 10.1111/j.1432-1033.1969.tb00522.x.

71. Watson, Z.L., Knudson, I.J., Ward, F.R., Miller, S.J., Cate, J.H.D., Schepartz, A., and Abramyan, A.M. (2023). Atomistic simulations of the Escherichia coli ribosome provide selection criteria for translationally active substrates. Nat. Chem. 15, 913–921. 10.1038/s41557-023-01226-w.

72. Leonarski, F., Henning-Knechtel, A., Kirmizialtin, S., Ennifar, E., and Auffinger, P. (2025). Principles of ion binding to RNA inferred from the analysis of a 1.55 Å resolution bacterial ribosome structure – Part I: Mg2+. Nucleic Acids Res. 53, gkae1148. 10.1093/nar/gkae1148.

73. Davis, I.W., Leaver-Fay, A., Chen, V.B., Block, J.N., Kapral, G.J., Wang, X., Murray, L.W., Arendall, W.B., III, Snoeyink, J., Richardson, J.S., et al. (2007). MolProbity: all-atom contacts and structure validation for proteins and nucleic acids. Nucleic Acids Res. 35, W375–W383. 10.1093/nar/gkm216.

74. Li, L., Rybak, M.Y., Lin, J., and Gagnon, M.G. (2024). The ribosome termination complex remodels release factor RF3 and ejects GDP. Nat. Struct. Mol. Biol. 31, 1909–1920. 10.1038/s41594-024-01360-0.

75. Johnson, P.Z., and Simon, A.E. (2023). RNAcanvas: interactive drawing and exploration of nucleic acid structures. Nucleic Acids Res. 51, W501–W508. 10.1093/nar/gkad302.

