## Supplemental Information for "Universal 3D Motif dynamics in RNA: The A-minor Switch"

### 1. Supplemental Figures

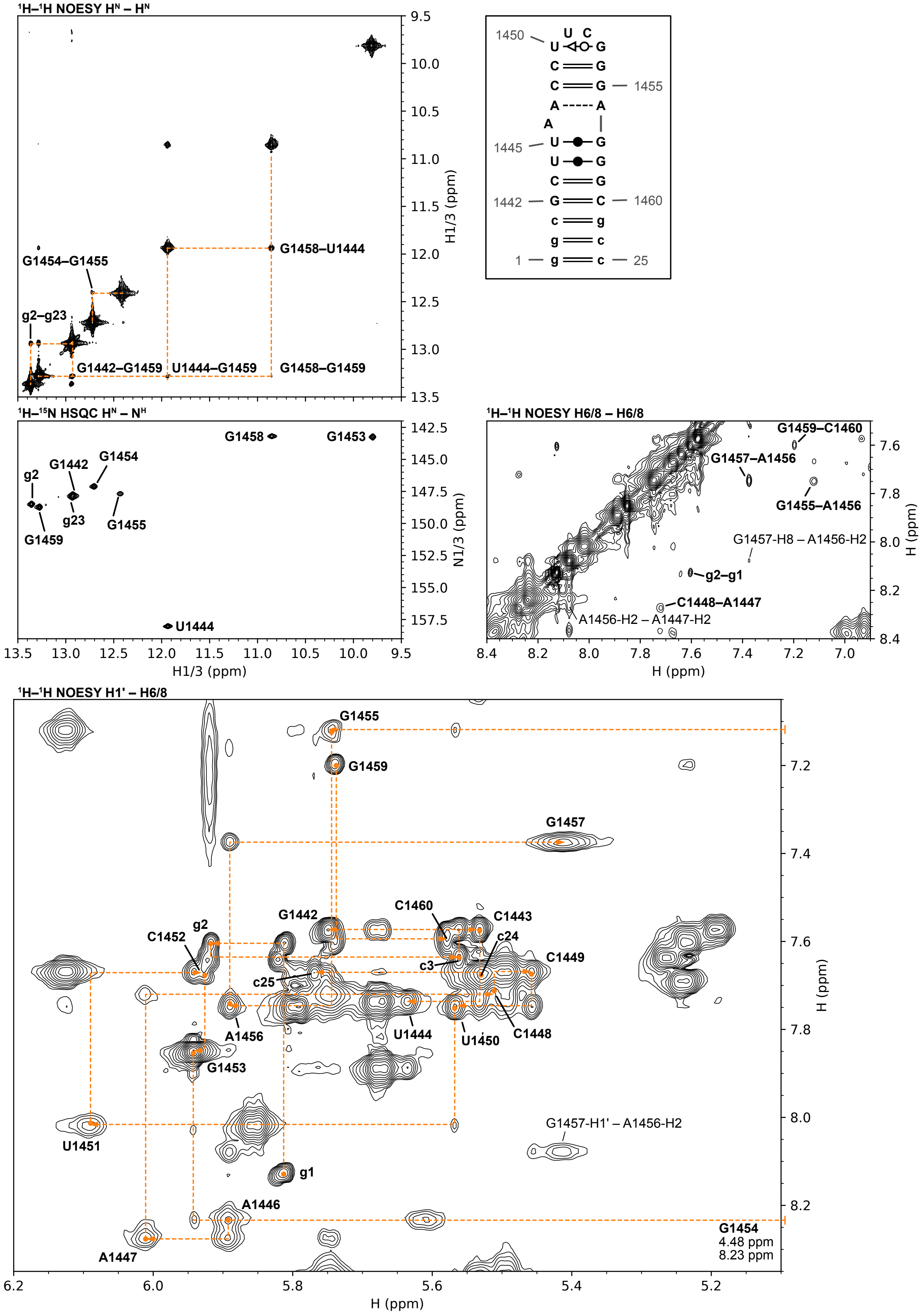

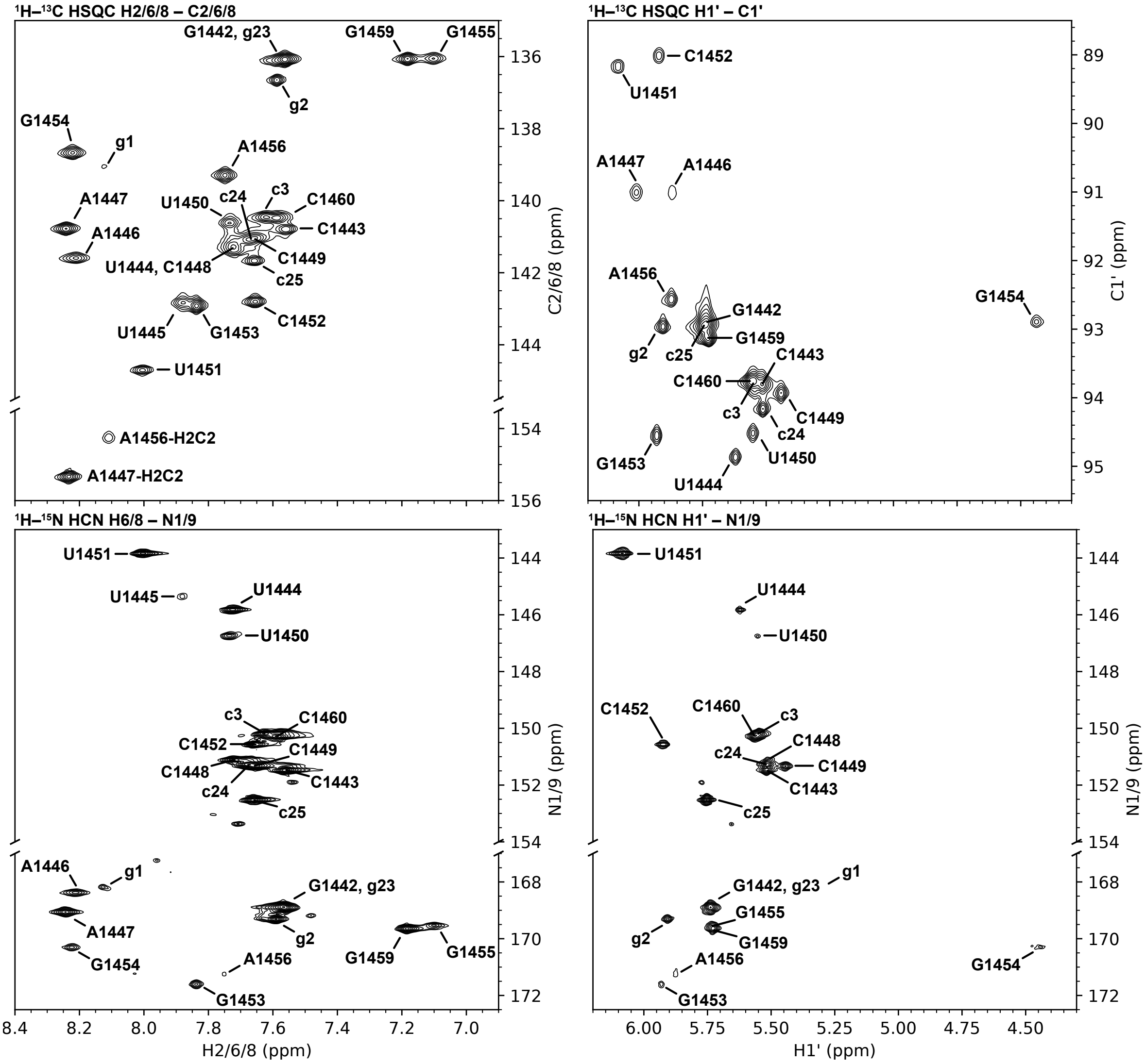


**Figure S1. Chemical shift assignments of the 25 nt h44-top$^{WT}$ hairpin construct.** Conditions: 15 mM NaP, 25 mM NaCl, 0.5 mM EDTA, pH 6.5, 90% $H_2O$/10% $D_2O$, 298 K. An unlabeled RNA sample was used to acquire the $^1H$–$^1H$ NOESY spectrum, while a $^{13}C$,$^{15}N$-labeled sample was used to acquire all other spectra. Sequential imino–imino and anomeric–aromatic correlations are indicated by dashed lines in the different regions of the NOESY spectrum.

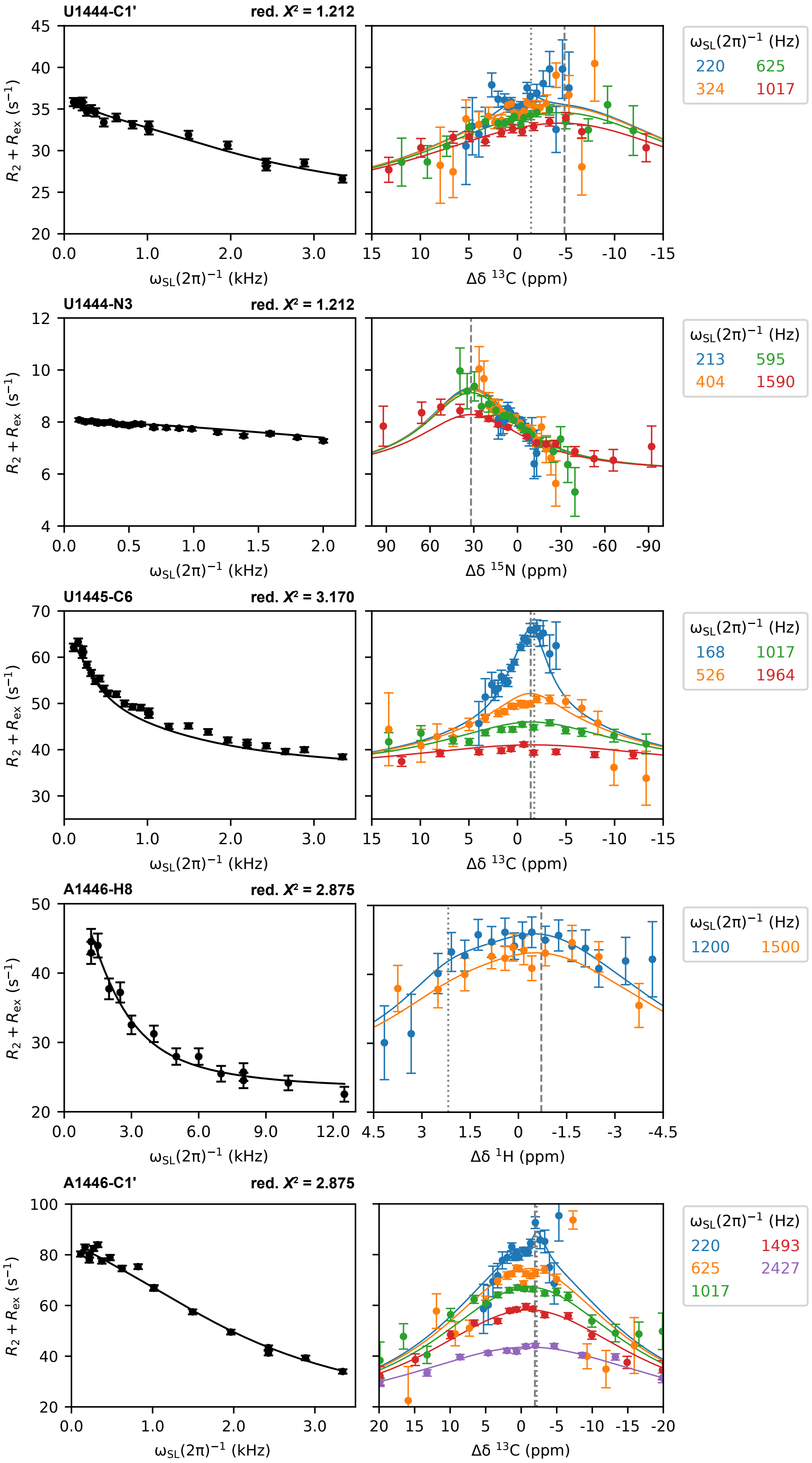
U1444-C1'
red. X² = 1.212
U1444-N3
red. X² = 1.212
U1445-C6
red. X² = 3.170
A1446-H8
red. X² = 2.875
A1446-C1'
red. X² = 2.875
R2 + Rex (s−1)
ωSL(2π)−1 (kHz)
Δδ 13C (ppm)
Δδ 15N (ppm)
Δδ 1H (ppm)
ωSL(2π)−1 (Hz)
220 625 324 1017
213 595 404 1590
168 1017 526 1964
1200 1500
220 1493 625 2427 1017

1

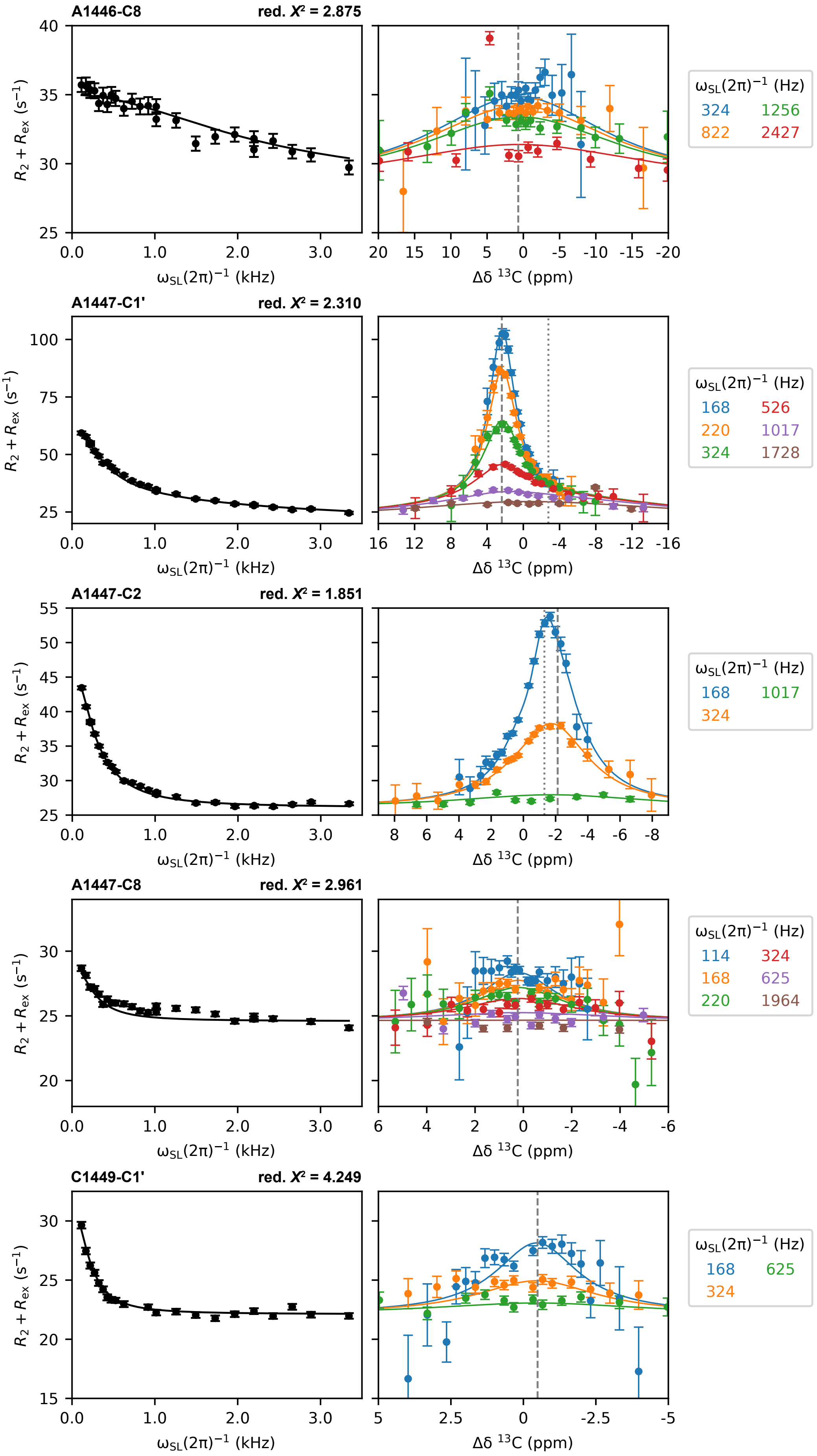

A1446-C8
red. X² = 2.875
A1447-C1'
red. X² = 2.310
A1447-C2
red. X² = 1.851
A1447-C8
red. X² = 2.961
C1449-C1'
red. X² = 4.249
R2 + Rex (s−1)
ωSL(2π)−1 (kHz)
Δδ 13C (ppm)
ωSL(2π)−1 (Hz)
324 1256
822 2427
168 526
220 1017
324 1728
168 1017
324
114 324
168 625
220 1964
168 625
324

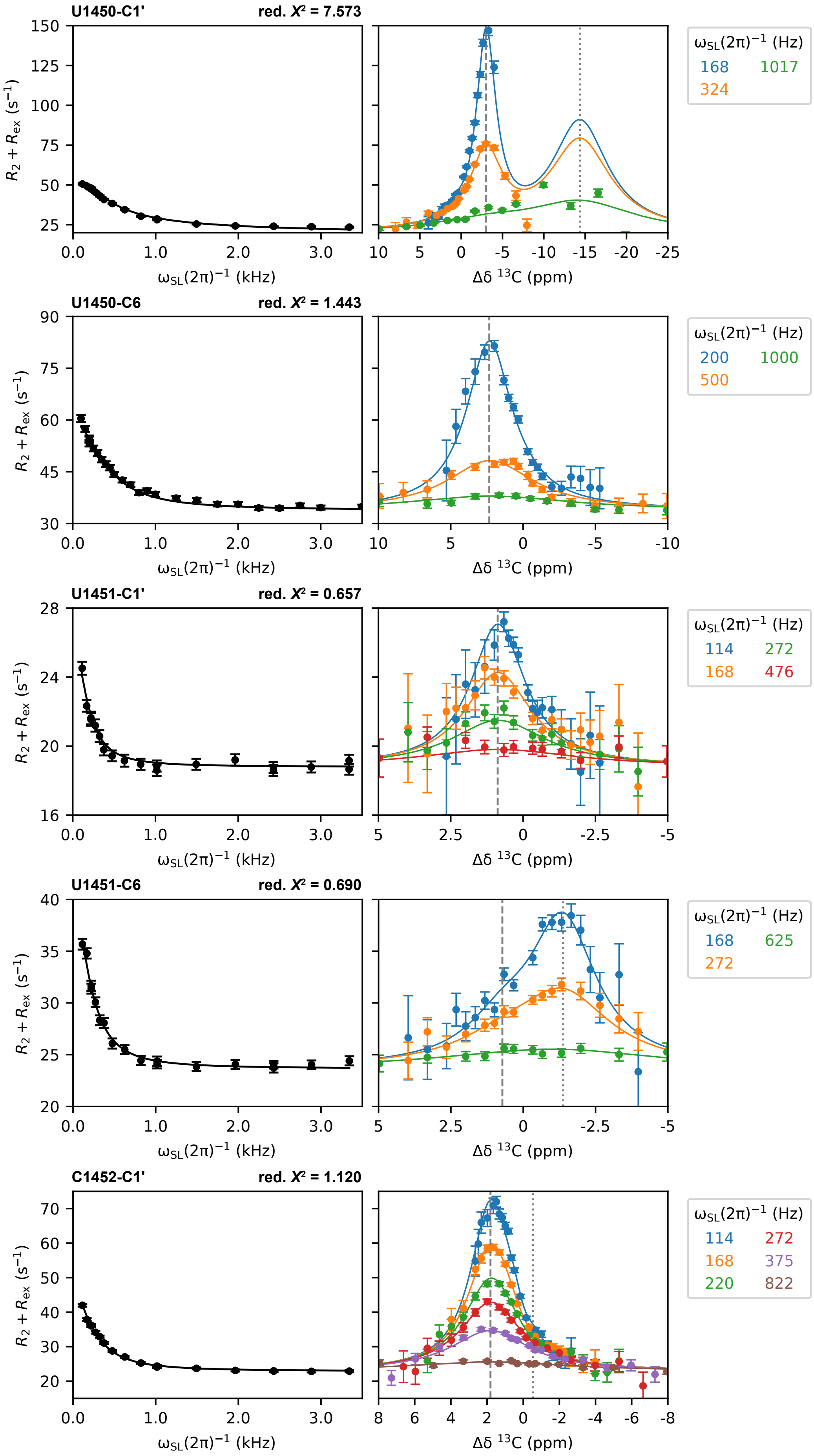
U1450-C1'
red. X² = 7.573
U1450-C6
red. X² = 1.443
U1451-C1'
red. X² = 0.657
U1451-C6
red. X² = 0.690
C1452-C1'
red. X² = 1.120
$R_2 + R_{ex}$ (s$^{-1}$)
$\omega_{SL}(2\pi)^{-1}$ (kHz)
Δδ $^{13}$C (ppm)
$\omega_{SL}(2\pi)^{-1}$ (Hz)
168 1017 324
200 1000 500
114 272 168 476
168 625 272
114 272 168 375 220 822

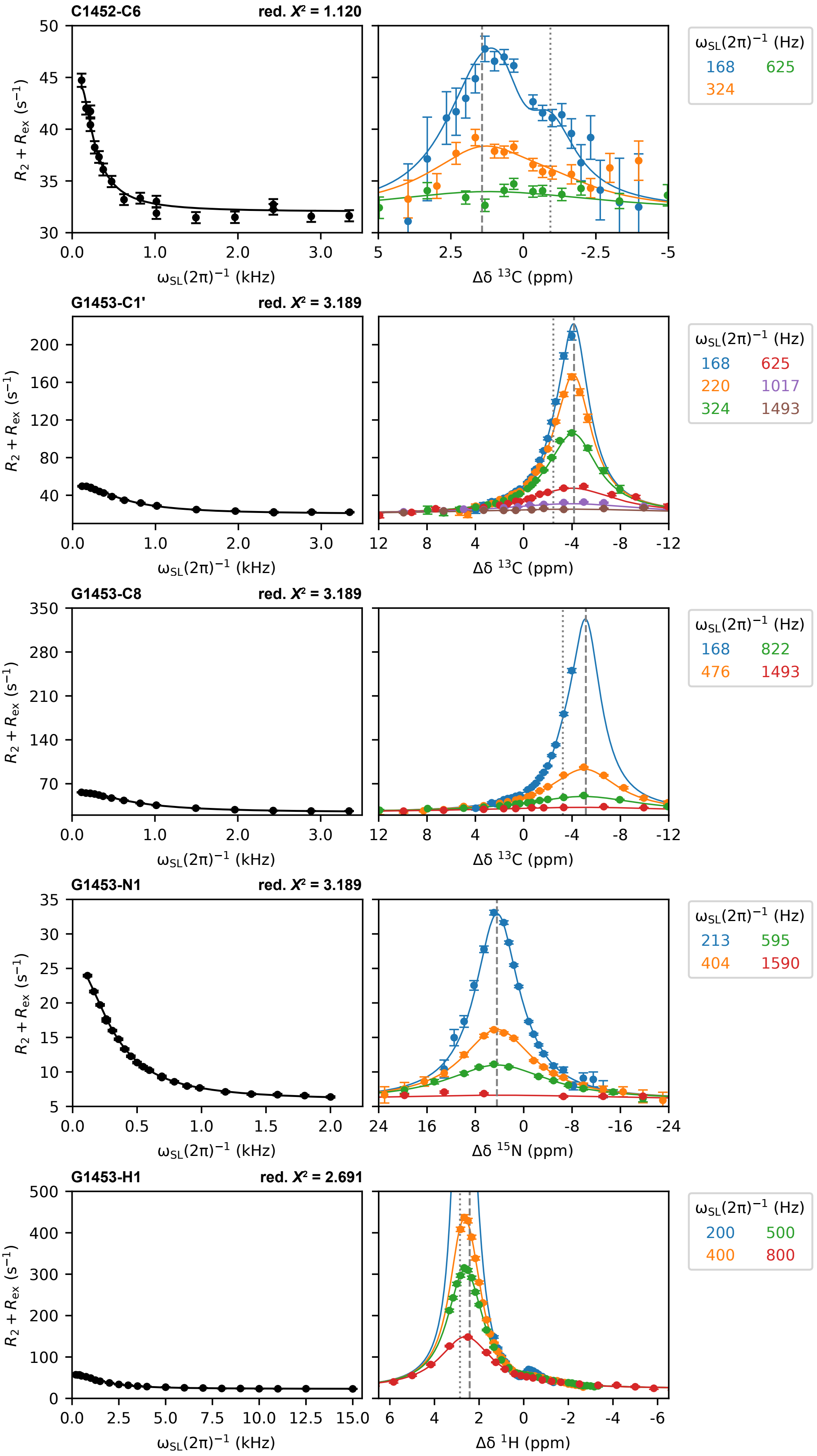
C1452-C6
red. X² = 1.120
G1453-C1'
red. X² = 3.189
G1453-C8
red. X² = 3.189
G1453-N1
red. X² = 3.189
G1453-H1
red. X² = 2.691
R2 + Rex (s−1)
ωSL(2π)−1 (kHz)
Δδ 13C (ppm)
Δδ 15N (ppm)
Δδ 1H (ppm)
ωSL(2π)−1 (Hz)
168 625
324
168 625
220 1017
324 1493
168 822
476 1493
213 595
404 1590
200 500
400 800

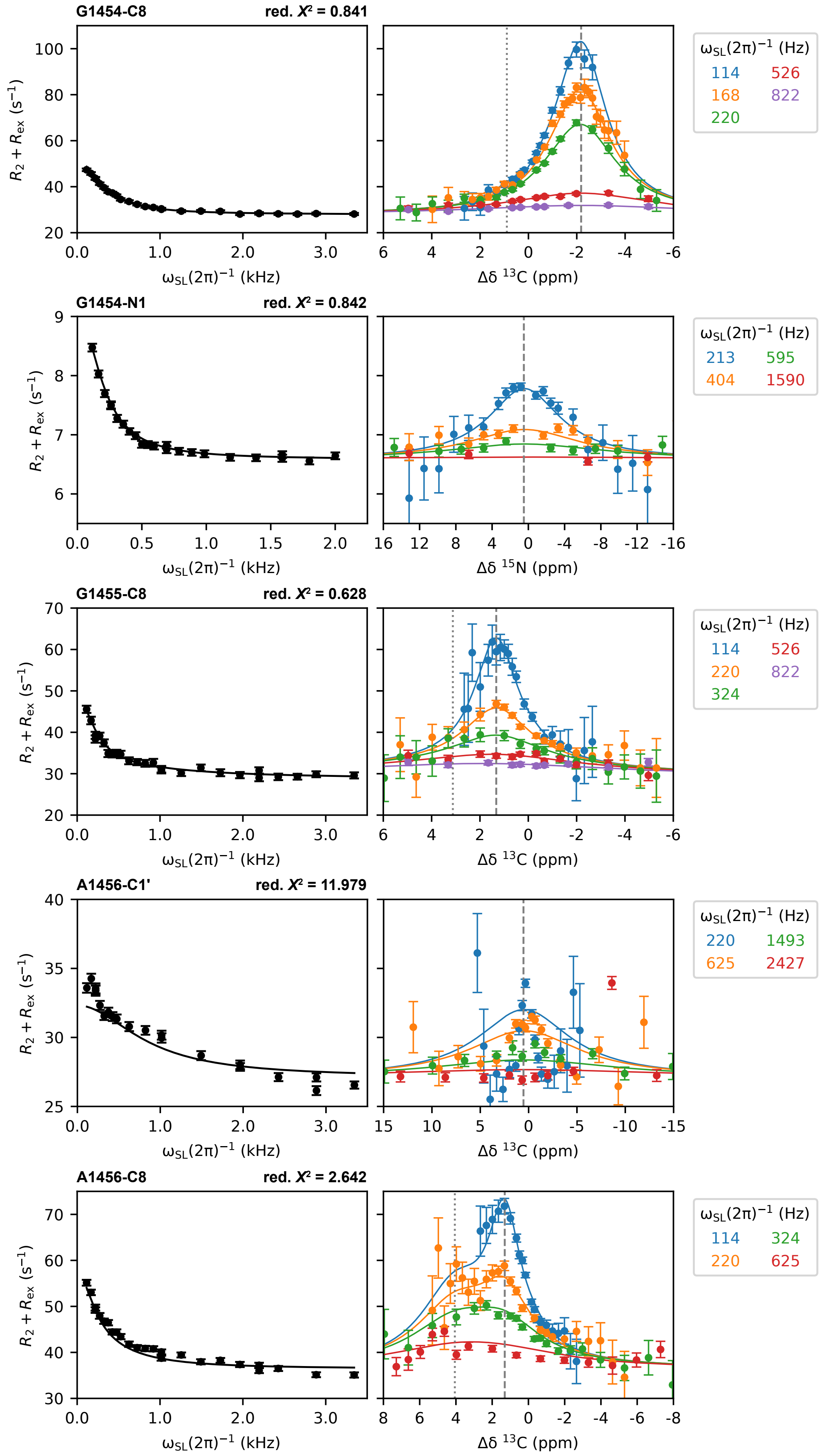

G1454-C8
red. $X^2$ = 0.841
$R_2 + R_{ex}$ ($s^{-1}$)
$\omega_{SL}(2\pi)^{-1}$ (kHz)
Δδ $^{13}$C (ppm)
$\omega_{SL}(2\pi)^{-1}$ (Hz)
114
526
168
822
220
G1454-N1
red. $X^2$ = 0.842
$R_2 + R_{ex}$ ($s^{-1}$)
$\omega_{SL}(2\pi)^{-1}$ (kHz)
Δδ $^{15}$N (ppm)
$\omega_{SL}(2\pi)^{-1}$ (Hz)
213
595
404
1590
G1455-C8
red. $X^2$ = 0.628
$R_2 + R_{ex}$ ($s^{-1}$)
$\omega_{SL}(2\pi)^{-1}$ (kHz)
Δδ $^{13}$C (ppm)
$\omega_{SL}(2\pi)^{-1}$ (Hz)
114
526
220
822
324
A1456-C1'
red. $X^2$ = 11.979
$R_2 + R_{ex}$ ($s^{-1}$)
$\omega_{SL}(2\pi)^{-1}$ (kHz)
Δδ $^{13}$C (ppm)
$\omega_{SL}(2\pi)^{-1}$ (Hz)
220
1493
625
2427
A1456-C8
red. $X^2$ = 2.642
$R_2 + R_{ex}$ ($s^{-1}$)
$\omega_{SL}(2\pi)^{-1}$ (kHz)
Δδ $^{13}$C (ppm)
$\omega_{SL}(2\pi)^{-1}$ (Hz)
114
324
220
625

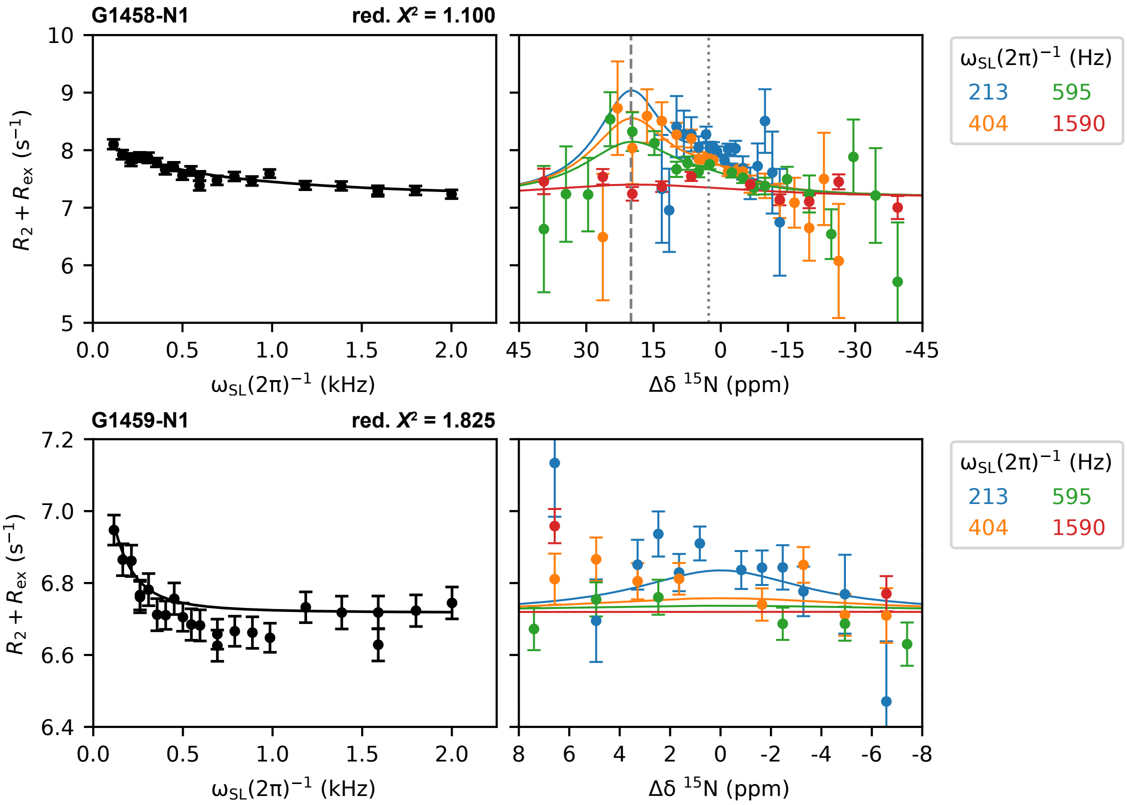


**Figure S2. On- and off-resonance $^1$H, $^{13}$C and $^{15}$N relaxation dispersion profiles for h44-top$^{WT}$.** In the off-resonance plots, Δδ = 0 ppm corresponds to the position of the observable average resonance. Chemical shift changes from that to the excited state(s) are indicated with dashed (from $\delta\omega_{ab}$) and dotted (from $\delta\omega_{ac}$) lines. Parameters for each dataset are given in **Table S2**. Source data are found in **Data S2**.

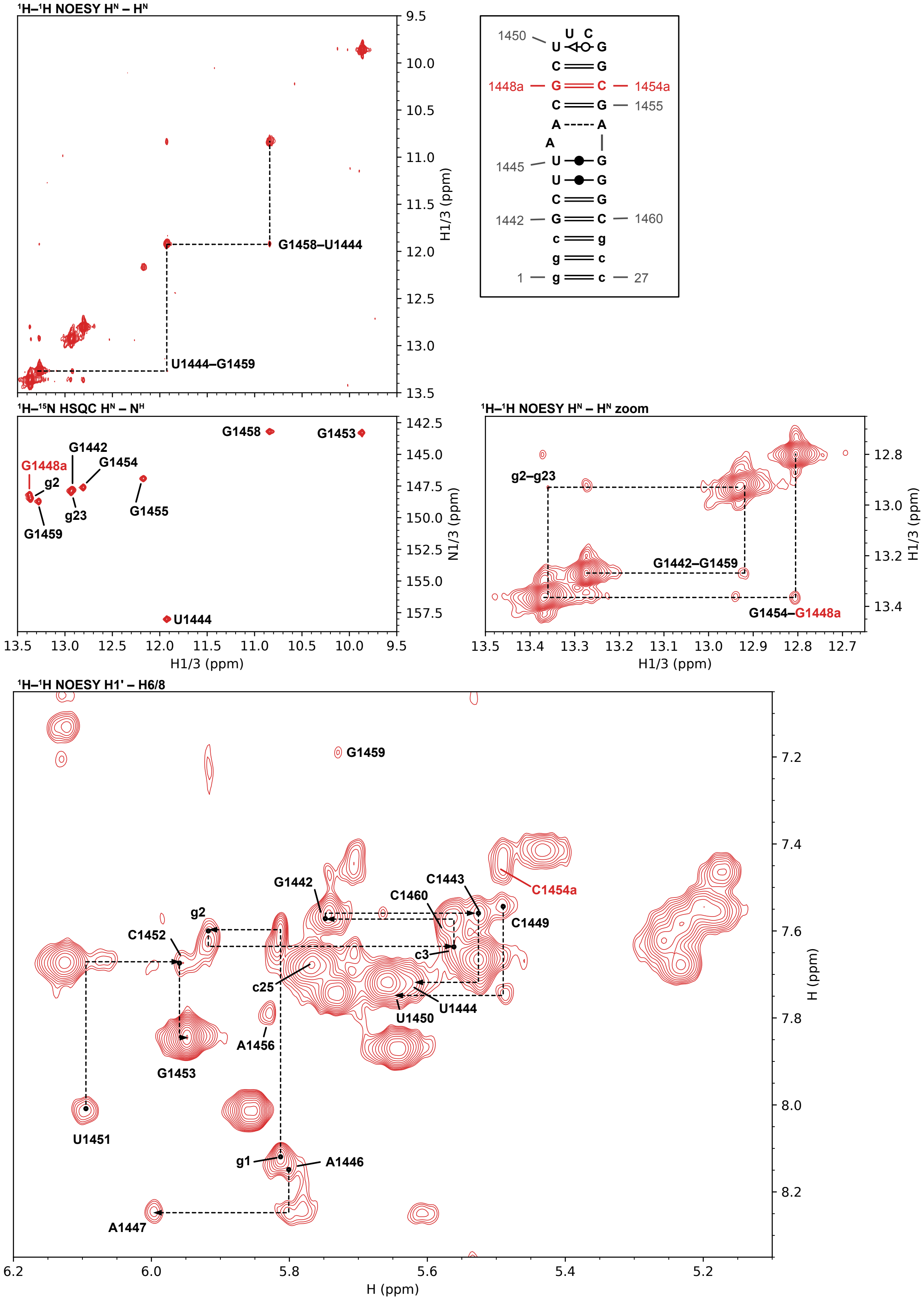

¹H–¹H NOESY Hᴺ – Hᴺ
G1458–U1444
U1444–G1459
H1/3 (ppm)
1450
1448a
1454a
1455
1445
1442
1460
1
27
¹H–¹⁵N HSQC Hᴺ – Nᴴ
G1458
G1453
G1442
G1448a
G1454
g2
g23
G1455
G1459
U1444
N1/3 (ppm)
H1/3 (ppm)
¹H–¹H NOESY Hᴺ – Hᴺ zoom
g2–g23
G1442–G1459
G1454–G1448a
¹H–¹H NOESY H1' – H6/8
G1459
C1454a
C1443
C1460
G1442
g2
C1449
C1452
c3
c25
U1444
U1450
A1456
G1453
U1451
g1
A1446
A1447
H (ppm)


1

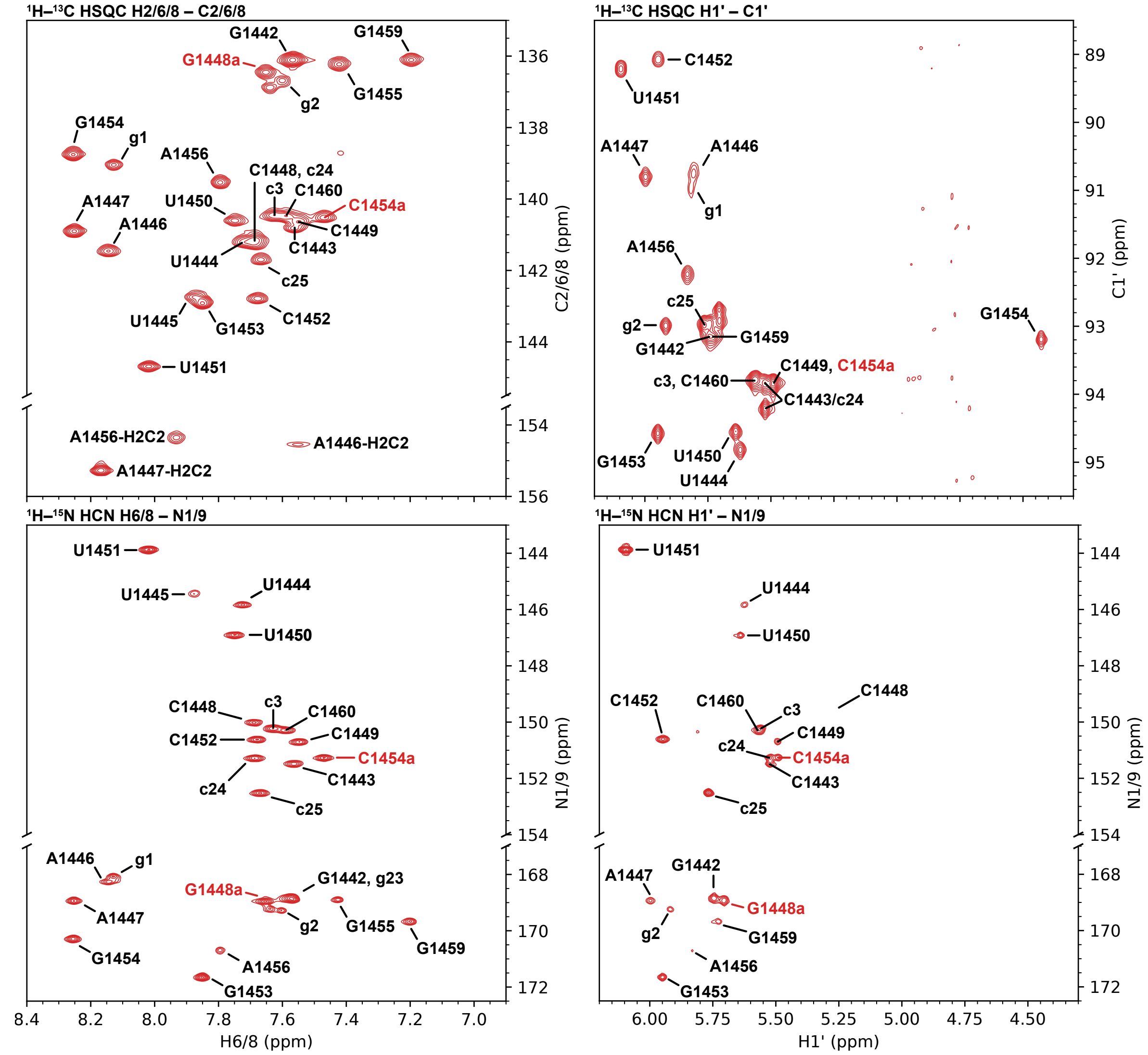


**Figure S3. Chemical shift assignments of the 27 nt h44-top$^{GS}$ hairpin construct.** Conditions: 15 mM NaP, 25 mM NaCl, 0.5 mM EDTA, pH 6.5, 90% $H_2O$/10% $D_2O$, 298 K. All spectra were acquired on a $^{13}C$,$^{15}N$-labeled sample. Sequential imino–imino (with a magnified view from 12.65 to 13.50 ppm) and anomeric–aromatic correlations are indicated by dashed lines in the different regions of the NOESY spectrum.

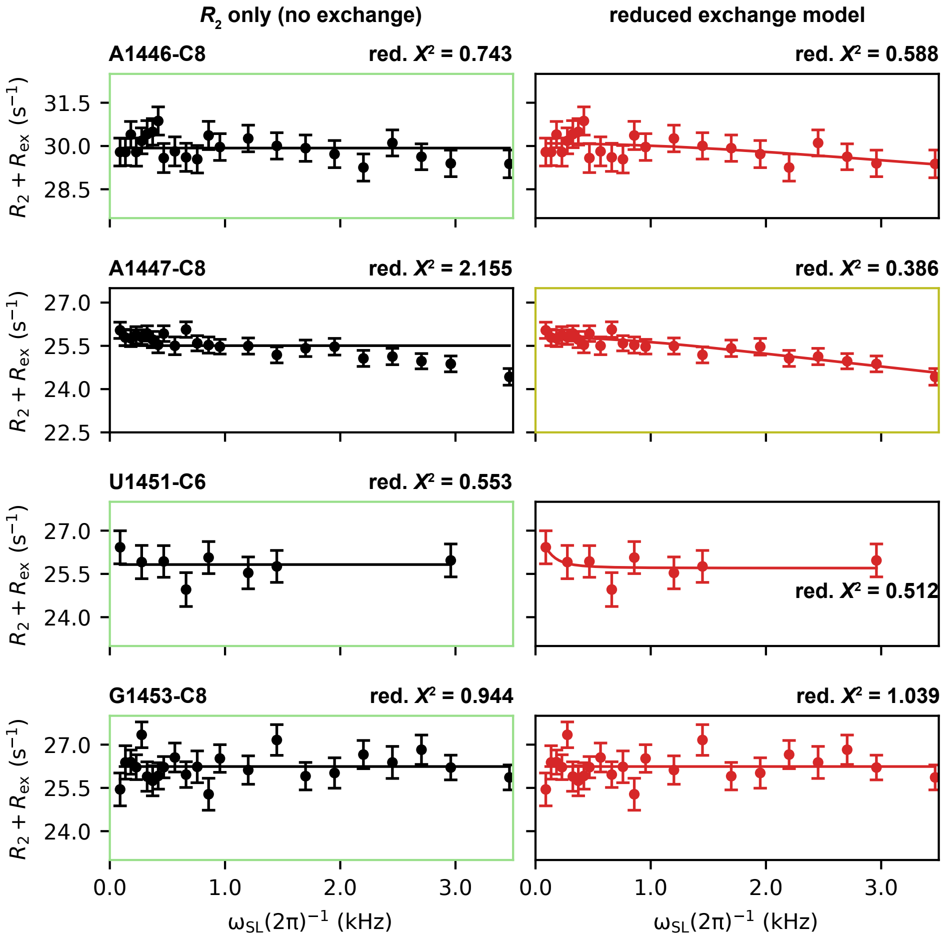


**Figure S4. On-resonance $^{13}C$ relaxation dispersion profiles for h44-top$^{GS}$.** No-exchange ($R_2$ only, left column) and reduced exchange (right column) models were tested against each other; the preferred model for each atom based on statistical criteria is shown with a shaded border. Clear evidence for residual exchange is only found for A1447-C8, albeit with high parameter uncertainties (yellow border). Parameters for each dataset are given in **Table S4**. Source data are found in **Data S3**.

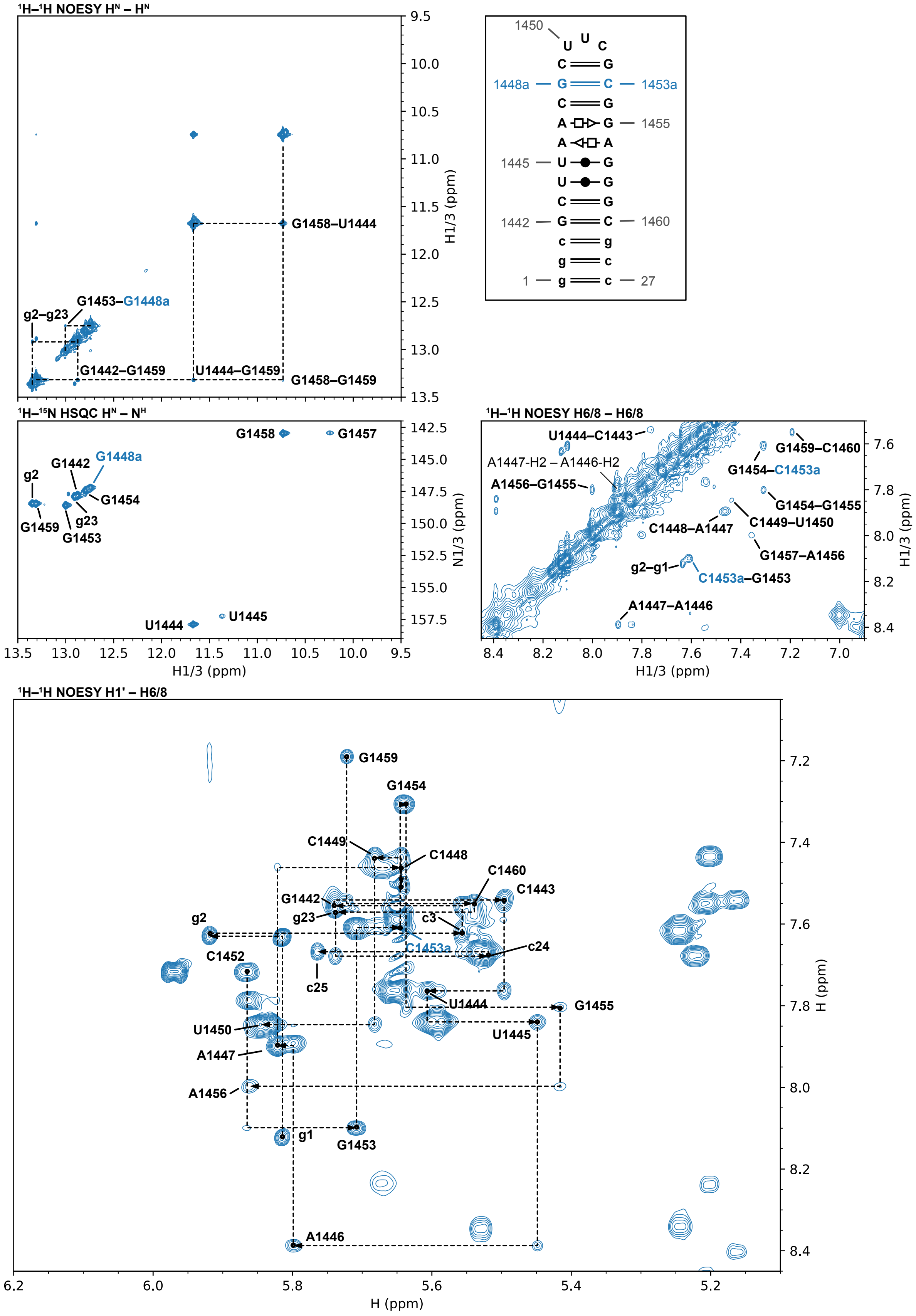

¹H–¹H NOESY Hᴺ – Hᴺ
G1458–U1444
G1453–G1448a
g2–g23
G1442–G1459
U1444–G1459
G1458–G1459
H1/3 (ppm)
1450
U U C
1448a
1453a
1455
1445
1442
1460
1
27
¹H–¹⁵N HSQC Hᴺ – Nᴴ
G1458
G1457
G1442
G1448a
g2
G1454
G1459
g23
G1453
U1444
U1445
N1/3 (ppm)
H1/3 (ppm)
¹H–¹H NOESY H6/8 – H6/8
U1444–C1443
G1459–C1460
A1447-H2 – A1446-H2
G1454–C1453a
A1456–G1455
G1454–G1455
C1449–U1450
C1448–A1447
G1457–A1456
g2–g1
C1453a–G1453
A1447–A1446
¹H–¹H NOESY H1' – H6/8
G1459
G1454
C1449
C1448
C1460
C1443
G1442
g23
c3
g2
C1453a
c24
C1452
c25
U1444
G1455
U1450
U1445
A1447
A1456
g1
G1453
A1446
H (ppm)


1

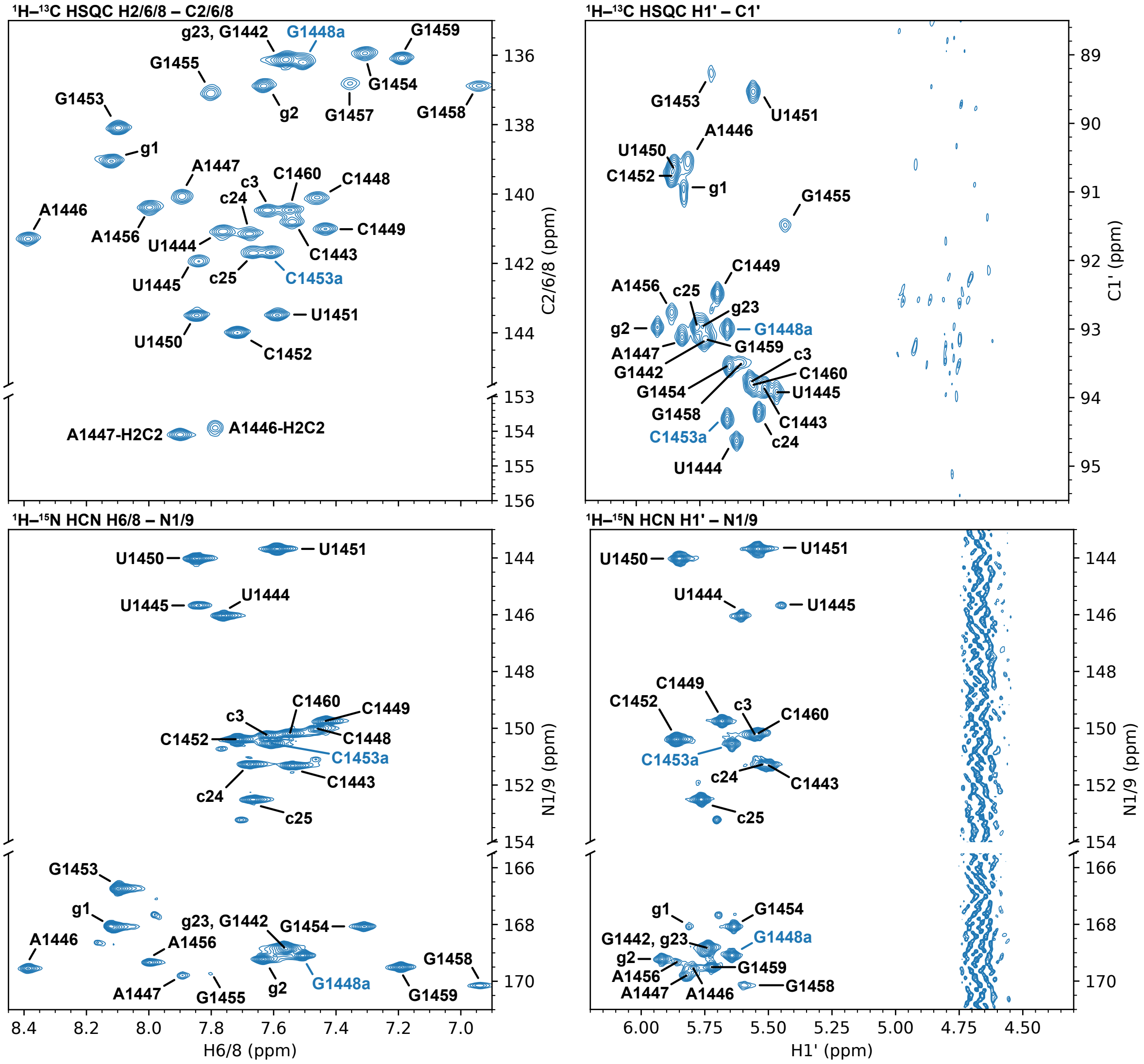


**Figure S5. Chemical shift assignments of the 27 nt h44-top$^{ES}$ hairpin construct.** Conditions: 15 mM NaP, 25 mM NaCl, 0.5 mM EDTA, pH 6.5, 90% $H_2O$/10% $D_2O$, 298 K. All spectra were acquired on a $^{13}C$,$^{15}N$-labeled sample. Sequential imino–imino and anomeric–aromatic correlations are indicated by dashed lines in the different regions of the NOESY spectrum.

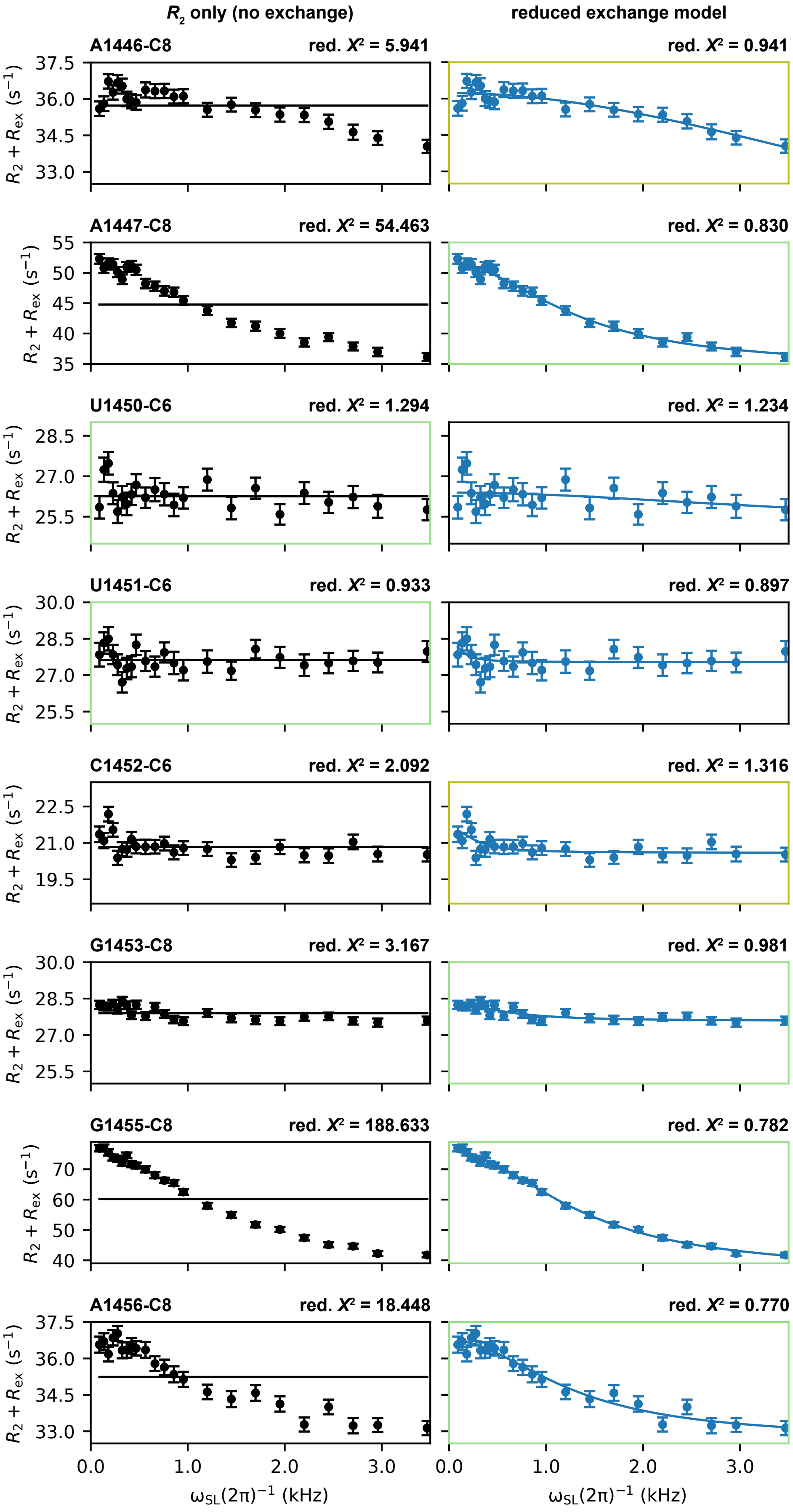

$R_2$ only (no exchange)
reduced exchange model
A1446-C8
red. $X^2$ = 5.941
red. $X^2$ = 0.941
A1447-C8
red. $X^2$ = 54.463
red. $X^2$ = 0.830
U1450-C6
red. $X^2$ = 1.294
red. $X^2$ = 1.234
U1451-C6
red. $X^2$ = 0.933
red. $X^2$ = 0.897
C1452-C6
red. $X^2$ = 2.092
red. $X^2$ = 1.316
G1453-C8
red. $X^2$ = 3.167
red. $X^2$ = 0.981
G1455-C8
red. $X^2$ = 188.633
red. $X^2$ = 0.782
A1456-C8
red. $X^2$ = 18.448
red. $X^2$ = 0.770
$R_2 + R_{ex}$ ($s^{-1}$)
$\omega_{SL}(2\pi)^{-1}$ (kHz)

**Figure S6. On-resonance $^{13}$C relaxation dispersion profiles for h44-top$^{ES}$.** No-exchange ($R_2$ only, left column) and reduced exchange (right column) models were tested against each other; the preferred model for each atom based on statistical criteria is shown with a shaded border. A yellow border indicates clear evidence for residual exchange, albeit with high parameter uncertainties. Parameters for each dataset are given in **Table S5**. Source data are found in **Data S4**.

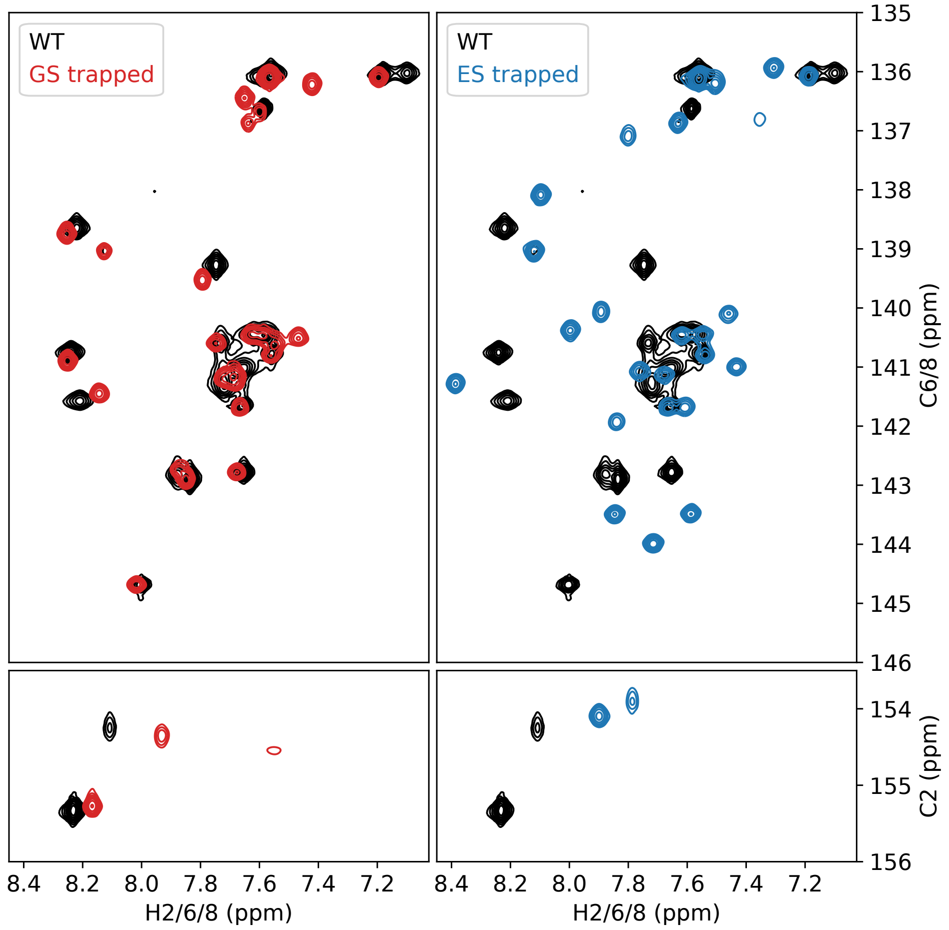


**Figure S7. Overlay of $^{1}$H–$^{13}$C HSQC spectra.** Comparison of the $^{1}$H–$^{13}$C HSQC spectrum in the nucleobase aromatic region for h44-top$^{WT}$ (black) with the h44-top$^{GS}$ mutant (red, left) shows closely matched chemical shift fingerprints in the aromatic region that indicate high structural homology. In contrast, the h44-top$^{ES}$ mutant (blue, right) features chemical shift changes for nearly all nucleobases, indicating the expected large structural rearrangement.

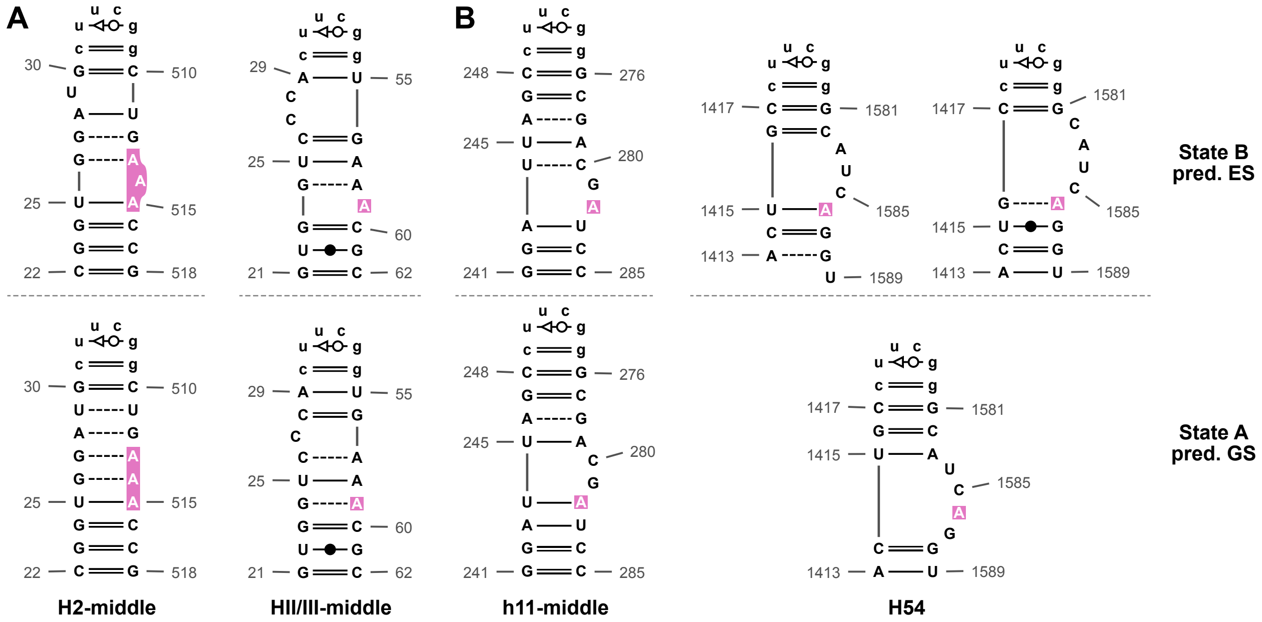


**Figure S8. Minimum-free-energy and alternative secondary structures of rRNA model hairpins undergoing adenosine switching.** (**A**) Hairpins derived from 23S rRNA H2 and 5S rRNA HII/III. Adenosine switching is accompanied by an expansion/contraction of a bulge elsewhere along the stem. (**B**) Hairpins derived from 16S rRNA h11 and 23S rRNA H54. Alternative secondary structures display a bulge shift across the adenosine rather than any of the prototypical switching mechanisms considered in this work. One low-energy alternative structure of the H54 construct displays single-nucleotide end-fraying. Secondary structures of hairpins were predicted with MC-Flashfold; lower-case nucleotides denote artificially added connecting loops that are not included in the sequence numbering; adenosines involved in A-minor interactions with neighboring parts of rRNA are highlighted in pink.

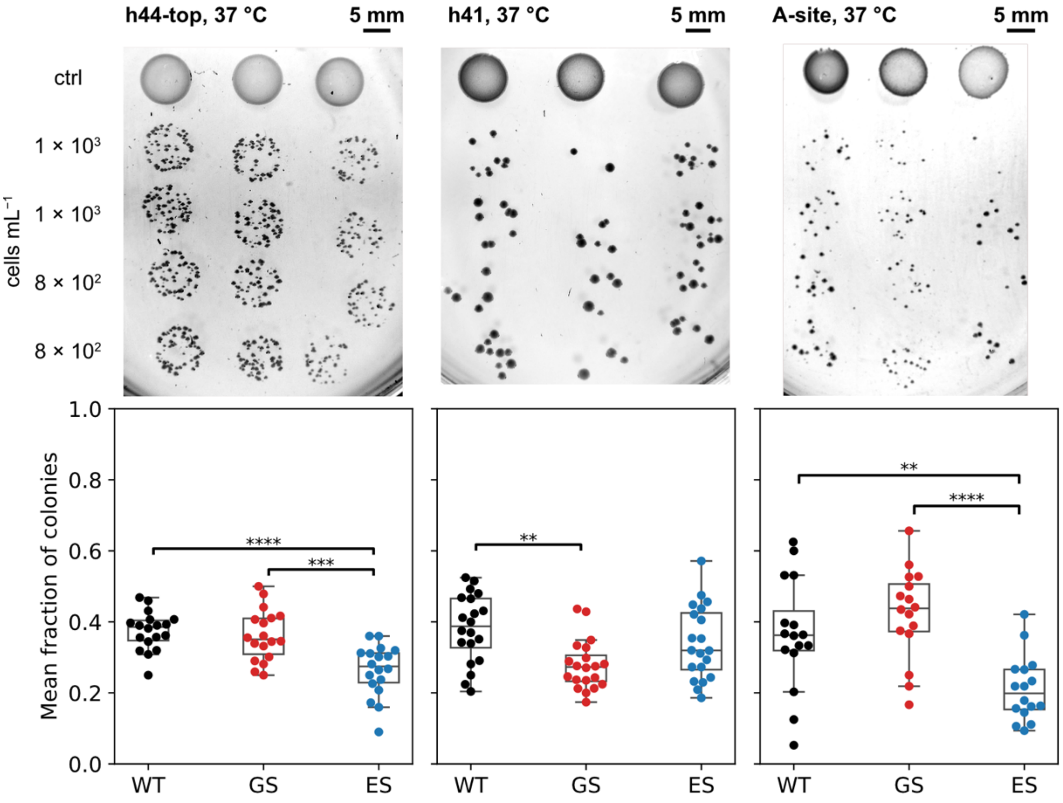


**Figure S9. Spot plating results of *E. coli* SQ171fg.** rRNA mutations in h44-top, h41 and the A-site relative to WT at 37 °C. As discussed in **Supplemental Note 3.3**, both A-site mutant strains contain a background of WT ribosomes. Statistical analysis: one-way ANOVA with Bonferroni correction for multiple means comparisons. ** $P < 0.01$, *** $P < 0.001$, **** $P < 0.0001$. Photographs show individual representative culture plates for each system; scale bars indicate 5 mm length. Source data are found in **Data S1**.

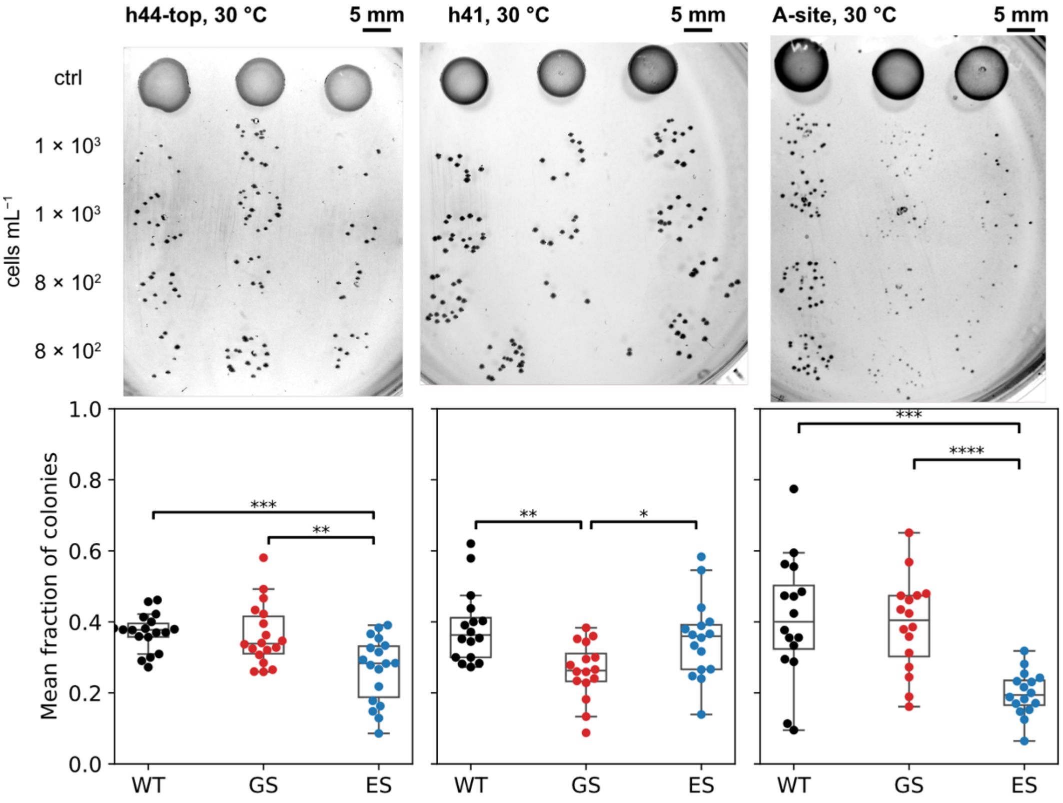


**Figure S10. Spot plating results of *E. coli* SQ171fg.** rRNA mutations in h44-top, h41 and the A-site relative to WT at 30 °C. As discussed in **Supplemental Note 3.3**, both A-site mutant strains contain a background of WT ribosomes. Statistical analysis: one-way ANOVA with Bonferroni correction for multiple means comparisons. * $P < 0.05$, ** $P < 0.01$, *** $P < 0.001$, **** $P < 0.0001$. Photographs show individual representative culture plates for each system; scale bars indicate 5 mm length. Source data are found in **Data S1**.

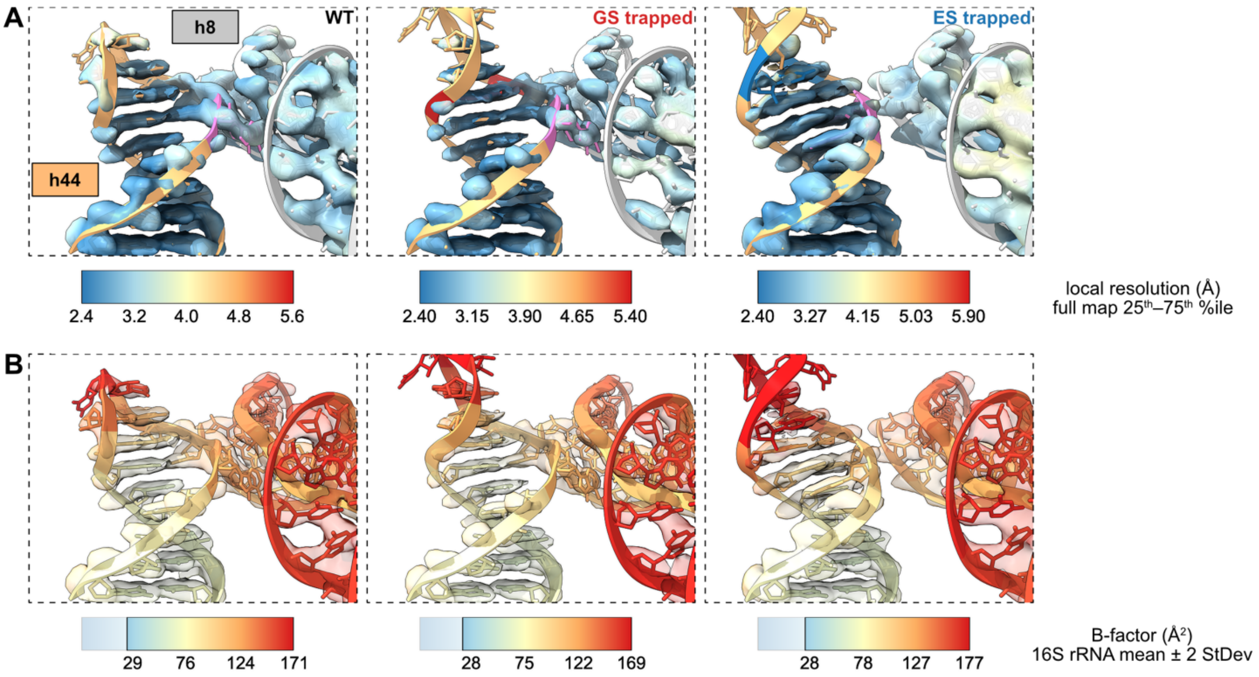


**Figure S11. Additional representations of rRNA mutation sites in cryo-EM models.** Locally filtered maps colored by local resolution (**A**); the same maps and models colored by per-residue B-factors (**B**) for the h44-top region in WT and trapped ribosomes. Color coding in panel (A) is the same as in main text **Figure 4C**. The color scale for local resolution spans the interquartile range of each whole map to facilitate comparison between the different samples. B-factor color scales span the mean ± 2 standard deviations for the 16S rRNA chain in each model (lower end truncated to min. value). Local resolution and B-factors are similar across samples, except for h8 in the h44-top$^{ES}$ ribosome, where a reduced local resolution indicates increased structural flexibility.

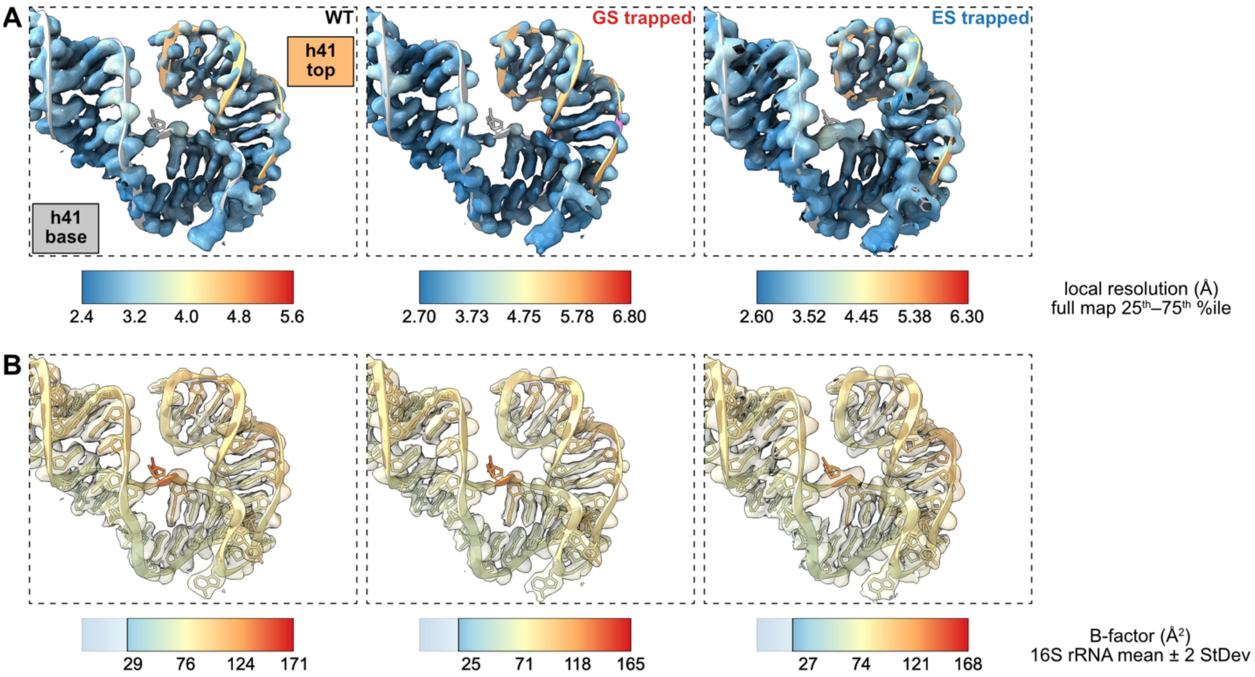


**Figure S12. Additional representations of rRNA mutation sites in cryo-EM models.** Locally filtered maps colored by local resolution (**A**); the same maps and models colored by per-residue B-factors (**B**) for the h41 region in WT and trapped ribosomes. Color coding in panel (A) is the same as in main text **Figure 4D**. The color scale for local resolution spans the interquartile range of each whole map to facilitate comparison between the different samples. B-factor color scales span the mean ± 2 standard deviations for the 16S rRNA chain in each model (lower end truncated to min. value). Local resolution and B-factors are similar across samples.

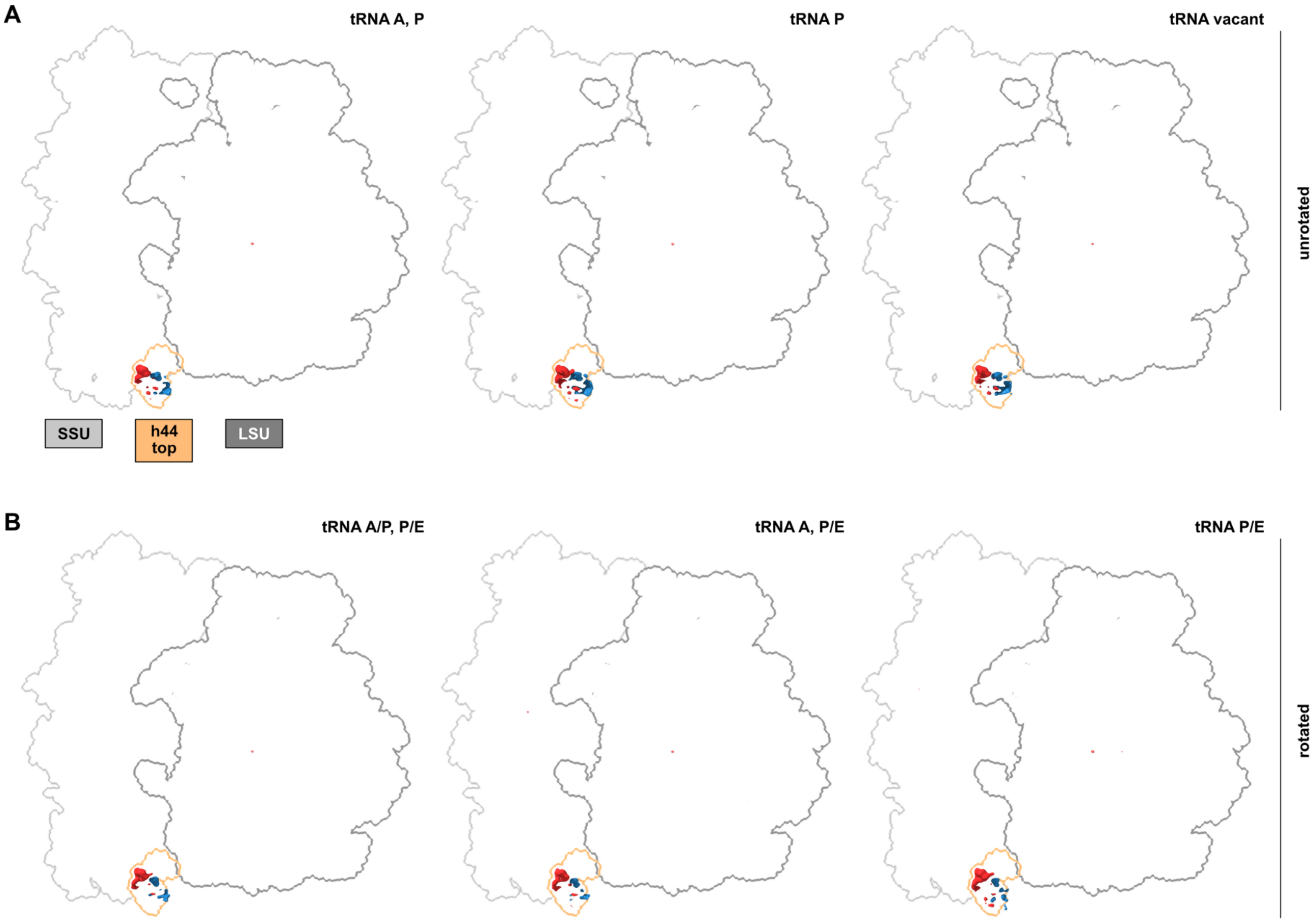
A
tRNA A, P
tRNA P
tRNA vacant
unrotated
SSU
h44 top
LSU
B
tRNA A/P, P/E
tRNA A, P/E
tRNA P/E
rotated

1

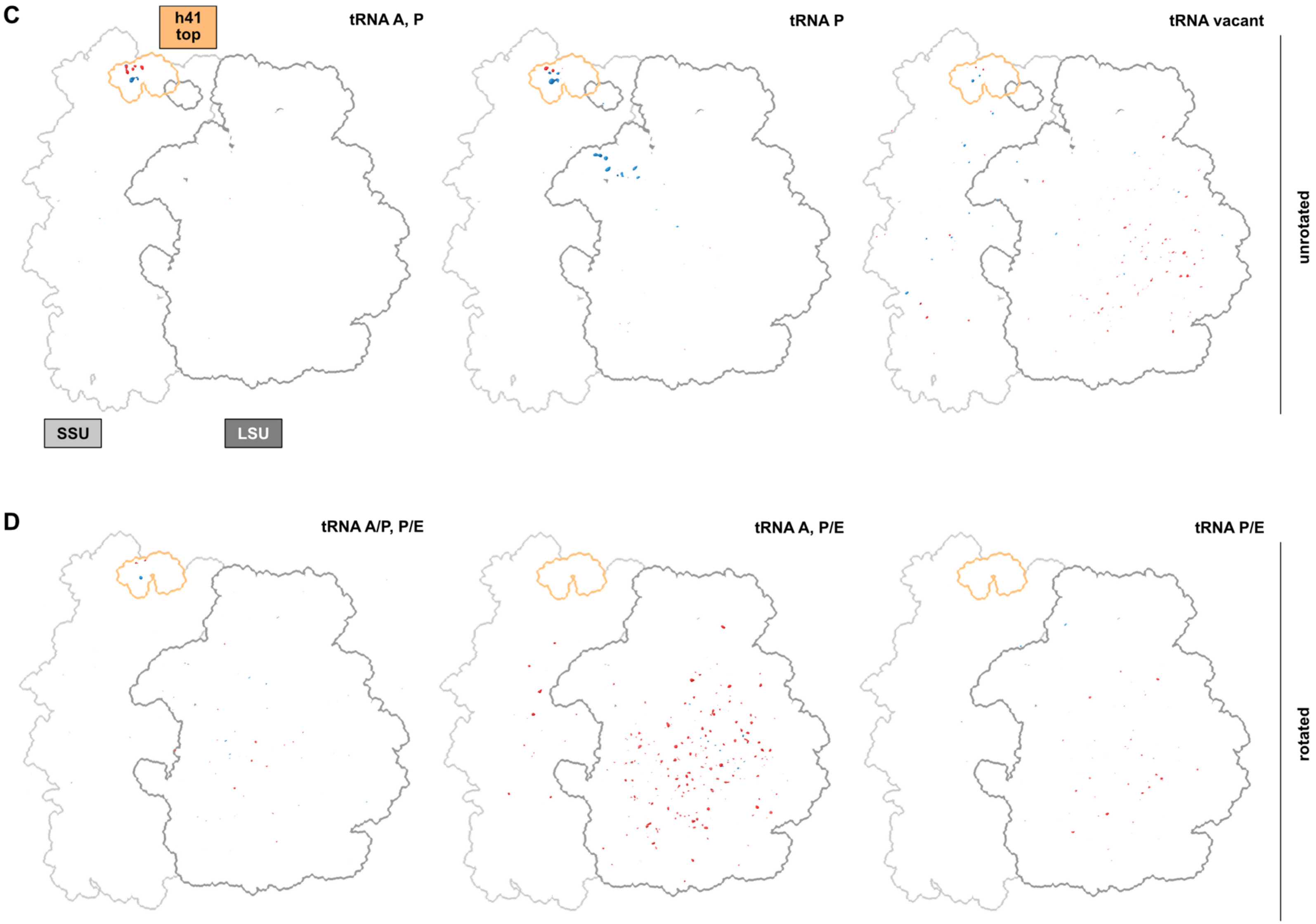
C
h41 top
tRNA A, P
tRNA P
tRNA vacant
unrotated
SSU
LSU
D
tRNA A/P, P/E
tRNA A, P/E
tRNA P/E
rotated

**Figure S13. GS–ES difference maps of mutant ribosomes in different states.** Maps of GS and ES mutant ribosomes in a given subunit rotation and tRNA occupation state were aligned with each other, followed by subtraction of GS−ES with least-squares minimization. Positive and negative contours of the difference map are shown at the same level: red = excess density in the GS mutant, blue = excess density in the ES mutant. The small and large subunits (of the GS model, unrotated with tRNA A, P or rotated with tRNA A/P, P/E) are shown with light and dark grey outlines, respectively. (**A**, **B**) h44-top mutant ribosomes with h44-top region (16S rRNA nt 1442–1460) outlined in orange. The only significant difference is found in the mutation site and associated extra-/intrahelical orientation of A1446/1447. (**C**, **D**) h41 mutant ribosomes with the apical loop of h41 (16S rRNA nt 1258–1277) outlined in orange. Density differences are minimal and arise mainly from P-site tRNA (C, middle) or metal ions and water (C, right; D) due to different map resolutions.

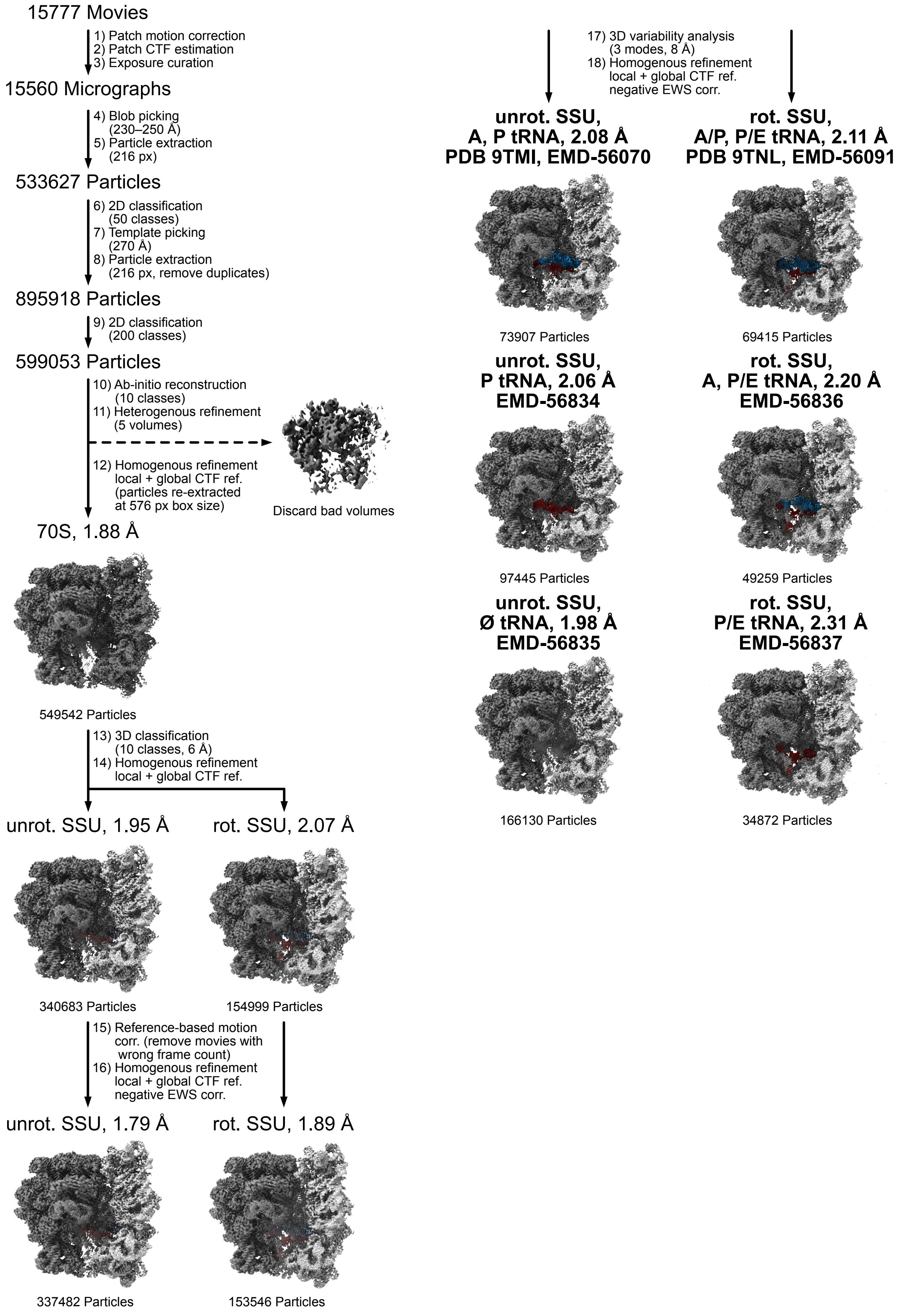


**Figure S14. Processing scheme for cryo-EM dataset of *E. coli* WT ribosomes.**

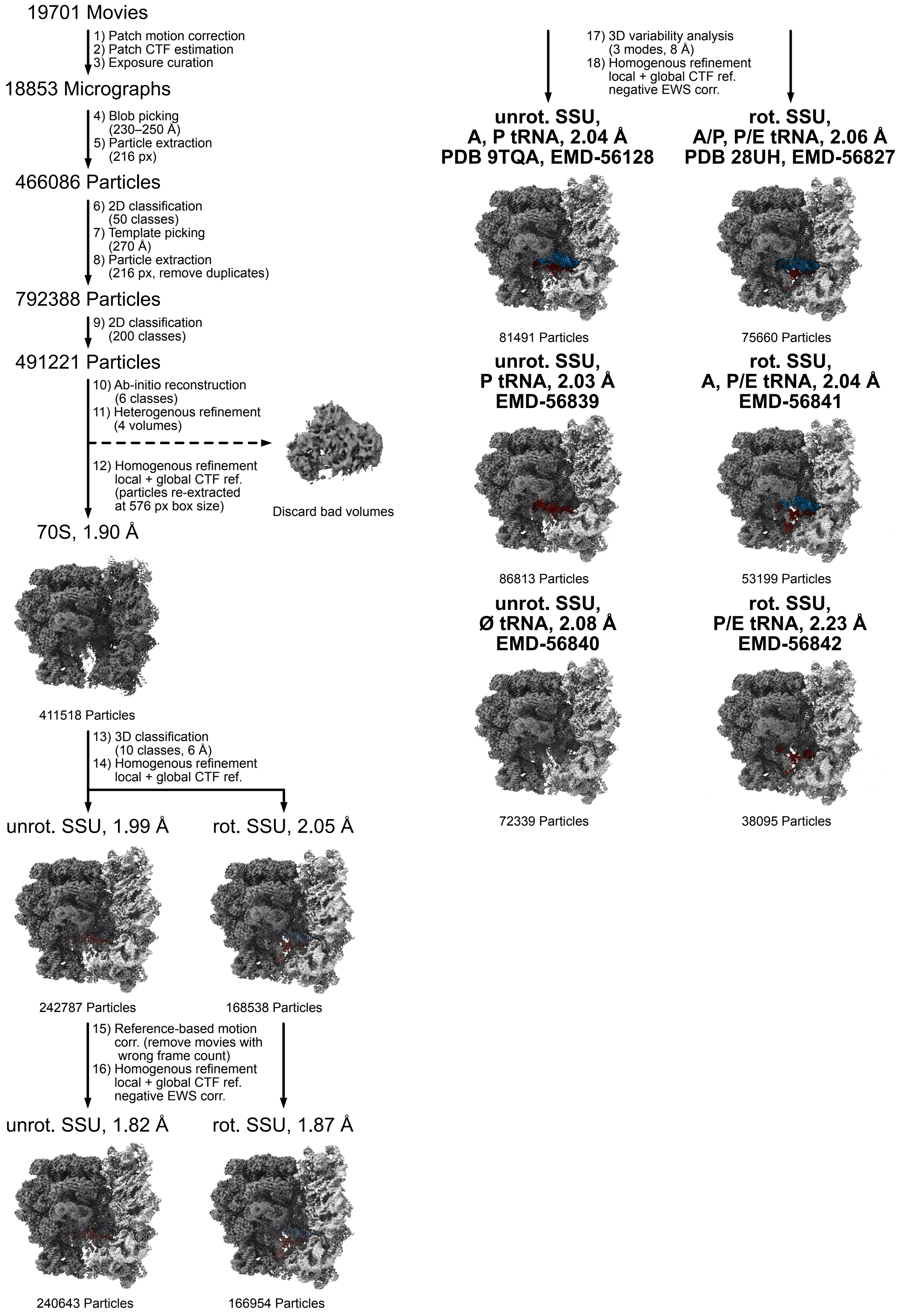


**Figure S15. Processing scheme for cryo-EM dataset of *E. coli* h44-top$^{GS}$ mutant ribosomes.**

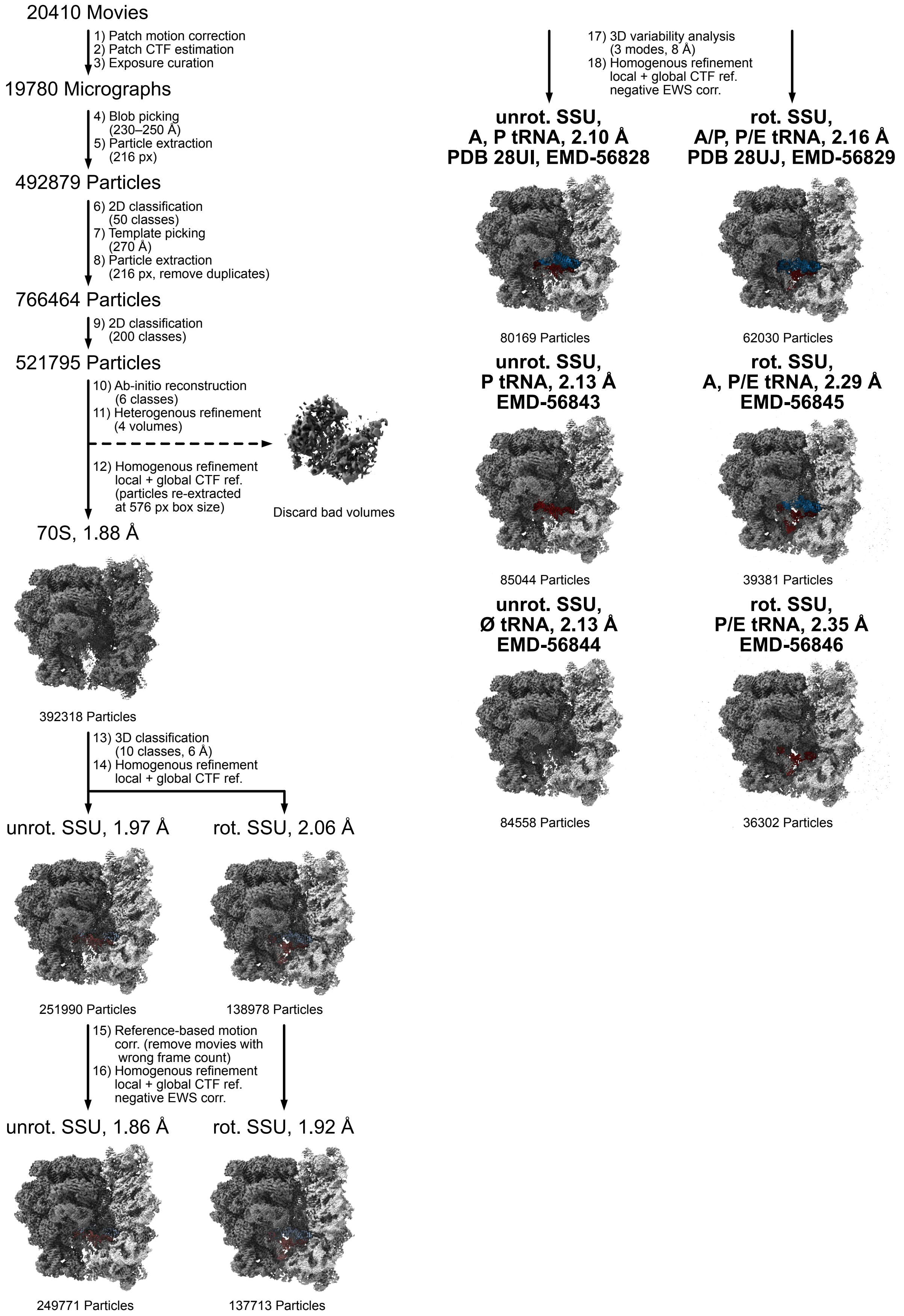


**Figure S16. Processing scheme for cryo-EM dataset of *E. coli* h44-top$^{ES}$ mutant ribosomes.**

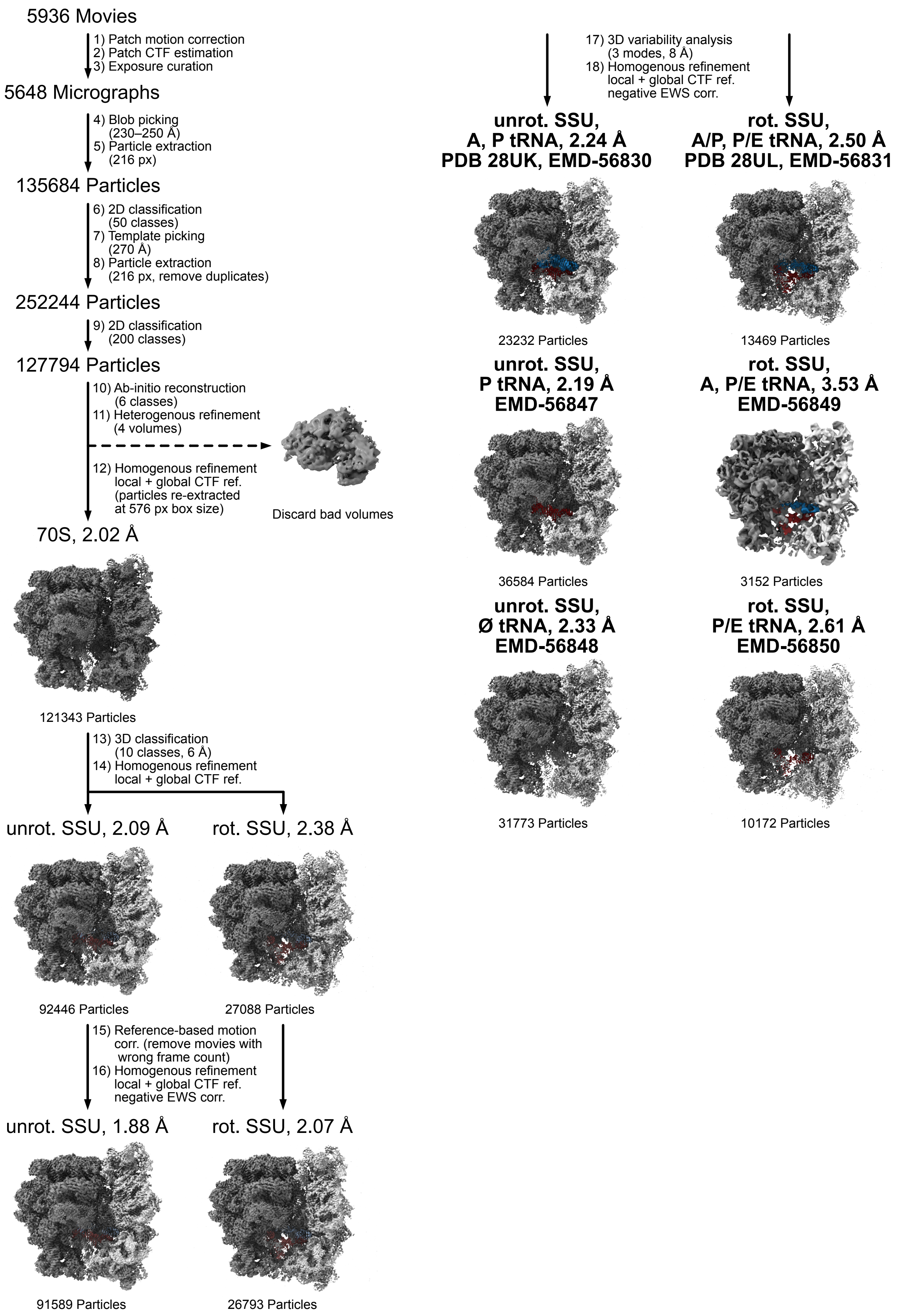


**Figure S17. Processing scheme for cryo-EM dataset of *E. coli* h41$^{GS}$ mutant ribosomes.**

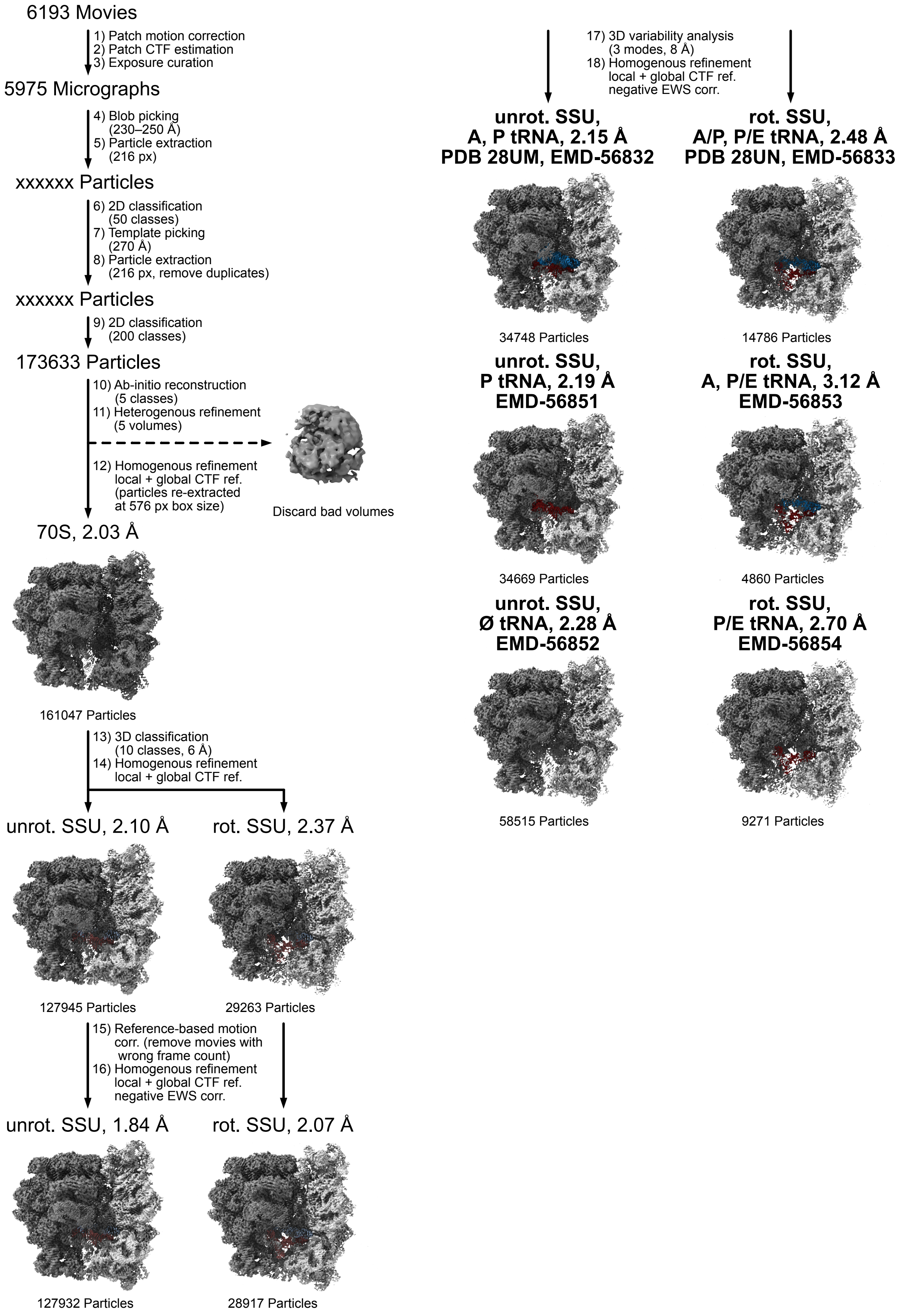


**Figure S18. Processing scheme for cryo-EM dataset of *E. coli* h41$^{ES}$ mutant ribosomes.**

### 2. Supplemental Tables

**Table S2. Fitted exchange parameters for h44-top$^{WT}$.** Cases where atoms from the same nucleotide could be fitted globally with shared exchange rates and populations are shown with grey shading.

| Signal | $R_1$ ($s^{-1}$) | $R_2$ ($s^{-1}$) | $k_{ex,ab}$ ($s^{-1}$) | $p_b$ (%) | $\Delta\omega_{ab}$ $(2\pi)^{-1}$ (Hz) | $k_{ex,ac}$ ($s^{-1}$) | $p_c$ (%) | $\Delta\omega_{ac}$ $(2\pi)^{-1}$ (Hz) | red. $\chi^2$ |
|---|---|---|---|---|---|---|---|---|---|
| U1444-C1' | 2.0 ± 0.06 | 23.2 ± 0.9 | 14631 ± 1297 | 0.9 ± 0.4 | −840 ± 122 | 7.05 ± 1.68 | 31.6 ± 11.3 | −321 ± 80 | 1.212 |
| U1444-N3 | 2.23 ± 0.01 | 6.1 ± 0.2 | | 0.032 ± 0.003 | 1940 ± 104 | – | – | – | |
| U1445-C6 | 1.91 ± 0.08 | 35.2 ± 0.7 | 11340 ± 1224 | 4.5 ± 3.1 | −280 ± 64 | 1916 ± 127 | 2.7 ± 0.3 | −224 ± 17 | 3.170 |
| A1446-H8 | 5.20 ± 0.28 | 23.2 ± 0.5 | | | −277 ± 18 | 46 ± 24 | 13.6 ± 3.1 | 1460 ± 458 | |
| A1446-C1' | 2.21 ± 0.14 | 14.9 ± 1.4 | 14036 ± 456 | 17.4 ± 2.7 | −430 ± 29 | | | −469 ± 24 | 2.875 |
| A1446-C8 | 2.44 ± 0.06 | 28.3 ± 0.3 | | | 128 ± 7 | – | – | – | |
| A1447-C1' | 1.87 ± 0.07 | 21.4 ± 1.2 | 1262 ± 38 | 3.2 ± 0.1 | 359 ± 6 | 16197 ± 2339 | 2.4 ± 1.5 | −416 ± 136 | 2.310 |
| A1447-C2 | 2.67 ± 0.04 | 26.1 ± 0.1 | 1709 ± 202 | 0.9 ± 0.1 | −353.4 | 80 ± 11 | 12.7 ± 1.4 | −229 ± 8 | 1.851 |
| A1447-C8 | 2.48 ± 0.03 | 24.6 ± 0.1 | 1641 ± 131 | 13.0 ± 9.9 | 40 ± 13 | – | – | – | 2.961 |
| C1449-C1' | 2.41 ± 0.06 | 22.1 ± 0.1 | 1259 ± 93 | 5.6 ± 3.0 | −78 ± 15 | – | – | – | 4.249 |
| U1450-C1' | 2.00 ± 0.07 | 19.9 ± 0.2 | 949 ± 20 | 3.0 ± 0.0 | −473 ± 3 | 3761 ± 188 | 0.2 ± 0.0 | −2188 ± 45 | 7.573 |
| U1450-C6 | 2.75 ± 0.08 | 33.9 ± 0.2 | 1374 ± 56 | 2.5 ± 0.1 | 361 ± 7 | – | – | – | 1.443 |
| U1451-C1' | 2.23 ± 0.05 | 18.8 ± 0.1 | 810 ± 75 | 1.7 ± 0.2 | 135 ± 7 | – | – | – | 0.657 |
| U1451-C6 | 2.04 ± 0.07 | 23.7 ± 0.1 | 819 ± 166 | 1.8 ± 1.1 | −210 ± 25 | 916 ± 396 | 1.6 ± 18.1 | 107 ± 81 | 0.690 |
| C1452-C1' | 1.95 ± 0.04 | 22.9 ± 0.1 | 1010 ± 21 | 5.8 ± 0.6 | 238 ± 6 | 28 ± 4 | 41.8 ± 4.5 | −117 ± 21 | 1.120 |
| C1452-C6 | 2.37 ± 0.09 | 32.0 ± 0.1 | | | 126 ± 8 | | | −229 ± 9 | |
| G1453-C1' | 2.33 ± 0.05 | 20.0 ± 0.1 | | | −648 ± 3 | 871 ± 32 | 0.9 ± 0.0 | −390 ± 10 | |
| G1453-C8 | 2.60 ± 0.04 | 24.4 ± 0.1 | 1037 ± 11 | 2.5 ± 0.0 | −801 ± 4 | | | −514 ± 11 | 3.189 |
| G1453-N1 | 1.52 ± 0.02 | 5.86 ± 0.03 | | | 277 ± 2 | – | – | – | |

| | | | | | | | | | |
|---|---|---|---|---|---|---|---|---|---|
| G1453-H1 | 6.96 ± 0.03 | 22.9 ± 0.1 | 1574 ± 67 | 1.6 ± 0.2 | 1535 ± 17 | 367 ± 37 | 3.1 ± 0.2 | 1795 ± 24 | 2.691 |
| G1454-C8 | 2.33 ± 0.07 | 27.9 ± 0.1 | 978 ± 24 | 3.5 ± 0.2 | −315 ± 3.85 | 22 ± 2 | 17.0 ± 3.8 | 149 ± 17 | 0.841 |
| G1454-N1 | 2.08 ± 0.01 | 6.58 ± 0.02 | 1271 ± 65 | 8.2 ± 9.5 | 33 ± 14 | – | – | – | 0.842 |
| G1455-C8 | 2.49 ± 0.09 | 28.9 ± 0.8 | 815 ± 270 | 2.7 ± 1.3 | 207 ± 16 | 8072 ± 4098 | 0.3 ± 5.4 | 478 ± 257 | 0.628 |
| A1456-C1' | 1.00 ± 0.04 | 27.1 ± 0.1 | 5285 ± 433 | 9.9 ± 6.4 | 89 ± 21 | – | – | – | 11.979 |
| A1456-C8 | 2.04 ± 0.09 | 36.4 ± 0.3 | 814 ± 81 | 2.7 ± 0.1 | 205 ± 10 | 2103 ± 447 | 0.3 ± 0.1 | 620 ± 69 | 2.642 |
| G1458-N1 | 2.27 ± 0.01 | 7.17 ± 0.07 | 3332 ± 1924 | 0.0 ± 0.8 | 1218 ± 212 | 1937 ± 685 | 0.1 ± 7.8 | 165 ± 126 | 1.100 |
| Signal | $R_1$ ($s^{-1}$) | $R_2$ ($s^{-1}$) | $k_{ex,ab}$ ($s^{-1}$) | $\phi_{ab}$ $(2\pi)^{-2}$ ($Hz^2$) | | | | | red. $\chi^2$ |
| G1459-N1 | 2.12 ± 0.01 | 6.72 ± 0.01 | 809 ± 229 | 9.0 ± 12.3 | | | | | 1.825 |

**Table S3. Global fit of the upper loop in h44-top$^{WT}$.** A common exchange rate $k_{ex,ab}$ was fitted to all indicated atoms; for C1452 (grey shading), $p_b$ had to be fitted separately. Additional minor exchange processes were fitted individually per atom or with shared parameters between different atoms in the same nucleotide as indicated. Various combinations of minor exchange processes were tested to arrive at the best fit shown here.

| Signal | $R_1$ ($s^{-1}$) | $R_2$ ($s^{-1}$) | $k_{ex,ab}$ ($s^{-1}$) | $p_b$ (%) | $\Delta\omega_{ab}$ $(2\pi)^{-1}$ (Hz) | $k_{ex,ac}$ ($s^{-1}$) | $p_c$ (%) | $\Delta\omega_{ac}$ $(2\pi)^{-1}$ (Hz) | red. $\chi^2$ |
|---|---|---|---|---|---|---|---|---|---|
| C1449-C1' | 2.39 ± 0.05 | 22.2 ± 0.1 | | | −115 ± 2 | – | – | – | |
| U1450-C1' | 2.11 ± 0.07 | 19.6 ± 0.2 | | | −480 ± 3 | 4202 ± 226 | 0.16 ± 0.01 | −2069 ± 44 | |
| U1450-C6 | 2.66 ± 0.09 | 34.6 ± 0.1 | | | 361 ± 4 | – | – | – | |
| U1451-C1' | 2.28 ± 0.05 | 18.6 ± 0.1 | | | 102 ± 2 | – | – | – | |
| U1451-C6 | 2.09 ± 0.08 | 23.6 ± 0.1 | | | −169 ± 3 | 119 ± 215 | 0.8 ± 4.6 | 238 ± 56 | |
| G1453-C1' | 2.33 ± 0.05 | 20.1 ± 0.1 | 1025 ± 8 | 2.74 ± 0.02 | −638 ± 3 | 763 ± 32 | 0.76 ± 0.03 | −343 ± 10 | 2.279 |
| G1453-C8 | 2.61 ± 0.04 | 24.4 ± 0.1 | | | −792 ± 3 | | | −461 ± 13 | |
| G1453-N1 | 1.55 ± 0.02 | 5.85 ± 0.03 | | | 264 ± 1 | – | – | – | |
| G1454-C8 | 2.41 ± 0.07 | 27.7 ± 0.1 | | | −329 ± 2 | 18 ± 20 | 1.2 ± 0.9 | 235 ± 383 | |
| G1454-N1 | 2.08 ± 0.01 | 6.62 ± 0.01 | | | 56 ± 1 | – | – | – | |
| G1455-C8 | 2.52 ± 0.08 | 28.7 ± 1.0 | | | 204 ± 4 | 13119 ± 11825 | 0.1 ± 0.2 | 809 ± 243 | |
| C1452-C1' | 1.96 ± 0.05 | 22.9 ± 0.1 | | 6.0 ± 0.5 | 237 ± 5 | 28 ± 3 | 42 ± 4 | −120 ± 19 | |
| C1452-C6 | 2.37 ± 0.09 | 32.0 ± 0.1 | | | 125 ± 7 | | | −230 ± 9 | |

**Table S4. Fitted exchange parameters for h44-top$^{GS}$.** No-exchange ($R_2$ only) and reduced exchange models were tested against each other; the preferred model for each atom based on statistical criteria is shown with a shaded background (green). Clear evidence for residual exchange is only found for A1447-C8, albeit with high parameter uncertainties (yellow shading).

| Signal | $R_2$ only ($s^{-1}$) | red. $\chi^2$ | $R_2$ ($s^{-1}$) | $k_{ex,ab}$ ($s^{-1}$) | $\phi_{ab}$ $(2\pi)^{-2}$ ($Hz^2$) | red. $\chi^2$ |
|---|---|---|---|---|---|---|
| A1446-C8 | 29.9 ± 0.1 | 0.743 | 27.5 ± 4.9 | 34862 ± 30905 | 2288 ± 16969 | 0.588 |
| A1447-C8 | 25.5 ± 0.1 | 2.155 | 23.0 ± 10.9 | 24697 ± 36592 | 1747 ± 27698 | 0.386 |
| U1451-C6 | 25.8 ± 0.2 | 0.553 | 25.7 ± 2.6 | 680 ± 9208 | 21 ± 5882 | 0.512 |
| G1453-C8 | 26.2 ± 0.1 | 0.944 | 26.2 ± 3.2 | 98 ± 17982 | 2 ± 12006 | 1.039 |

**Table S5. Fitted exchange parameters for h44-top$^{ES}$.** No-exchange (R2 only) and reduced exchange models were tested against each other; the preferred model for each atom based on statistical criteria is shown with a shaded background (green). Yellow shading indicates clear evidence for residual exchange, albeit with high parameter uncertainties.

| Signal | $R_2$ only ($s^{-1}$) | red. $\chi^2$ | $R_2$ ($s^{-1}$) | $k_{ex,ab}$ ($s^{-1}$) | $\phi_{ab}$ $(2\pi)^{-2}$ ($Hz^2$) | red. $\chi^2$ |
|---|---|---|---|---|---|---|
| A1446-C8 | 35.7 ± 0.1 | 5.941 | 26.6 ± 13.5 | 39665 ± 25551 | 9654 ± 32315 | 0.941 |
| A1447-C8 | 44.8 ± 0.2 | 54.463 | 34.5 ± 0.7 | 8202 ± 677 | 3543 ± 398 | 0.830 |
| U1450-C6 | 26.3 ± 0.1 | 1.294 | 25.1 ± 9.8 | 24616 ± 48435 | 783 ± 32777 | 1.234 |
| U1451-C6 | 27.6 ± 0.1 | 0.933 | 27.5 ± 0.2 | 801 ± 5586 | 19 ± 454 | 0.897 |
| C1452-C6 | 20.8 ± 0.1 | 2.092 | 20.6 ± 0.1 | 1707 ± 1951 | 44 ± 120 | 1.316 |
| G1453-C8 | 27.9 ± 0.1 | 3.167 | 27.6 ± 0.1 | 3845 ± 1412 | 69 ± 33 | 0.981 |
| G1455-C8 | 60.2 ± 0.2 | 188.633 | 36.3 ± 0.9 | 8541 ± 373 | 8574 ± 509 | 0.782 |
| A1456-C8 | 35.2 ± 0.1 | 18.448 | 32.7 ± 0.3 | 7923 ± 1135 | 816 ± 165 | 0.770 |

**Table S6. Compiled examples of A-minor interactions spanning the four different kinds of 3D arrangements.** This table is reproduced from ref. [1], where the adenosine residue in the type III motif from PDB 2J00 was mislabeled. The corrected residue number is A1332.

##### A-minor type I

| PDB ID | Chain & residue | | |
|---|---|---|---|
| 1FFK | 0:A521 | 0:G1364 | 0:C637 |
| 1F27 | B:A23 | A:C4 | A:G19 |
| 1VQO | 0:A1007 | 0:A2311 | 0:U2297 |
| 1NJP | 0:A2413 | 0:U2222 | 0:A2060 |

##### A-minor type II

| PDB ID | Chain & residue | | |
|---|---|---|---|
| 1FFK | 0:A520 | 0:G1363 | 0:C638 |
| 1GID | A:A183 | A:C211 | A:G110 |
| 1JJ2 | 0:A1930 | 0:A2266 | 0:U2127 |
| 1VQO | 0:A1458 | 0:U862 | 0:A784 |

##### A-minor type III

| PDB ID | Chain & residue | | |
|---|---|---|---|
| 2J00 | A:A1332 | A:G947 | A:C1234 |
| 1FFK | 0:A519 | 0:A639 | 0:U1362 |

##### A-minor type 0

| PDB ID | Chain & residue | | |
|---|---|---|---|
| 2A64 | A:A287 | A:G272 | A:C260 |
| 1VQO | 0:A1476 | 0:C1862 | 0:G1867 |
| 1FFK | 9:A104 | 0:A957 | 0:U1009 |
| 1K8A | A:A1485 | A:A861 | A:U785 |

### 3. Supplemental Notes

#### 3.1. Residual conformational dynamics in h44-top$^{GS}$ and h44-top$^{ES}$ mutants

No residual dynamics were observed in the loop region of h44-top$^{GS}$, while the A-rich bulge still displayed an expected minor fast conformational exchange process (**Figure S4**, **Table S4**). This is consistent with the exchange model proposed for h44-top$^{WT}$ (main concerted ES for loop rearrangement, additional fast ES in the loop due to intra-/extra-helical adenosine flip) and shows that the inserted base pair effectively prevents the global loop switch but preserves local contributions from individual bulge nucleotides undergoing fast exchange. Similarly, h44-top$^{ES}$ still shows significant relaxation dispersion in the nucleotides previously constituting the bulge, notably at A1447-C8 and G1455-C8 (**Figure S6**, **Table S5**), which can be ascribed to the small free energy contribution of the A:A/G:A tandem mismatch to helix stability.[2]

#### 3.2. A-minor dynamics prediction by secondary structure ensemble analysis

##### *3.2.1. A-minor motif search*

Studies on potential examples for dynamics in A-minor motifs were done on a recent cryo-EM model of the *E. coli* ribosome with a resolution of 2.0 Å (PDB 7K00). Candidate motifs were extracted from the model using a geometric search with WebFR3D.[3] A suitable set of 14 example motifs covering type I, II, III and 0 A-minor interactions was reported by Schlick and coworkers.[1] These base triples are reproduced in **Table S6**, correcting a minor mistake in the description of the type III motif in PDB 2J00. The ribosome model was queried for each motif with a discrepancy cutoff value of 0.5. Base triples in which a nucleotide other than A interacts with the minor groove of a base pair (G-minor etc.) were removed from the output. In case the same base triple matched more than one type of example motif, the higher-discrepancy entries were removed, resulting in a total of 502 individual A-minor interactions. Redundant interactions, i.e., those in which the same adenosine interacts with different neighboring base pairs, were compared against each other and the base triple with the lowest overall discrepancy value was retained. There were 226 redundant and 276 unique interactions, two of which were discarded because they involve ribosome-bound tRNAs.

Because the dynamics under consideration involve an intra-to-extrahelical flipping of adenosine residues, those located within hairpin loops were not taken under further consideration. A list of such hairpin loop segments was obtained from the annotated entry for PDB 7K00 in the URSDB via its web interface.[4] Of the remaining 206 base triples, 71 were located in the 16S rRNA, 133 in the 23S rRNA and two in the 5S rRNA.

##### *3.2.2. Automated construct design*

For building short hairpin constructs, these base triples were categorized either as isolated (two or more nucleotides separation between A-minor adenosines, 118 instances) or as cumulated (not more than one nucleotide separation, 88 instances) motifs and one construct was generated for each such motif. Constructs were designed according to the following principle: 1) 5' and 3' of the A-minor motif, add the appropriate neighboring nucleotides from the same strand of the rRNA sequence until either side has been extended to include three base pairs. A list of categorized base pairs was obtained from WebFR3D. In case of ambiguity, priority was given to WW over other kinds of base pairs. Inter-subunit and cSS/tSS/ncSS/ntSS base pairs were filtered out as they would not provide meaningful interactions during secondary structure prediction. 2) Fill in all nucleotides for the opposite strand. At this stage, when the length of the two strands was different by more than a factor of two (indicating the presence of a large, unstructured bulge), the construct was discarded. 3) If adjacent ends of the two strands are separated by not more than seven nucleotides, connect them with the corresponding native loop found in the rRNA sequence. Otherwise, add a cUUCGg tetraloop. In certain edge cases (e.g., when the helix surrounding the A-minor motif is not strictly antiparallel), this algorithm would produce invalid constructs that were discarded.

##### *3.2.3. Secondary structure analysis*

The sequence of each construct was used for prediction of secondary structure ensembles with MC-Flashfold using an energy threshold value of 8.0 kcal $mol^{-1}$.[5] High-energy folding states that differ from lower-energy ones simply by the symmetric opening or closing of internal loops were removed as they do not represent meaningful structural dynamics. Similarly, structures containing rare and unusual single-base-pair bridges within larger bulge loops were removed. Next, the secondary structure ensemble was partitioned according to the A-minor adenosine being either paired or unpaired; for cumulated motifs with n adenosines, all 2n tuples were considered. The resulting folding states were inspected for the indications of relevant dynamics.

Two different scenarios of intra-to-extrahelical dynamics can be distinguished: 1) Movement or size change of an existing bulge loop along a helical region, during which the A-minor adenosine changes its orientation. This situation mirrors what has been reported for A1492 and A1493 located in the ribosomal A-site.[6] 2) Register shift of a helical region that is accompanied by a size change of the adjacent stem loop, which is what is being reported in this work for A1446 and A1447 in the apical region of h44. For constructs containing an artificially inserted loop, only the first scenario was considered meaningful because such a loop could introduce spurious structural perturbations that are not present in the actual rRNA helix. Among 52 constructs, four new ones were identified that met those conditions and did not exhibit formation of frayed ends upon undergoing intra-to-extrahelical transitions. Their locations in the rRNA and sequences are given in **Table S1**. Two further constructs could undergo said adenosine transition but did not conform to either of the two scenarios outlined above, while 33 constructs with potential secondary structure switches in the designated energy range were prone to more pronounced fraying or other indicators of missing secondary structure context.

Like with the ribosomal A-site and the apical region of h44, mutations were introduced into the four new constructs with the aim of trapping the interconversion of folding states. To that end, two different sets of one or two nucleotides were substituted or deleted, which would either increase the amount of canonical base pairs in the A-paired state and decrease it in the A-bulged state or vice versa. New secondary structures for the mutated constructs were again generated with MC-Flashfold and processed as described above.

The lower-energy state was considered trapped when its energy difference to the higher energy state was increased by at least 2.0 kcal $mol^{-1}$ compared with the unmutated construct. Likewise, the higher-energy state was considered trapped when the sign of the energy difference was inverted, and the magnitude of the difference was at least 2.0 kcal $mol^{-1}$. For all four constructs, trapping of the lower-energy state was successful according to these metrics. For two constructs, the higher-energy states were successfully trapped as well, while for two others the energy difference was less than 2.0 kcal $mol^{-1}$ after state inversion. The respective sequences and secondary structures are given in **Table S1**.

#### 3.3. Viability of *E. coli* strains with mutations to affect A-minor dynamics

During the preparation of *E. coli* strains harboring known or predicted point mutations that would affect A-minor GS–ES conformational equilibria, plasmids were checked by Sanger sequencing in regular intervals.

For the H28 pAM552 mutant plasmid, the expected sequences were present following site-directed mutagenesis. Transformation of the plasmid into *E. coli* SQ171fg with expulsion of the WT *rrnB* operon on the plasmid pCSacB due to sucrose selection was confirmed via PCR, which showed the absence of a band for *sacB*. However, Sanger sequencing failed to confirm the presence of the correct mutations afterwards. In repeated transformations, either WT H28 or a mix of several sequences at the intended mutation site was observed and efforts to generate H28 mutant ribosomes were not pursued further.

The same behavior was found for the h44-A-site mutants. Therefore, an alternative approach was followed in which synthetic 16S mutant sequences were inserted into pAM552 by restriction cloning. After transformation into *E. coli* SQ171fg, Sanger sequencing confirmed the presence of the intended point mutations. These h44-A-site mutant strains were used in spot plating. However, repeated sequencing at later time points showed mixed mutant and WT 16S rRNA populations, suggesting that a previous minor contamination was more viable and overtook the mutants. The spot plating results should therefore be interpreted with the caveat that any observed biological effect would be attenuated due to the coexistence of both WT and mutant ribosomes in the same strain. Due to this heterogeneity, no ribosomes were isolated for cryo-EM analysis.

#### 3.4. Comparison of h44-top$^{GS}$ and h44-top$^{ES}$ structures by NMR and cryo-EM

NMR data suggests good agreement between the secondary structure of the h44-top$^{WT}$ hairpin and the corresponding region of nucleotides 1442–1460 in the 16S rRNA of the *E. coli* 70S ribosome (cf. cryo-EM models such as PDB 7K00). Base pair C1448:G1455 flanking the adenosine-rich bulge is stably formed based on a cross-peak in the $^{1}H$–$^{15}N$ HSQC spectrum, as is the U1444:G1458 wobble base pair. Imino signals for U1445:G1457 are not observed likely due to rapid exchange with water as this site is adjacent to the internal loop. Sequential ribose–base and base–base correlations in the $^{1}H$–$^{1}H$ NOESY together with nucleobase chemical shifts are consistent with a stacked, intrahelical orientation of A1456, while A1446 is bulged; the latter is further supported by broadening of the A1446-C2H2 resonance due to conformational flexibility. A1447, however, appears to adopt a more helical and less bulged orientation than in the ribosomal environment, as inferred from NOE contacts from its nucleobase H2 and H6 to C1448-H6. This is not entirely unexpected as it lacks the interaction in the NMR construct.

The excellent agreement between chemical shifts for the 25 nt h44-top$^{WT}$ construct and the 27 nt h44-top$^{GS}$ mutant demonstrates that both adopt the same secondary structures, with only local perturbations arising due to insertion of the G1448a:C1454a base pair. The adenosine-rich bulges are in fact superimposable between the WT and trapped GS mutant 70S ribosome cryo-EM models obtained in this study, irrespective of the rotation state of the ribosome. Notably, the A1446-C2H2 resonance in the $^{1}H$–$^{13}C$ HSQC spectrum has sharpened significantly (**Figure S3**) due to trapping of the intermediate timescale conformational exchange process.

Signatures of the alternative secondary structure stabilized in the 27 nt h44-top$^{ES}$ mutant are the disappearance of signals characteristic for a classical UUCG tetraloop fold, chemical shift perturbations throughout the adenosine-rich bulge, and observable imino signals for the U1445:G1457 wobble base pair. Strong NOE cross peaks from A1446-H8 to U1445-H1' and U1445-H6 support a standard N9–C1' *anti* torsion for A1446 in its new, intrahelical stacked orientation (**Figure S5**). The resulting A1446:A1456 tSH / A1447:G1455 tHS tandem mismatch is a common motif in ribosomal RNA[7]. Our cryo-EM data of the respective mutant ribosome confirms this interpretation and allows unambiguous model building of the tandem mismatch. Density in the new UUC triloop region is relatively weak but clearly shows the absence of a UUCG tetraloop fold.

### 4. Supplemental Methods

#### 4.1. General information

##### 4.1.1. Materials

All commercially available reagents were purchased from Sigma-Aldrich in RNase-free quality.

DNA templates for in vitro transcription were purchased from Integrated DNA Technologies (IDT) with purification by standard desalting and used without further purification. Unlabeled and $^{13}C$,$^{15}N$-labeled ribonucleotide triphosphates (NTPs), guanosine monophosphate (GMP) and inorganic pyrophosphatase (IPPase) were purchased from Sigma-Aldrich. RNaseOUT ribonuclease inhibitor was purchased from Thermo Fisher. T7 RNA polymerase was prepared by the Protein Science Facility at Karolinska Institutet (Stockholm, Sweden).

Standard 5'-O-DMT/2'-O-TBDMS-protected 3'-β-cyanoethyl phosphoramidites of $N^6$-benzoyladenosine, $N^4$-acetylcytidine, $N^2$-acetylguanosine and uridine were purchased from Glen Research. Standard 5'-O-DMT/2'-O-Ac-protected CPG support (1000 Å, 20–30 µmol/g) was purchased from Sigma-Aldrich.

*E. coli* strain SQ171fg and plasmid pAM552 in *E. coli* DH5α (addgene #154131) were generously donated by Chandra Sekhara Mandava and Suparna Sanyal (Department of Cell and Molecular Biology, Uppsala University).

##### 4.1.2. Bacterial work

Where applicable, antibiotics were used at a concentration of 100 µg $mL^{-1}$ for ampicillin or 50 µg $mL^{-1}$ for kanamycin and spectinomycin.

#### 4.2. NMR spectroscopy

##### 4.2.1. Instrumentation

NMR experiments were performed on a Bruker Avance III HD 600 spectrometer equipped with a 5 mm QCI-P cryoprobe at a temperature of 298 K unless indicated otherwise

##### 4.2.2. Relaxation dispersion

###### Fast exchange model and reduced model

For systems undergoing fast exchange between two sites a and b, $R_{1\rho}$ is given by ref. [8] as:

$$R_{1\rho} = R_1 \cos^2\theta + R_2 \sin^2\theta + \frac{\phi_{\mathrm{ab}} k_{\mathrm{ex,ab}}}{k_{\mathrm{ex,ab}}^2 + \omega_{\mathrm{eff}}^2} \sin^2\theta \tag{1}$$

in which

$$\phi_{\mathrm{ab}} = p_{\mathrm{a}} p_{\mathrm{b}} \Delta\omega_{\mathrm{ab}}^2 \tag{2}$$

$$\Delta\omega_{\mathrm{ab}} = \omega_{\mathrm{b}} - \omega_{\mathrm{a}} \tag{3}$$

$$k_{\mathrm{ex,ab}} = k_{\mathrm{a}\to\mathrm{b}} + k_{\mathrm{b}\to\mathrm{a}} \tag{4}$$

$$\omega_{\mathrm{eff}}^2 = \omega_1^2 + \Delta\Omega^2 \tag{5}$$

$$\Delta\Omega = \omega_{\mathrm{obs}} - \omega_{\mathrm{rf}} \tag{6}$$

$$\theta = \arctan\frac{\omega_1}{\Delta\Omega} \tag{7}$$

Note that the offset ΔΩ in equation (6) is defined as the resonance frequency of the observed signal in the rotating frame of the spin lock. However, in the relaxation dispersion experiments we adjust the placement of the carrier with respect to the frequency of the observed signal and define $\Delta\Omega_{\mathrm{exp}} = -\Delta\Omega$. The exchange contribution will therefore be maximal when $\Delta\Omega_{\mathrm{exp}} = \Delta\omega_{\mathrm{ab}}$. When plotting data, this is emphasized by the $\Delta\Omega_{\mathrm{exp}}$ axis running from positive to negative, i.e., in the same direction as a conventional chemical shift axis.

It has been shown[9] that even under fast exchange conditions, the sign and magnitude of the chemical shift difference $\Delta\omega_{\mathrm{ab}}$ can be determined to a certain extent and the use of equation (9) was preferred over equation (1).

When testing for the presence of statistically significant exchange contributions compared to an $R_2$-only model without collecting full off-resonance datasets, a reduced version of equation (1) was used:

$$R_{1\rho} = R_2 + \frac{\phi_{\mathrm{ab}} k_{\mathrm{ex,ab}}}{k_{\mathrm{ex,ab}}^2 + \omega_{\mathrm{eff}}^2} \tag{8}$$

#### Two-site exchange model

A highly accurate approximation for $R_{1\rho}$ in the slow-to-intermediate exchange regime has been derived in ref. [10] by applying Laguerre's method to the characteristic polynomial of the 6 × 6 Bloch–McConnell evolution matrix for two-site exchange:

$$R_{1\rho} = R_1 \cos^2\theta + R_2 \sin^2\theta + \frac{p_{\mathrm{a}} p_{\mathrm{b}} \Delta\omega_{\mathrm{ab}}^2 k_{\mathrm{ex,ab}}}{\frac{\omega_{\mathrm{a}}^2 \omega_{\mathrm{b}}^2}{\omega_{\mathrm{eff}}^2} + k_{\mathrm{ex,ab}}^2 - p_{\mathrm{a}} p_{\mathrm{b}} \Delta\omega_{\mathrm{ab}}^2 \sin^2\theta \left(1 + \frac{2k_{\mathrm{ex,ab}}^2 \left(p_{\mathrm{a}}\omega_{\mathrm{a}}^2 + p_{\mathrm{b}}\omega_{\mathrm{b}}^2\right)}{\omega_{\mathrm{a}}^2\omega_{\mathrm{b}}^2 + \omega_{\mathrm{eff}}^2 k_{\mathrm{ex,ab}}^2}\right)} \sin^2\theta \tag{9}$$

In addition to equations (3)–(7), the following definitions are used:

$$\omega_{\mathrm{a}}^2 = \omega_1^2 + \delta_{\mathrm{a}}^2 \tag{10}$$

$$\omega_{\mathrm{b}}^2 = \omega_1^2 + \delta_{\mathrm{b}}^2 \tag{11}$$

$$\delta_{\mathrm{a}} = \Delta\Omega - p_{\mathrm{b}}\Delta\omega_{\mathrm{ab}} \tag{12}$$

$$\delta_{\mathrm{b}} = \Delta\Omega + p_{\mathrm{a}}\Delta\omega_{\mathrm{ab}} \tag{13}$$

#### Three-site exchange model

Palmer and coworkers have derived a general model for n-site exchange without minor exchange using perturbation theory.[11] In analogy to that formalism, it has been proposed in ref. [12] to extend the two-state model in equation (9) by treating the exchange a–b and a–c as independent processes:

$$R_{1\rho} = R_1 \cos^2\theta + R_2 \sin^2\theta + \frac{p_\mathrm{a} p_\mathrm{b} \Delta\omega_\mathrm{ab}^2 k_\mathrm{ex,ab}}{\frac{\omega_\mathrm{a}^2 \omega_\mathrm{b}^2}{\omega_\mathrm{eff}^2} + k_\mathrm{ex,ab}^2 - p_\mathrm{a} p_\mathrm{b} \Delta\omega_\mathrm{ab}^2 \sin^2\theta \left(1 + \frac{2k_\mathrm{ex,ab}^2 \left(p_\mathrm{a}\omega_\mathrm{a}^2 + p_\mathrm{b}\omega_\mathrm{b}^2\right)}{\omega_\mathrm{a}^2\omega_\mathrm{b}^2 + \omega_\mathrm{eff}^2 k_\mathrm{ex,ab}^2}\right)} \sin^2\theta + \frac{p_\mathrm{a} p_\mathrm{c} \Delta\omega_\mathrm{ac}^2 k_\mathrm{ex,ac}}{\frac{\omega_\mathrm{a}^2 \omega_\mathrm{c}^2}{\omega_\mathrm{eff}^2} + k_\mathrm{ex,ac}^2 - p_\mathrm{a} p_\mathrm{c} \Delta\omega_\mathrm{ac}^2 \sin^2\theta \left(1 + \frac{2k_\mathrm{ex,ac}^2 \left(p_\mathrm{a}\omega_\mathrm{a}^2 + p_\mathrm{c}\omega_\mathrm{c}^2\right)}{\omega_\mathrm{a}^2\omega_\mathrm{c}^2 + \omega_\mathrm{eff}^2 k_\mathrm{ex,ac}^2}\right)} \sin^2\theta \tag{14}$$

The definitions given above have been amended as follows:

$$\Delta\omega_\mathrm{ac} = \omega_\mathrm{c} - \omega_\mathrm{a} \tag{15}$$

$$k_\mathrm{ex,ac} = k_{\mathrm{a}\to\mathrm{c}} + k_{\mathrm{c}\to\mathrm{a}} \tag{16}$$

$$\omega_\mathrm{c}^2 = \omega_1^2 + \delta_\mathrm{c}^2 \tag{17}$$

$$\delta_\mathrm{a} = \Delta\Omega - p_\mathrm{b}\Delta\omega_\mathrm{ab} - p_\mathrm{c}\Delta\omega_\mathrm{ac} \tag{18}$$

$$\delta_\mathrm{b} = \Delta\Omega + (1 - p_\mathrm{b})\Delta\omega_\mathrm{ab} - p_\mathrm{c}\Delta\omega_\mathrm{ac} \tag{19}$$

$$\delta_\mathrm{c} = \Delta\Omega - p_\mathrm{b}\Delta\omega_\mathrm{ab} + (1 - p_\mathrm{c})\Delta\omega_\mathrm{ac} \tag{20}$$

Equation (14) is sufficiently accurate when the exchange rates for the two processes are different, and the populations of b and c are small.

### 5. Supplemental References


1. Xin, Y., Laing, C., Leontis, N.B., and Schlick, T. (2008). Annotation of tertiary interactions in RNA structures reveals variations and correlations. RNA *14*, 2465–2477. https://doi.org/10.1261/rna.1249208.

2. Mathews, D.H., Sabina, J., Zuker, M., and Turner, D.H. (1999). Expanded sequence dependence of thermodynamic parameters improves prediction of RNA secondary structure1. J. Mol. Biol. *288*, 911–940. https://doi.org/10.1006/jmbi.1999.2700.

3. Petrov, A.I., Zirbel, C.L., and Leontis, N.B. (2011). WebFR3D--a server for finding, aligning and analyzing recurrent RNA 3D motifs. Nucleic Acids Res. *39*, W50–W55. https://doi.org/10.1093/nar/gkr249.

4. Baulin, E., Yacovlev, V., Khachko, D., Spirin, S., and Roytberg, M. (2016). URS DataBase: universe of RNA structures and their motifs. Database J. Biol. Databases Curation *2016*, baw085. https://doi.org/10.1093/database/baw085.

5. Dallaire, P., and Major, F. (2016). Exploring Alternative RNA Structure Sets Using MC-Flashfold and db2cm. Methods Mol. Biol. Clifton NJ *1490*, 237–251. https://doi.org/10.1007/978-1-4939-6433-8_15.

6. Dethoff, E.A., Petzold, K., Chugh, J., Casiano-Negroni, A., and Al-Hashimi, H.M. (2012). Visualizing transient low-populated structures of RNA. Nature *491*, 724–728. https://doi.org/10.1038/nature11498.

7. Gautheret, D., Konings, D., and Gutell, R.R. (1994). A major family of motifs involving G · A mismatches in ribosomal RNA. J. Mol. Biol. *242*, 1–8. https://doi.org/10.1006/jmbi.1994.1552.

8. Davis, D.G., Perlman, M.E., and London, R.E. (1994). Direct Measurements of the Dissociation-Rate Constant for Inhibitor-Enzyme Complexes via the $T_{1\rho}$ and $T_2$ (CPMG) Methods. J. Magn. Reson. B *104*, 266–275. https://doi.org/10.1006/jmrb.1994.1084.

9. Bothe, J.R., Stein, Z.W., and Al-Hashimi, H.M. (2014). Evaluating the uncertainty in exchange parameters determined from off-resonance R1ρ relaxation dispersion for systems in fast exchange. J. Magn. Reson. *244*, 18–29. https://doi.org/10.1016/j.jmr.2014.04.010.

10. Miloushev, V.Z., and Palmer, A.G. (2005). R1ρ relaxation for two-site chemical exchange: General approximations and some exact solutions. J. Magn. Reson. *177*, 221–227. https://doi.org/10.1016/j.jmr.2005.07.023.

11. Trott, O., and Palmer III, A.G. (2004). Theoretical study of R1ρ rotating-frame and R2 free-precession relaxation in the presence of n-site chemical exchange. J. Magn. Reson. *170*, 104–112. https://doi.org/10.1016/j.jmr.2004.06.005.

12. Kimsey, I.J., Petzold, K., Sathyamoorthy, B., Stein, Z.W., and Al-Hashimi, H.M. (2015). Visualizing transient Watson–Crick-like mispairs in DNA and RNA duplexes. Nature *519*, 315–320. https://doi.org/10.1038/nature14227.
